# Coupling gene expression dynamics to cell size dynamics and cell cycle events: exact and approximate solutions of the extended telegraph model

**DOI:** 10.1101/2022.06.15.496247

**Authors:** Chen Jia, Ramon Grima

## Abstract

The standard model describing the fluctuations of mRNA numbers in single cells is the telegraph model which includes synthesis and degradation of mRNA, and switching of the gene between active and inactive states. While commonly used, this model does not describe how fluctuations are influenced by the cell cycle phase, cellular growth and division, and other crucial aspects of cellular biology. Here we derive the analytical time-dependent solution of an extended telegraph model that explicitly considers the doubling of gene copy numbers upon DNA replication, dependence of the mRNA synthesis rate on cellular volume, gene dosage compensation, partitioning of molecules during cell division, cell-cycle duration variability, and cell-size control strategies. Based on the time-dependent solution, we obtain the analytical distributions of transcript numbers for lineage and population measurements in steady-state growth and also find a linear relation between the Fano factor of mRNA fluctuations and cell volume fluctuations. We show that generally the lineage and population distributions in steady-state growth cannot be accurately approximated by the steady-state solution of extrinsic noise models, i.e. a telegraph model with parameters drawn from probability distributions. This is because the mRNA lifetime is often not small enough compared to the cell cycle duration to erase the memory of division and replication. Accurate approximations are possible when this memory is weak, e.g. for genes with bursty expression and for which there is sufficient gene dosage compensation when replication occurs.

## Introduction

Experiments have revealed a large cell-to-cell variation in the number of mRNA molecules in isogenic populations [1–3]. This can in part be explained by stochastic effects in gene expression due to the low copy numbers of many components, including DNA and important regulatory molecules [4]. Live-cell imaging approaches allow a direct visualization of stochastic bursts of gene expression in living cells [5]. However these experiments are challenging and hence more commonly one measures the mRNA expression per cell from single-molecule fluorescence *in situ* hybridization (smFISH) [5] or single-cell RNA sequencing (scRNA-seq) experiments [6].

The experimental distributions of mRNA numbers are fitted to the predictions of mathematical models, by which one can obtain estimates of the rates of several important transcriptional processes [7–10]. The most common model of this type is the so-called two-state or random telegraph model of gene expression [11, 12]. This is composed of four (effective) reactions

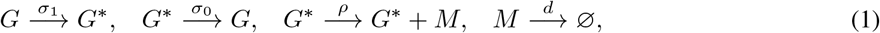

where the first two reactions describe the switching of the gene between an active state *G*\* and an inactive state *G*, the third reaction describes transcription while the gene is in the active state, and the fourth reaction describes the degradation of the mRNA *M*. The chemical master equation (CME) describing the telegraph model can be exactly solved in steady-state, as well as in time [12–15]. Extensions of this model to include more than two gene states have also been considered [16–18].

A substantial number of genes are inactive most of the time and in the brief time that they are active, a large number of mRNA molecules are transcribed but not degraded [19]. This leads to bursty expression. The probability of *r* new mRNA molecules being transcribed before the gene switches off, i.e. a burst of size *r*, is *P*(*r*) = *p^r^* (1–*p*), where *p* = *ρ*/(*ρ* + *σ*_0_) is the probability that the gene synthesizes an mRNA molecule, conditional on it being in the active state [20]. This distribution is geometric with mean *ρ/σ*_0_. The average time between two consecutive bursts is 1/*σ*_0_ + 1/*σ*_1_ ≈ 1/*σ*_1_ since the gene spends most of its time off (*σ*_0_ ≫ *σ*_1_); in other words the rate of burst production is approximately *σ*_1_. It follows that the reaction scheme given in Eq. (1) can be reduced to an effective one-state model composed of only two reactions

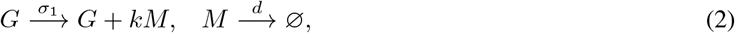

where *k* is the transcriptional burst size which is geometrically distributed with mean *ρ/σ*_0_. The geometric burst size distribution has been validated experimentally [1]. The CME for this model can be solved exactly in steady-state leading to the well-known negative binomial distribution of mRNA numbers [21, 22], which is also widely used in scRNA-seq analysis [23]. Because of the unimodality of this distribution, this simplified model cannot explain bimodality in gene expression [24, 25], a feature that can be explained by the two-state model.

However, the conventional one-state and two-state models are very limited in their predictive power because they lack a description of many cellular processes that are known to have a profound impact on the distribution of mRNA numbers in single cells, e.g. the doubling of gene copy numbers upon DNA replication [26], partitioning of molecules during cell division [27], scaling of the mRNA synthesis rate with cell volume [28–32], and stochasticity in the cell cycle duration and growth rate that is related to cell-size control strategies [33–39]. Recently, numerous efforts have been made to extend the conventional one-state and two-state models to include some description of these processes and yet retain analytical tractability. Some studies focused on the moment statistics (mean and variance) of mRNA and protein numbers [40–44], while other studies additionally obtain the analytical distributions of molecule numbers [22, 45–49]. Please refer to Table 1 for a summary of exactly solvable extensions of the one-state and two-state models that explicitly capture cell birth, growth, and division.

**Table 1:** Exactly solvable gene expression models that explicitly describe cell birth, growth, and division.

| Effective one-state models |  |  |
| --- | --- | --- |
|  | gene replication not considered | gene replication considered |
| volume-independent transcription | Ref. [45] | Refs. [46–48] |
| volume-dependent transcription | Ref. [49] | Refs. [22, 48] |
| Two-state models |  |  |
|  | gene replication not considered | gene replication considered |
| volume-independent transcription | × | × |
| volume-dependent transcription | Ref. [49] | × |

Due to mathematical complexity, most previous work is limited to the effective one-state model with the gene product (mRNA or protein) produced in a constitutive or bursty manner [22,45–48]. Some of these models incorporate the scaling of transcription activity with cell volume [22, 48], while the rest do not. We note that the latter case is not to be seen as unphysical since while the scaling of transcription with volume is commonly observed, it is by no means a universal phenomenon (in both prokaryotic [50, 51] and eukaryotic cells [52–55] there are examples where there is no such scaling). As for the conventional one-state model shown in Eq. (2), the main limitation is the assumption of instantaneous bursts, while in reality there is a finite time for the bursts to occur. A distinct advantage of the extended one-state models over the conventional one is that those which describe gene replication [47] are able to produce bimodal distributions.

The exact solution of extended two-state models that incorporate cell birth, growth, and division has not received much attention. A recent study [49] made progress in this direction. In particular, the two-state telegraph model in growing and dividing cells was shown to be exactly solvable when (i) the mRNA synthesis rate scales linearly with cell volume and (ii) there is no variation of gene copy numbers across the cell cycle, i.e. gene replication is not taken into account. As previously mentioned, while (i) is common, it is not universal. The assumption behind (ii) is of course a means to simplify the model but clearly unrealistic. Relaxing any one of these two properties means that within the theoretical framework presented in [49], it is not possible to obtain an exact solution for the distribution of mRNA numbers.

While the aforementioned literature summarised in Table 1 has sought to fix the biological limitations of the conventional one-state and two-state models by directly introducing more processes and solving the master equation of the resulting complex models, a different indirect approach has also been proposed. This approach takes the point of view that biological processes not explicitly modelled by the conventional models can be incorporated by considering the model parameters themselves to vary between cells, and therefore to be drawn from probability distributions [4, 56–58] — we call this an extrinsic noise model (ENM). This model can be solved exactly in steady-state for various distributions of parameter values (see Table I of [58]). It is expected that such an approach produces meaningful results provided the parameters controlling cell-to-cell variability change very slowly. Under certain conditions, the solution of the ENM might even exactly match that of complex models of stochastic gene expression. For example, it has recently been shown that the exact solution of the two-state telegraph model in growing and dividing cells where gene replication is ignored and where the mRNA synthesis rate scales with cell volume is precisely the same as that of the ENM with the mRNA synthesis rate sampled from the distribution of cellular volume and with the mRNA degradation rate being replaced by an effective rate that also incorporates the dilution of molecules at cell division [49]. A natural question is, if in a two-state telegraph model we introduce gene replication and allow the mRNA synthesis rate potentially to be volume-dependent, then does the ENM still provide an exact or at least an accurate approximation of this model?

In this paper, we first exactly solve an extension of the telegraph model that explicitly describes cell birth, growth, division, replication, and an mRNA synthesis rate that can be either independent of cell volume or else that linearly scales with it. Many of the known exact solutions of the one-state and two-state models to-date can be shown to be special cases of the present theory. The analytical distribution of transcript numbers is subsequently used to study the accuracy of the ENM. We show that the transcript number distribution in steady-state growth is generally not well approximated by the steady-state distribution of the ENM. Conditions under which the ENM provides an accurate approximation are derived and verified using simulations.

## Results

### Model

We consider an extension of the telegraph model which takes into account cell growth, cell division, gene replication, gene dosage compensation, and volume-dependent transcription (see Fig. 1 for an illustration). The specific meaning of all model parameters can be found in Table 2. The model has the following properties.

**Figure 1:**
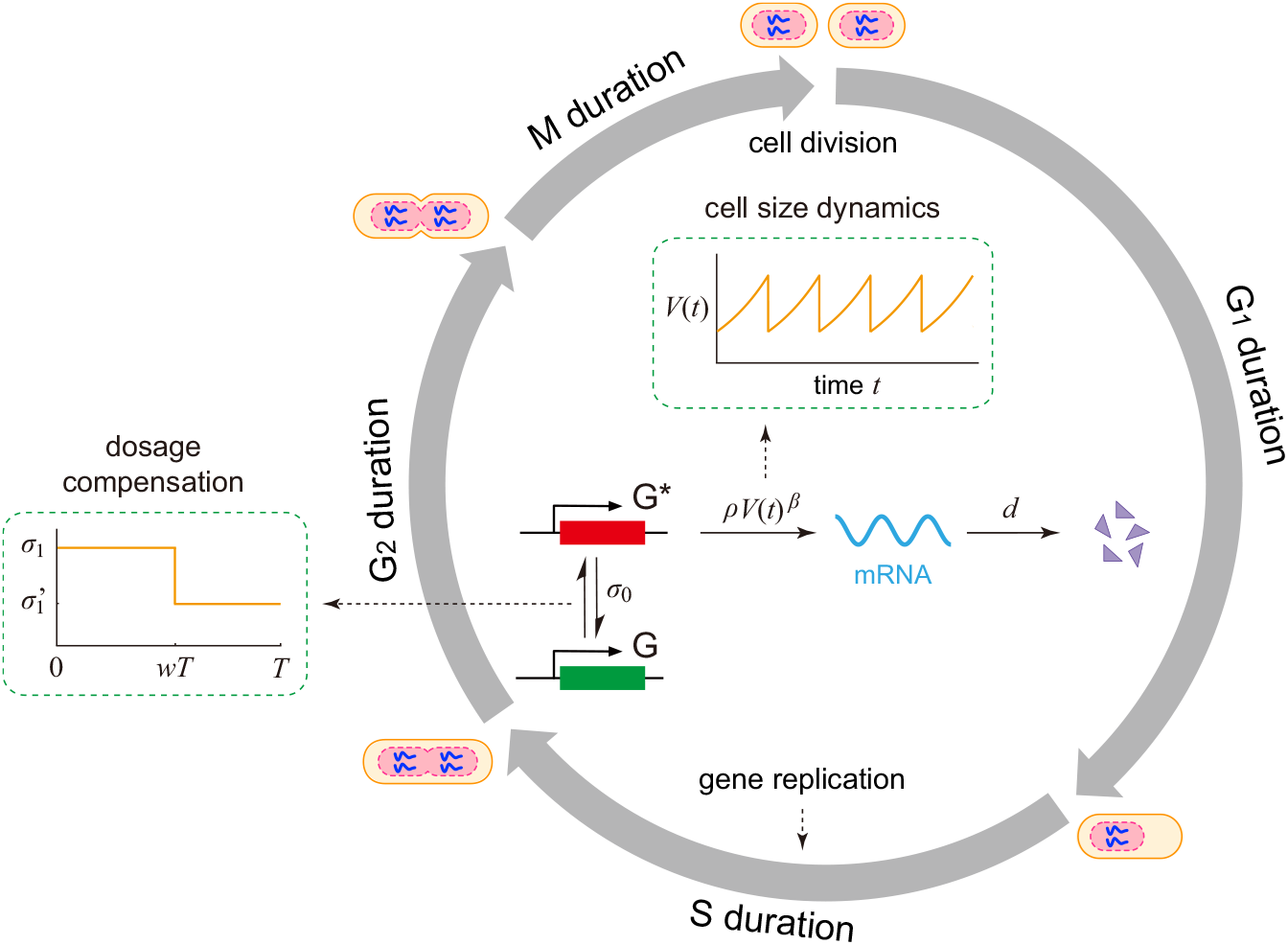
Model. Schematic of an extension of the telegraph model of gene expression in growing and dividing cells. The volume *V* (*t*) of a cell grows exponentially with constant growth rate *g* and doubling time *T*. The gene expression dynamics is characterized by a two-state model with volume-dependent transcription and volume-independent degradation. Specifically, the gene can switch between an active state *G*\* and an inactive state *G*. Transcription occurs when the gene is active. The synthesis rate of mRNA depends on cell volume *V* (*t*) via a power law form with power *β* ∈ [0,1], and the degradation rate of mRNA is a constant. Gene replication occurs at a time *T*_0_ where *w* = *T*_0_/*T* ∈ (0,1) is some fixed proportion of the cell cycle. Upon replication, the activation rate for each gene copy decreases from *σ*_1_ to 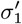 due to gene dosage compensation.

**Table 2:** Model parameters and their meaning.

| parameters | meaning |
| --- | --- |
| $V_b$ | cell volume at birth |
| $g$ | growth rate of cell volume |
| $T = \log(2)/g$ | cell cycle duration |
| $\beta$ | strength of balanced mRNA synthesis |
| $w$ | proportion of cell cycle before replication |
| $\sigma_1$ | switching rate of the gene from OFF to ON before replication |
| $\sigma'_1$ | switching rate of the gene from OFF to ON after replication |
| $\sigma_0$ | switching rate of the gene from ON to OFF |
| $\rho$ | proportionality constant of the mRNA synthesis rate |
| $d$ | mRNA degradation rate |
| $d_{\text{eff}} = d + g$ | effective mRNA decay rate |
| $\eta = d/g$ | ratio of the degradation rate to the growth rate |

1) Let *T* denote the cell cycle duration and let *V*(*t*) denote the cell volume at time *t*. We assume that cell volume grows exponentially within each cell cycle, i.e. *V* (*t*) = *V*_*b*_*e*^*g*^^t^ for any 0 ≤ *t* ≤ *T*, where *V_b_* is the cell volume at birth and *g* is the growth rate. The exponential growth of cell volume is commonly observed for various types of cells [37, 39, 59, 60]. For simplicity, we assume that the doubling time T and the growth rate *g* do not involve any stochasticity [22]. Generalization of the model to stochastic cell volume dynamics will be discussed at the end of the paper.

2) In each cell cycle, we use a two-state model to describe the gene expression dynamics. Let *G* and *G*\* denote the inactive and active states of the gene, respectively, and let *M* denote the corresponding mRNA. Consider a gene expression model described by the effective reactions

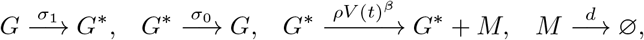

where *σ*_0_ and *σ*_1_ are the switching rates between the two gene states, and *d* is the mRNA degradation rate. For many genes in fission yeast [28, 29], mammalian cells [30, 31], and plant cells [32], there is evidence that the mRNA number scales linearly with cell volume in order to maintain approximately constant concentrations (concentration homeostasis; for a recent review see [61]). This is due to a coordination of the mRNA synthesis rate with cell volume — we shall refer to this mechanism as balanced mRNA synthesis. However, in both prokaryotic [50, 51] and eukaryotic cells [52–55] there are examples where there is no such scaling. Since each cell has a different volume, the mechanism of volume-dependent transcription is a source of extrinsic noise [57], potentially accounting for a significant amount of the observed cell-to-cell variation in mRNA numbers. To unify non-balanced and balanced mRNA synthesis, we assume that the mRNA synthesis rate depends on cell volume *V(t)* via a power law form with proportionality constant *ρ* and power *β* ∈ [0,1]. Then *β* =1 (*β* = 0) corresponds to the situation where the mRNA synthesis rate scales linearly with cell volume (does not depend on cell volume). It has recently been postulated that the non-linear scaling between gene expression levels and cellular volume is due to the heterogeneous recruitment abilities of promoters to RNA polymerases [62].

3) The replication of the gene of interest occurs at a fixed proportion *w* ∈ (0,1) of the cell cycle. This is known as a stretched cell cycle model, which is supported by experiments [63]. Under this assumption, the time before replication within a cell cycle is *wT* and the time after replication is (1 - *w*)*T*. We shall refer to the gene copy that is replicated as the mother copy and to the duplicated gene copies as the daughter copies. For haploid cells, there is only one mother copy before replication and two daughter copies after replication; for diploid cells, the number of gene copies varies from two to four upon replication. For diploid cells, we assume that the two alleles act independently of each other [64, 65].

4) At replication, the daughter copies inherit the gene state from the mother copy [22, 66]. The presence of specific histone marks dictate transcription permissiveness [67] and the landscape of histone modifications is copied during DNA replication [68]. An alternative case is the one where all daughter copies are reset to the inactive state upon replication — potentially a mechanism to avoid the risk of conflict between replication and transcription (and the resulting DNA damage) [22]. Here we only consider the former perfect state copying mechanism.

5) A doubling of gene copy numbers upon replication would be expected to also double the amount of mRNA molecules. However, experiments show that this is not always the case [26, 30, 69] principally due to a decrease of the gene activation rate upon replication, a phenomenon known as gene dosage compensation. We model this by choosing the gene activation rate before replication *σ*_1_ to be potentially different than that after replication 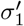. In the absence of dosage compensation, we have 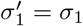. Perfect dosage compensation occurs when 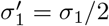; in this case, the total burst frequency for all gene copies is unaffected by replication (since the gene copy number doubles upon replication, the burst frequency for each gene copy halves when dosage compensation is perfect).

6) At division, the mother cell is divided into two daughter cells. The volumes of the two daughter cells are assumed to be the same and exactly one half of the volume of the mother cell before division (of course there is some stochasticity in the partitioning of cell size [70, 71] which we are here ignoring). Moreover, we assume that each mRNA molecule has probability 1/2 of being allocated to each daughter cell. With this assumption, the number of transcripts that are allocated to each daughter cell has a binomial distribution. We also assume that gene state is not changed upon cell division.

### Time-dependent mRNA distribution within a cell cycle

Here we compute the time-dependent distribution of the mRNA number within a cell cycle under arbitrary initial conditions. We first consider the dynamics before replication for haploid cells. The microstate of the gene of interest can be described by an ordered pair (*i, n*), where *i* denotes the state of the gene with *i* = 0,1 corresponding to the inactive and active states, respectively, and n denotes the number of mRNA molecules. Let *p_i,n_*(*t*) denote the probability of having n transcripts at time *t* ∈ [0, *wT*] when the gene is in state i. Note that *t* = 0 corresponds to cell birth. Then the stochastic gene expression dynamics before replication is governed by the coupled set of CMEs

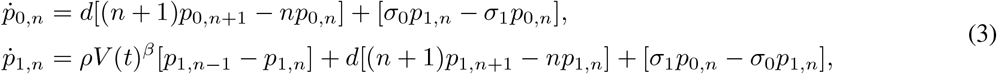

where *p*_1,-1_ = 0 by default, the term involving *ρ* represents mRNA synthesis, the terms involving *d* represent mRNA degradation, and the terms involving *σ*_0_ and *σ*_1_ represent gene state switching, which will be referred to simply as gene switching in what follows. To solve them, we define a pair of generating functions 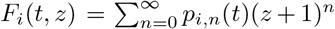 for *i* = 0,1. Note that here we use (*z* + 1)^*n*^ rather than the conventional *z^n^* in the definition of the generating function — with this choice, the formulas given below are much more concise. In addition, let *p_n_*(*t*) = *p*_0,*n*_(*t*)+*p*_1,*n*_(*t*) denote the probability of having *n* transcripts at time *t* and let *F* (*t, z*) = *F*_0_(*t, z*)+*F*_1_(*t, z*) be the corresponding generating function. In terms of the generating functions, Eq. (3) can be converted into the first-order linear partial differential equations (PDEs)

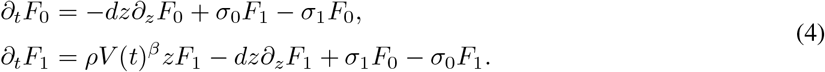

To solve them, we first convert them into a second-order parabolic PDE and then transform the second-order PDE into a hypergeometric differential equation through a change of variables. Complicated computations show that for each *t* ∈ [0, *wT*], the generating functions *F_i_, i* = 0,1 can be computed in closed form as (see Supplementary Section 1 for the proof)

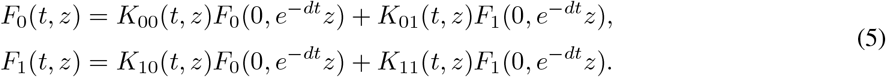

Here *F_i_*(*O, z*), *i* = 0,1 are the generating functions at *t* = 0 which can be determined by the initial conditions, and the functions *K_ij_*, *i,j* = 0,1 are given by

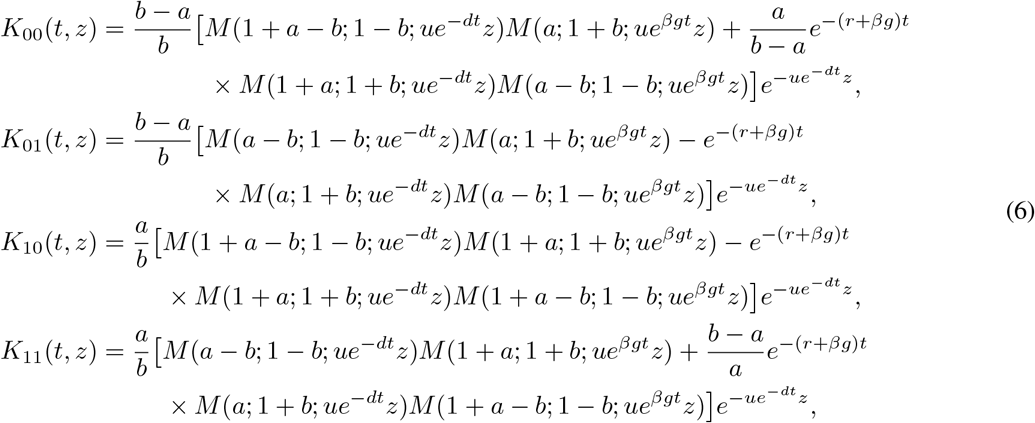

where the parameters *r, a, b,* and *u* are given by

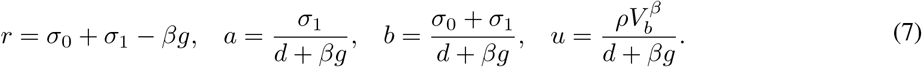

Adding the two identities in Eq. (5) gives the explicit expression of the generating function F before replication, i.e.

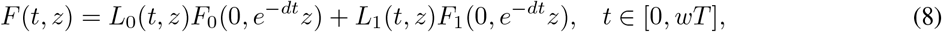

where the functions *L_i_, i* = 0,1 are given by

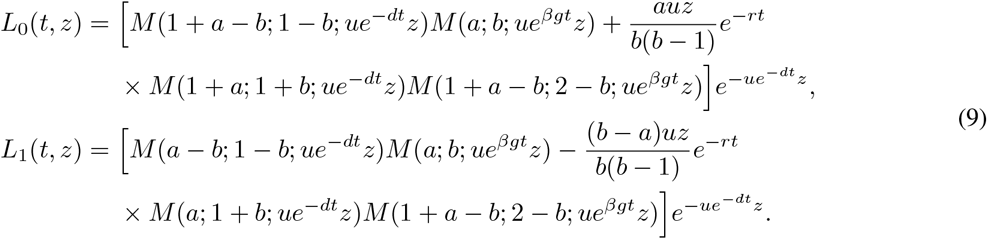

When *b* = 1, the term *b* – 1 appears in the dominator of these equations and the equalities should be understood in the limiting sense. Note that when the mRNA synthesis rate is volume-independent (*β* = 0), the expression of *F* given in Eq. (8) coincides with the time-dependent solution of the standard telegraph model [14] (the analytical solution given in [14, Eq. (5)] has a typo where a plus sign should be replaced by a minus sign and the correct one can be found in [15, Eq. (75)]).

We next focus on the dynamics after replication for haploid cells. Since there are two daughter gene copies after replication, to distinguish them, we call them daughter copy *A* and daughter copy *B.* The dynamics of each gene copy is governed by the CMEs given in Eq. (3) with *σ*_1_ being replaced by 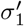. Let *p_n_*(*t*) denote the probability of having *n* transcripts at time *t* ∈ [*wT, T*] and let 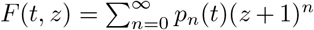 be the corresponding generating function. In Supplementary Section 2, we prove that the generating function *F* after replication can be computed in closed form as

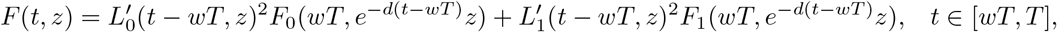

where 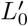 and 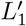 are functions obtained from *L*_0_ and *L*_1_ by replacing the parameters *r, a, b* and *u* with

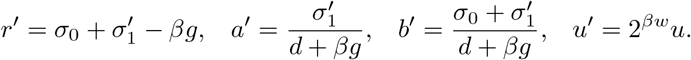

In summary, we have derived the analytical expression of the generating function *F* at any time *t* ∈ [0, *T*] within a cell cycle, which is given by

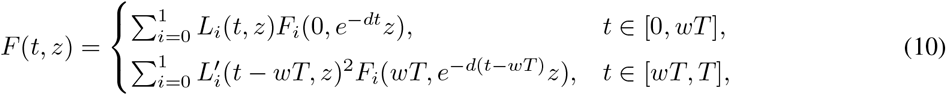

where *F_i_*(*wT, z*), *i* = 0,1 are determined by Eq. (5). The time-dependent distribution of the mRNA number can be recovered by taking the derivatives of the generating function *F* at *z* = −1, i.e.

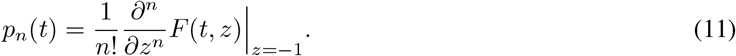

Our analytical expression of the transient mRNA distribution is rather complicated. However, it can be greatly simplified in some special cases. In Methods, we show how the analytical solution can be simplified for two non-trivial special cases: (i) the gene switches rapidly between the active and inactive states (*σ*_0_, *σ*_1_ ≫ *g*); (ii) the mRNA is produced in a bursty manner (*σ*_0_ ≫ *σ*_1_), i.e. the gene is mostly inactive but transcribes a large number of mRNA when it becomes active [72–75]. In the latter case, the burst frequency is *σ*_1_ before replication and the total burst frequency for the two gene copies is 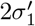 after replication.

Thus far, we have obtained the transient mRNA distribution for haploid cells. For diploid cells, since the two alleles act independently and since each allele has the mRNA distribution given in Eq. (11), the generating function for the total number of transcripts at any time *t* ∈ [0, *T*] is given by *F*_diploid_(*t, z*) = *F*(*t, z*)^2^, where *F*(*t, z*) is given by Eq. (10). Here we have used the fact that the generating function of two independent random variables is the product of their respective generating functions. Due to independence of the two alleles, when the rate parameters for each allele are fixed, the gene expression noise (measured by the coefficient of variation squared of mRNA numbers) in diploid cells is one half that in haploid cells. Without loss of generality, we always focus on haploid cells in what follows.

### Time-dependent mRNA distribution across cell cycles

Thus far, we have derived the exact mRNA distribution at any time within a cell cycle. Here we focus on the full time-dependence of the mRNA distribution across cell cycles under arbitrary initial conditions. To this end, we not only need the expression of *F* at any time *t* ∈ [0, *T*], but also need the expressions of *F_i_, i* = 0,1.

Recall that Eq. (5) gives the analytical expressions of the generating functions *F_i_, i* = 0,1 before replication under any initial conditions. In particular, at replication, we have

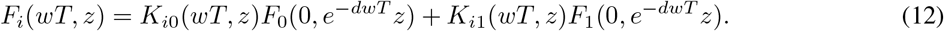

Now we focus on the dynamics of daughter copy *A* after replication. Let *p_i,n_*(*t*) denote the probability of having n transcripts at time *t* ∈ [*wT, T*] when the daughter copy A is in state *i* and let 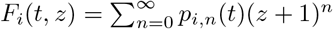 be the corresponding generating function. In Supplementary Section 2, we prove that the generating functions *F_i_, i* = 0, 1 after replication can be computed exactly as

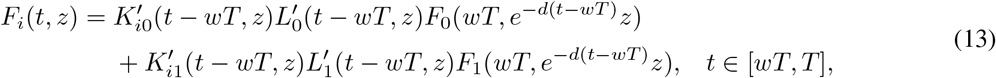

where 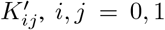 are functions obtained from *K_ij_* by replacing the parameters *r, a, b*, and *u* with the parameters *r*’, *a*’, *b*’, and *u*’, respectively. Inserting Eq. (12) into Eq. (13) and taking *t* = *T*, we obtain

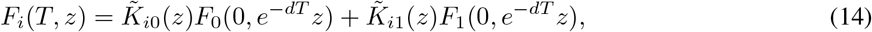

where

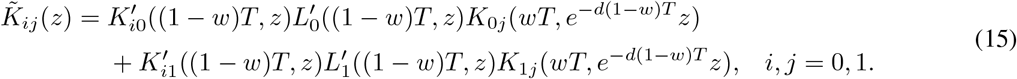

Suppose that the daughter cell with daughter copy A is tracked after division. Since we have assumed binomial partitioning of molecules at division, the probability 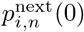 at birth in the next generation is given by

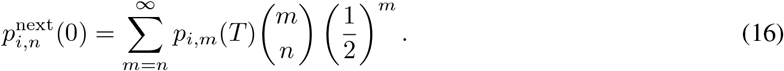

In terms of the generating function, the above relation can be written as

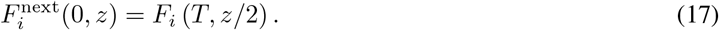

This gives the initial conditions for the next generation and the time-dependent mRNA distribution within the next cell cycle can be computed via Eq. (10). Applying Eqs. (14), and (17) repeatedly, we are able to compute the full time-dependence of the *F_i_* functions across cell cycles; substituting these in Eq. (10) gives the full-time dependence of the mRNA distribution across cell cycles.

As a check of our analytical solutions, we compare the exact distributions of the mRNA number with the numerical ones obtained from a modified version of the finite-state projection (FSP) [76] algorithm at three different time points (birth, replication, and division) across four cell cycles (Fig. 2). In this algorithm, we couple the standard FSP with cell cycle events; for details see Supplementary Section 3. Here we assume that initially there is no mRNA molecules in the cell and the gene is off. This mimics the situation where the gene has been silenced by some repressor over a period of time such that all transcripts have been removed via degradation (while after silencing there may be some background level of mRNA, for simplicity we assume that all transcripts have been degraded). At time *t* = 0, the repressor is removed and we study how gene expression recovers. When using FSP, we truncate the state space (to exclude states that are visited very rarely) and solve the associated truncated master equation numerically using the MATLAB function ODE45 with the dynamics before and after replication solved separately. Note that while the FSP and the stochastic simulation algorithm (SSA) yield comparable distributions of molecule numbers, the computational time of the former is much less than of the latter provided the biochemical reaction networks are small enough — hence here we used the FSP. As expected, the analytical and simulated solutions coincide with each other completely at all times, and the mRNA distributions at birth, replication, and division reach a steady state within a few cell cycles. This can be also seen from Fig. S1, where we illustrate the time-dependent mean and Fano factor of the mRNA number across four cell cycles.

**Figure 2:**
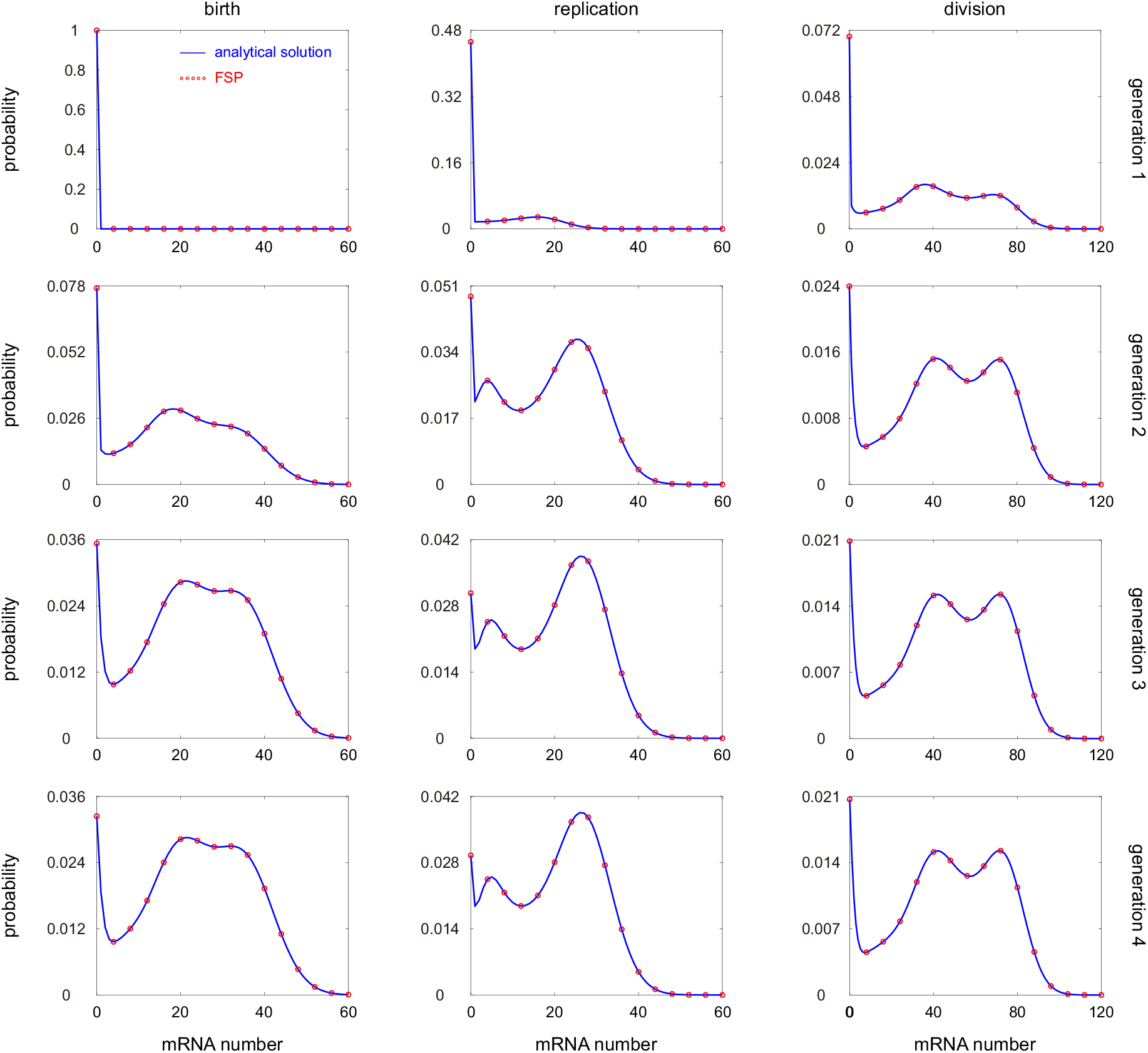
Time-dependent mRNA distributions at birth, replication, and division across four cell cycles. The blue curves show the analytical distributions computed by applying Eqs. (10), (14), and (17) repeatedly, and the red circles show the numerical ones obtained from FSP. The model parameters are chosen as *V_b_* = 1, *g* = 1, *β* = 1, *w* = 0.4, *d* = 5, *ρ* = 20 *d*_eff_, *σ*_0_ = 1.5, *σ* = _3_, 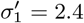.

Another interesting observation is that the time-dependent mRNA distributions for our detailed telegraph model may exhibit three modes (Fig. 2) – this is the combined effect of gene replication and slow switching between gene states. The zero mode is since there is no transcription when the gene is off while the two non-zero modes are due to transcription when the gene is turned on during the pre-replication and post-replication phases of the cell cycle. According to simulations, distributions with more than three modes are not observed in our model. This is different from the prediction of the conventional telegraph model [12] whose distribution has at most two modes.

### Time-dependent mRNA distribution under cyclo-stationary conditions

Thus far, we have obtained the full time-dependence of the mRNA distribution across cell cycles under arbitrary initial conditions. After several generations, the distribution at any fixed time within a cell cycle (such as the distributions at birth, replication, and division) becomes independent of the generation number. This is also called the cyclo-stationary condition in the literature [46] or steady-state growth [20]. Next we compute the time-dependent mRNA distribution within a cell cycle under cyclo-stationary conditions.

Before computing the mRNA distribution, we first derive the probabilities of the gene being in the active and inactive states at any time within a cell cycle under cyclo-stationary conditions. Let *p*_on_(*t*) denote the probability of each gene copy being in the active state at time *t* ∈ [0, *T*]. Before replication, the dynamics of the active probability satisfies the differential equation 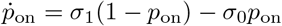. Solving this equation gives rise to

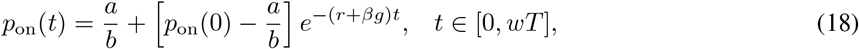

where we have used the fact that *a*/*b* = *σ*_1_/(*σ*_0_ + *σ*_1_) and *r* + *βg* = *σ*_0_ + *σ*_1_ (see Eq. (7)). Recall that the gene activation rate decreases from *σ*_1_ to 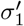 upon replication. After replication, the dynamics of the active probability satisfies the differential equation 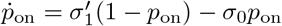. Solving this equation yields

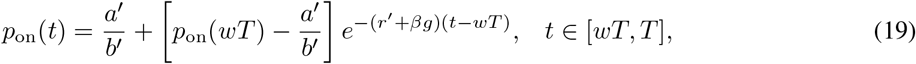

where *p*_on_(*wT*) is determined by Eq. (18). Combining Eqs. (18) and (19), we obtain the active probability of the gene at division, i.e.

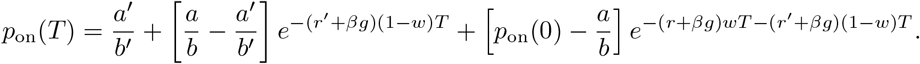

Under cyclo-stationary conditions, the active probabilities at cell birth in two successive generations must be the same, i.e. 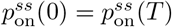. Then the steady-state active probability of the gene at birth is given by

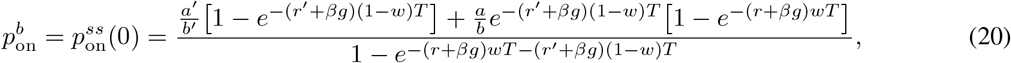

and thus the steady-state inactive probability at birth is given by 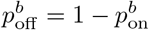. It then follows from Eq. (18) that the steady-state active probability of the gene at replication is given by

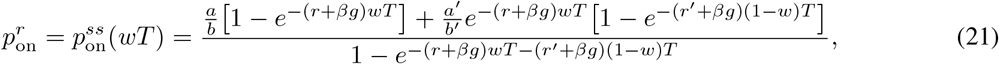

and thus the steady-state inactive probability at replication is given by 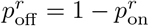.

Next we focus on the time-dependent mRNA distributions under cyclo-stationary conditions. Recall that we have obtained the time-dependent mRNA distributions within a cell cycle, whose generating function *F*(*t, z*) is given by Eq. (10), provided that the initial conditions *F_i_*(0, *z*), *i* = 0,1 are known. Under cyclo-stationary conditions, the values of *F_i_*(0, *z*) in two successive generations must be the same, i.e. 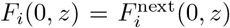, where 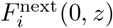 has been derived in Eqs. (14) and (17). It then follows that the steady-state values of *F_i_* (0, *z*) should satisfy

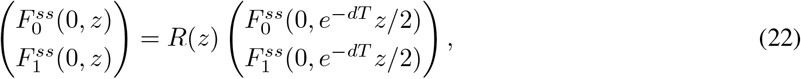

where

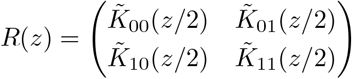

is a matrix-valued function with 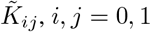 being given in Eq. (15). Applying Eq. (22) repeatedly, we obtain

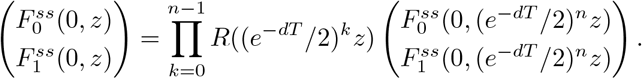

Taking *n* → ∞ in the above equation yields

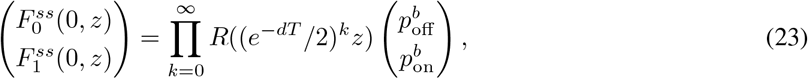

where we have used the fact that

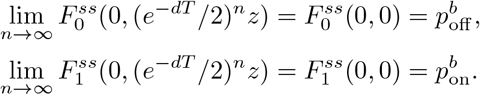

Once we have derived the steady-state values of *F_i_*(0, *z*), *i* = 0,1, it immediately follows from Eq. (10) that the time-dependent generating function *F* under cyclo-stationary conditions is given by

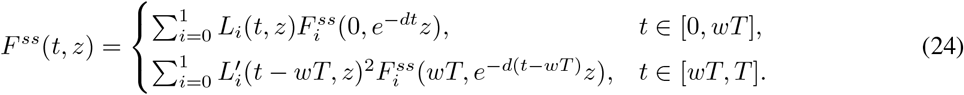

### Comparison with the effective dilution model

#### Special case 1

Consider the case where gene replication is not taken into account (*w* = 1) and when the mRNA synthesis rate scales with cell volume (*β* = 1) [49]. In this case, the functions 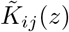 given in Eq. (15) reduce to 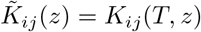, and it is not difficult to see that Eq. (22) can be solved analytically as

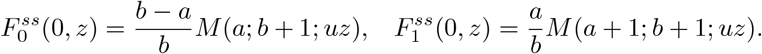

Inserting these equations into Eq. (24) yields

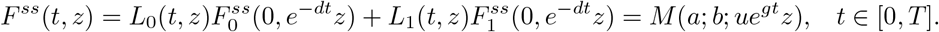

For a given cell of volume *V*, its age is given by *t* = log(*V/V_b_*)/*g*. Substituting *t* = log(*V/V_b_*)/*g* in the above equation shows that the steady-state generating function for a cell of constant volume *V* is given by 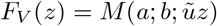, where *a* = *σ*_1_/(*d* + *g*), *b* = (*σ*_0_ + *σ*_1_)/(*d + g*), and 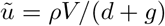. We make a crucial observation that this is exactly the steady-state generating function of the mRNA distribution for the conventional telegraph model [12]

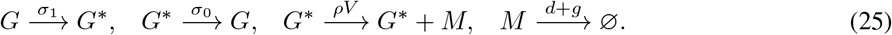

This result has been found in [49], which states that when *w* = *β* = 1, the steady-state mRNA distribution for a cell of constant volume *V* of the detailed telegraph model is the same as that of the conventional telegraph model with effective decay rate *d*_eff_ = *d* + *g*. Note that the two terms in this rate capture the fact that transcripts are lost both by active degradation (with rate *d*) and by dilution at cell division (with rate *g*) – hence a model of this type is known as an effective dilution model (EDM) [77]. Intuitively, the EDM considers a population of cells with synchronised cell cycles so that at each time, all cells have the same volume.

#### Special case 2

Experiments have shown that in bacteria, most mRNAs have a half-life that is much shorter than the cell cycle duration, i.e. *d* ≫ *g* (see Supplementary Section 4 for the typical values of *d* and *g* in various cell types), and thus are very unstable. The value of *η* = *d/g* can be used to measure the stability of mRNA. For unstable mRNAs (*η* ≫ 1), the terms *e*^-*dt*^ and *e*^-*d*(*t*-*wT*)^ in Eq. (10) are very small and thus can be approximated by zero (whenever *t* is not very close to 0 and *wT*). In this case, the time-dependent generating function *F* under cyclo-stationary conditions reduces to

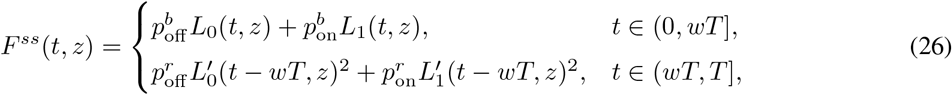

where we have used the fact that 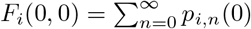 and 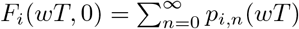 are the probabilities of the gene being in state *i* at birth and at replication, respectively. Imposing the term *e*^-*dt*^ as zero in Eq. (9) yields

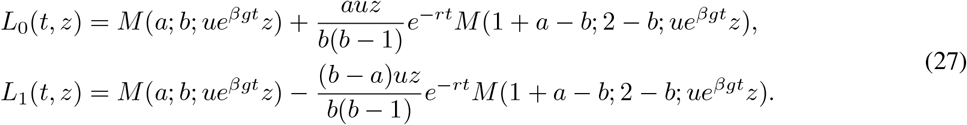

When one of the gene switching rates *σ*_0_ and *σ*_1_ is very large, we have *r* = *σ*_0_ + *σ*_1_ - *βg* ≫ *g* and thus the second term on the right-hand side of Eq. (27) can be neglected. This may occur when (i) the gene switches rapidly between the two states (*σ*_0_, *σ*_1_ ≫ g), or (ii) the mRNA is produced in a constitutive manner (*σ*_1_ ≫ *σ*_0_, *g*), or (iii) the mRNA is produced in a bursty manner (*σ*_0_ ≫ *σ*_1_, *g*). In this case, the cyclo-stationary generating function *F^ss^* can be simplified significantly as

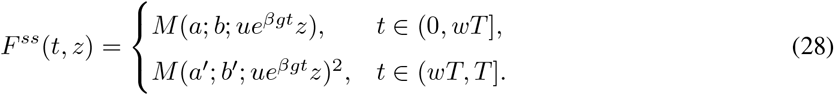

This contains much information. For a given cell of volume *V* < 2^*w*^ *V_b_*, its age is given by *t* = log(*V/V_b_*)/*g* < *wT* and hence there is only one gene copy in the cell. Substituting *t* = log(*V/V_b_*)/*g* in the above equation shows that the steady-state generating function for a cell of constant volume *V* is given by 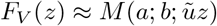, where

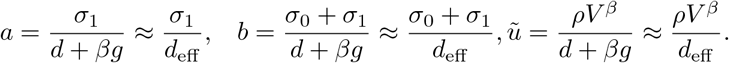

Here we have used the fact that *d*_eff_ /(*d* + *g*) ≈ 1 when mRNA is very unstable. Note that *F_V_* (*z*) is exactly the steady-state generating function of the mRNA distribution for the EDM

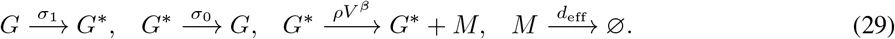

On the other hand, for a given cell of volume *V* > 2^*w*^ *V_b_*, its age is given by *t* = log(*V/V_b_*)/*g* > *wT* and hence there are two gene copies in the cell. In this case, the EDM should be modified as

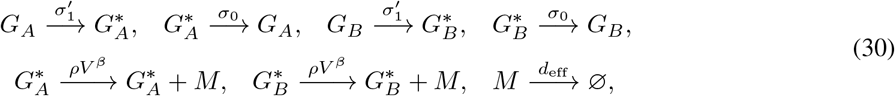

where *G_A_* and *G_B_* denote the two daughter copies whose dynamics are both governed by the conventional telegraph model. Substituting *t* = log(*V/V_b_*)/*g* in Eq. (28) shows that the steady-state generating function for a cell of constant volume *V* is given by 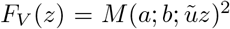, where 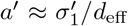 and 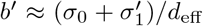. Note that *F_V_* (*z*) is exactly the steady-state generating function of the mRNA distribution for the EDM given in Eq. (30) since the two gene copies are independent of each other.

In summary, our analysis shows that for mRNAs with short lifetimes, the EDM makes a good approximation when one of the gene switching rates *σ*_0_ and *σ*_1_ is large (here the cell age *t* cannot be very close to 0 and *wT*, i.e. newborn cells and cells that have just finished gene replication should be excluded). This can be understood as follows. Previous studies [78] have shown that the relaxation speed of the EDM to the steady state is governed by both the mRNA degradation rate *d* and the total gene switching rate *σ*_tot_ = *σ*_0_ + *σ*_1_. When *d* and *σ*_tot_ are both large, any memory at birth from the previous cycle (due to binomial partitioning of molecules at division and to the gene state prior to division) and any memory at replication (due to gene state copying of the two daughter copies) will be rapidly erased. Each time that the volume changes, the mRNA distribution instantaneously equilibrates and hence the EDM works. Note that when the cell age t is close to 0 and *wT*, the memory at birth and at replication cannot be erased, which leads to the failure of the EDM. Relatively slow mRNA degradation and relative slow gene switching will both result in a deviation of the EDM from the full model.

#### Testing the accuracy of the EDM approximation

In Fig. 3 we compare the exact mRNA distributions with the numerical ones obtained from FSP at three different time points (birth, replication, and division) across the cell cycle under cyclo-stationary conditions. The truncated master equations are solved across several (usually less than five) cell cycles until the Hellinger distance between mRNA distributions at birth in two successive generations is less than 10^-4^. This guarantees that cyclo-stationary conditions are reached. When gene replication is not taken into account (*w* = 1) and when the mRNA synthesis rate scales with cell size (*β* = 1), the distributions of the full model agree perfectly with those of the EDM given in Eq. (25) (Fig. 3(a)). This coincides with our theoretical predictions. When gene replication is taken into account, the EDMs before and after replication are given by Eqs. (29) and (30), respectively. In this case, the EDM may deviate remarkably from the full model with the deviation being much larger at early stages of the cell cycle (Fig. 3(b)), especially when mRNA degradation and gene switching are relatively slow. This can be understood as follows. According to the steady-state properties of the conventional telegraph model, in the presence of gene replication, the mean and the Fano factor of the mRNA number at birth for the EDM are given by

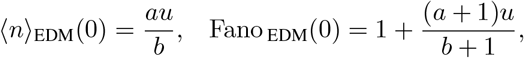

and the mean and the Fano factor of the mRNA number at division are given by

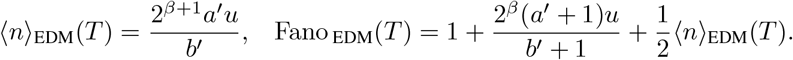

**Figure 3:**
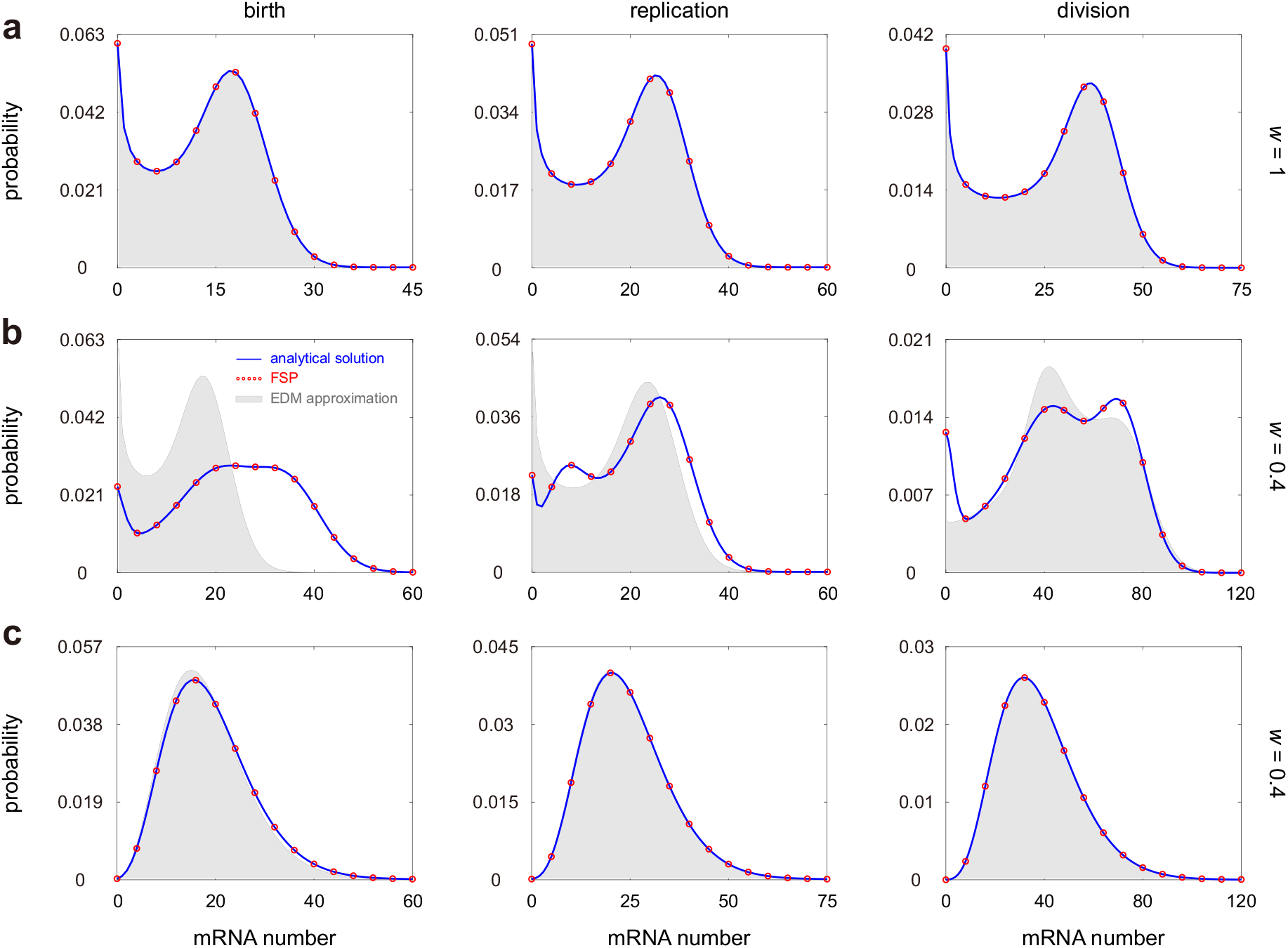
Comparison between the full model and the EDM. (**a**) Steady-state mRNA distributions at birth, replication, and division for the full model and the EDM when gene replication is not taken into account. The blue curves show the analytical distributions given in Eqs. (23) and (24), the red circles show the numerical ones obtained from FSP, and the grey regions show the distributions of the EDM. (**b**) Same as (a) but when gene replication is taken into account. In (a),(b), the model parameters are chosen as *V_b_* = 1,*g* = 1, *β* = 1,*d* = 4,*ρ* = 20 *d*_eff_, *σ*_0_ = 1.5, *σ*_1_ = 3, 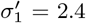. The parameter *w* is chosen as *w* = 1 in (a) and *w* = 0.4 in (b). (**c**) Same as (b) but in the special case where mRNA synthesis is balanced and bursty, and dosage compensation is perfect. The model parameters are chosen as *V_b_* = 1, *g* = 1, *β* = 1, *w* = 0.4,*d* = 4, *ρ* = 200*d*_eff_, *σ*_0_ = 300, *σ*_1_ = 30, 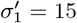.

Under cyclo-stationary conditions, it follows from Eq. (16) that the mean mRNA numbers at birth and at division for the full model should satisfy 〈*n*〉 (*T*) = 2〈*n*〉 (0) and Fano (*T*) = 2Fano (0) −1. However, these two restrictions in general do not hold for the EDM – the EDM satisfies these two restrictions only when

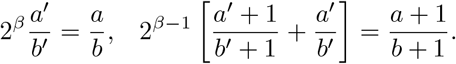

Note that when mRNA synthesis is balanced (*β* = 1) and bursty (*σ*_0_ ≫ *σ*_1_), the above restrictions are satisfied when dosage compensation is perfect 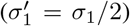, i.e. when the total burst frequency does not change when replication occurs. When these three conditions are satisfied, the EDM makes accurate predictions and the mRNA number follows a negative binomial distribution (Fig. 3(c)). The breakdown of the above restrictions will give rise to the deviation of the EDM from the full model, as observed in Fig. 3(b). Intuitively, this is because the mRNA distribution at birth is affected by the fluctuations of the two gene copies at division and thus in general it cannot be captured solely by an EDM with only one gene copy. Note that special case 2 discussed above may not satisfy the above moment equalities since in this special case, the EDM fails for newborn cells.

### Distribution and moment analysis for lineage and population measurements

We next compute the steady-state distributions of transcript numbers measured over a cell lineage or from a population snapshot. In lineage measurements, the mRNA number from an individual cell is tracked at any point in time, i.e. once the cell divides, only one of the two daughter cells is tracked. Clearly, the probability of observing a cell of age *t* ∈ [0, *T*] is 1/*T* for lineage measurements. As a result, the generating function of the steady-state distribution along a cell lineage is given by

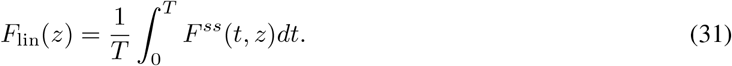

In contrast, in population measurements, the mRNA numbers in a population of isogenic cells are observed at a particular time. Previous studies [20] have shown that the probability of observing a cell of age *t* ∈ [0, *T*] is 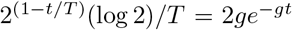 for population measurements. Thus the generating function of the steady-state distribution in a population of cells is given by

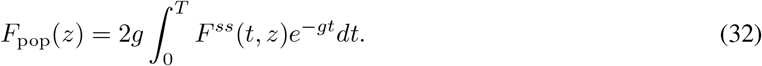

Our analytical expression of the steady-state distribution is rather complicated since we have to integrate the time-dependent distribution over time which involves complex confluent hypergeometric functions. However, it can be simplified to a large extent in some special cases. In Methods, we show how the analytical solution can be simplified in two non-trivial special cases: (i) the mRNA is unstable and the gene switches rapidly between the two states; (ii) the mRNA is unstable and the gene switches slowly between the two states. In particular, in the latter case, the the steady-state distribution for lineage measurements is given by

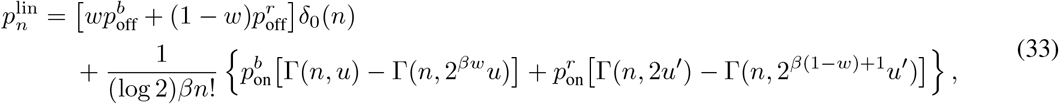

and the steady-state distribution for population measurements is given by

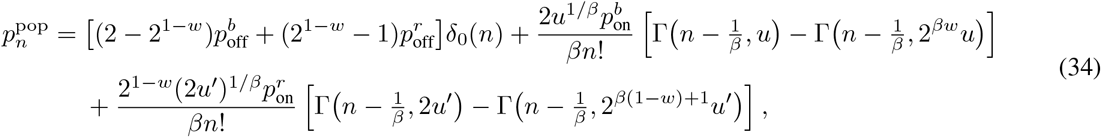

where 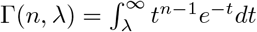 is the incomplete gamma function and *δ*_0_(*n*) is the Kronecker delta which takes the value of 1 when *n* = 0 and takes the value of 0 otherwise. Interestingly, for the two types of measurements, the mRNA distribution has a zero-inflated part (the first part). Indeed, numerous biostatistical papers [79–81] have used zero-inflated models to characterize mRNA distributions in single-cell RNA-sequencing (scRNA-seq) data analysis. For population data such as in scRNA-seq, our theory predicts that the probability of true zeros, i.e. dropout events that are not due to purely technical reasons, is 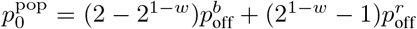 This is important for interpreting scRNA-seq data and may be potentially useful for data imputation.

Fig. 4(a),(b) compare the population mRNA distribution given in Eq. (34) and the numerical one obtained from FSP. Clearly, the two distributions agree perfectly when the transcripts are highly unstable and when the gene switching rates are very small. Particularly, we find that the mRNA distribution is capable of exhibiting three modes, corresponding to the three terms in Eq. (34) (left panel of Fig. 4(a)). This shows that a trimodal distribution may occur in the special case of unstable mRNA and slow gene switching. When the transcripts are relatively stable, trimodality becomes less apparent and Eq. (34) fails to capture the real mRNA distribution, while it can still well capture the probability of zero observations (right panel of Fig. 4(a)).

**Figure 4:**
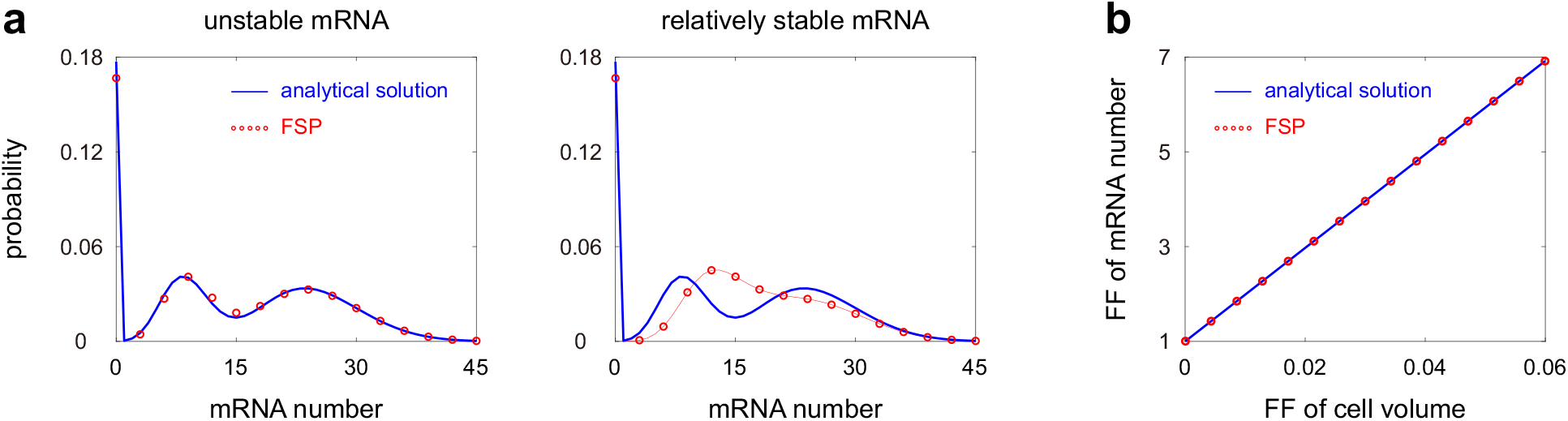
Distribution and moment analysis for population measurements. (**a**) Steady-state mRNA distribution in a cell population when gene switching is very slow. The blue curves show the analytical distributions given in Eq. (34) and the red circles show the numerical ones obtained from FSP. The model parameters are chosen as *V_b_* = 1, *g* = 1, *β* = 1, *w* = 0.3, *ρ* = 8 *d*_eff_, *σ*_0_ = 2 × 10^-4^, *σ*_1_ = 10^-3^, 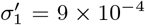. The mRNA degradation rate is chosen as *d* = 50 in the left panel and *d* = 10 in the right panel. (**b**) Fano factor of the mRNA number versus the Fano factor of cell volume in a cell population. The blue line is computed from the analytical expressions given in Eqs. (35) and (37) and the red circles are obtained from FSP. The model parameters are chosen to be the same as in the left panel of (a).

We next analyze the moments of transcript numbers. Here we consider the general case without making any simplifying assumptions. The mean and the second factorial moment of the mRNA number at any time *t* ∈ [0, *T*] within a cell cycle can be recovered by taking the derivatives of the generating function *F* at *z* = 0, i.e.

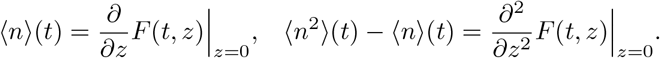

Straightforward computations show that

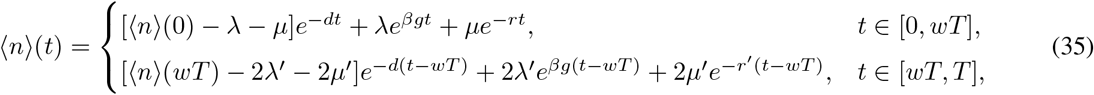

where 〈*n*〉(0) is the mean of the initial mRNA number, λ = *au/b*, *λ*’ = *a*’*u*’/*b*’, and

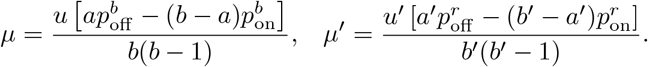

Under cyclo-stationary conditions, the mean at division should be twice that at birth, i.e. 〈*n*〉 (*T*) = 2〈*n*〉 (0). This shows that the steady-state mean at birth is given by

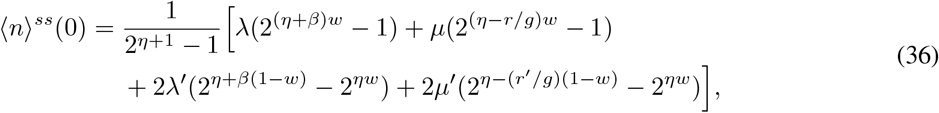

where *η* = *d/g*. Inserting this equation into Eq. (35) gives the steady-state mean at any time within a cell cycle.

The explicit expression for the second moment is extremely complicated since we need to take the second derivative of a complex generating function. However, when the mRNA is unstable, the generating function has a relative simple expression (see Eq. (26)) and taking the second derivative of this function yields the second factorial moment of the mRNA number at any time within a cell cycle:

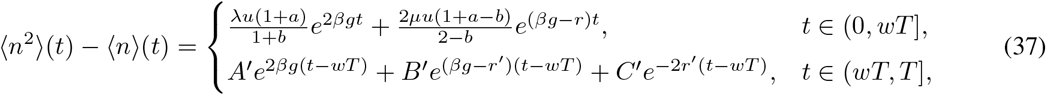

where

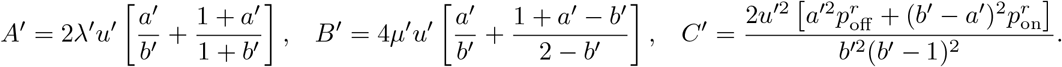

The steady-state moments of the mRNA number for lineage and population measurements can then be obtained by integrating Eqs. (35) and (37) over time. For example, the mean and second factorial moment of the mRNA number for population measurements are given by

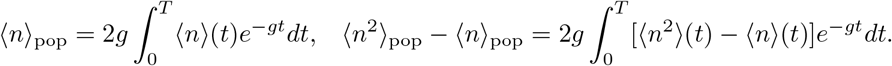

This explicit expressions can be obtained but are omitted here since they are too complicated. In Supplementary Section 5 and Fig. S2, we find that the lineage mean is always greater than the population mean, and the difference between them is at most 10%.

A crucial observation made from the analytical results is that for both types of measurements, the Fano factor of mRNA number fluctuations, FanomRNA, and the Fano factor of cell volume fluctuations, Fano_volume_, must satisfy the following relation when mRNA synthesis is balanced (see Methods for the proof):

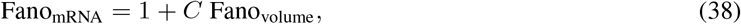

where *C* is a constant independent of the birth volume *V_b_* and the growth rate *g*. In Fig. 4(b), we validate this relation using both the exact solution and FSP. Our result shows that the fluctuations in gene expression and cell volume, characterized by the Fano factors, are linearly correlated when the mRNA synthesis rate scales with cell size. This may be potentially useful for checking whether mRNA synthesis is balanced in living organisms.

In particular, when the mRNA are unstable and when gene expression is bursty, the Fano factor of mRNA number fluctuations can be computed explicitly by integrating Eqs. (35) and (37) over time:

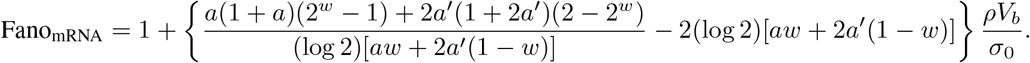

When dosage compensation is perfect, we have 2*a*′ = *a* and thus the above equation reduces to

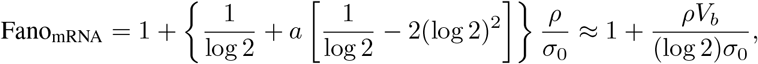

where we have used the fact that 2(log 2)^2^ ≈ 1. Note that when gene expression is bursty, the mean burst size at time *t* is *ρV*(*t*)/*σ*_0_ and hence the mean burst size over the whole cell cycle is given by *B* = (*ρV_b_*)/[(log2)*σ*_0_], which can be obtained by averaging *ρV*(*t*)/*σ*_0_ over time. In this case, we have Fano_mRNA_ ≈ 1 + *B*, which reduces to the well-known result for the conventional telegraph model [72].

### Comparison with the extrinsic noise model

Our detailed telegraph model involves the coupling between gene expression dynamics, cell volume dynamics, and cell cycle events. In Supplementary Section 6 and Fig. S3, we show that the steady-state distribution of the detailed model cannot be captured by the steady-state solution of the conventional telegraph model given in Eq. (1) with volume-independent rates, even when gene replication is not taken into account (*w* = 1). In previous studies, the lineage and population distributions for the detailed model have often been approximated by the distributions for the ENM [49]. In the ENM, the mRNA distribution for a cell of constant volume *V* is exactly the one predicted by the EDM, and the fluctuations of cell volume *V* are regarded as extrinsic noise [57, 58]. In other words, the mRNA distribution for the ENM is given by

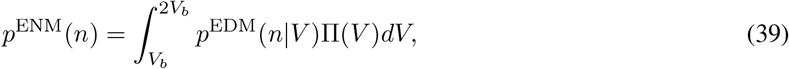

where Π(*V*) is the distribution of cell volume. We emphasize here that the EDM varies depending on the number of gene copies and thus also depending on cell volume. For a cell of volume *V* < 2^*w*^ *V_b_*, there is only one gene copy and the EDM is given by Eq. (29); for a cell of volume *V* ≥ 2^*w*^ *V_b_*, there are two gene copies and the EDM is given by Eq. (30). In addition, note that the distribution of cell volume is different for lineage and population measurements. Since cell volume *V*(*t*) and cell age *t* are related by 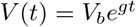, the cell volume distribution can be obtained from the cell age distribution which has already been given above (see the paragraphs before Eqs. (31) and Eq. (32)). Specifically, the volume distribution for lineage measurements is given by [70]

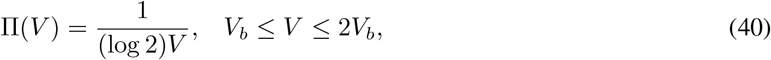

and the volume distribution for population measurements is given by

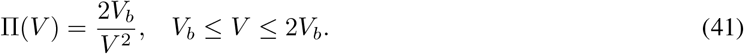

Inserting the above two equations into Eq. (39) gives the mRNA distribution for the ENM.

To evaluate the performance of the ENM approximation, we first illustrate the Hellinger distance D between the lineage distributions of the full model and the ENM as a function of *σ*_0_/*σ*_1_ and 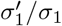 when mRNA synthesis is balanced, i.e. *β* = 1 (Fig. 5(a)). It can be seen that the ENM serves as a good approximation when gene expression is bursty (*σ*_0_ ≫ *σ*_1_) and when dosage compensation is perfect 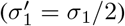. This is indeed a sufficient condition for mRNA to display concentration homeostasis when gene replication is taken into account [48]. A proof of this condition can be found in Methods. The breaking of either dosage compensation or bursty expression will lead to a significant deviation of the ENM from the full model (Fig. 5(b)). In particular, the distribution of the ENM model can show bimodality whereas that of the full model is unimodal.

**Figure 5:**
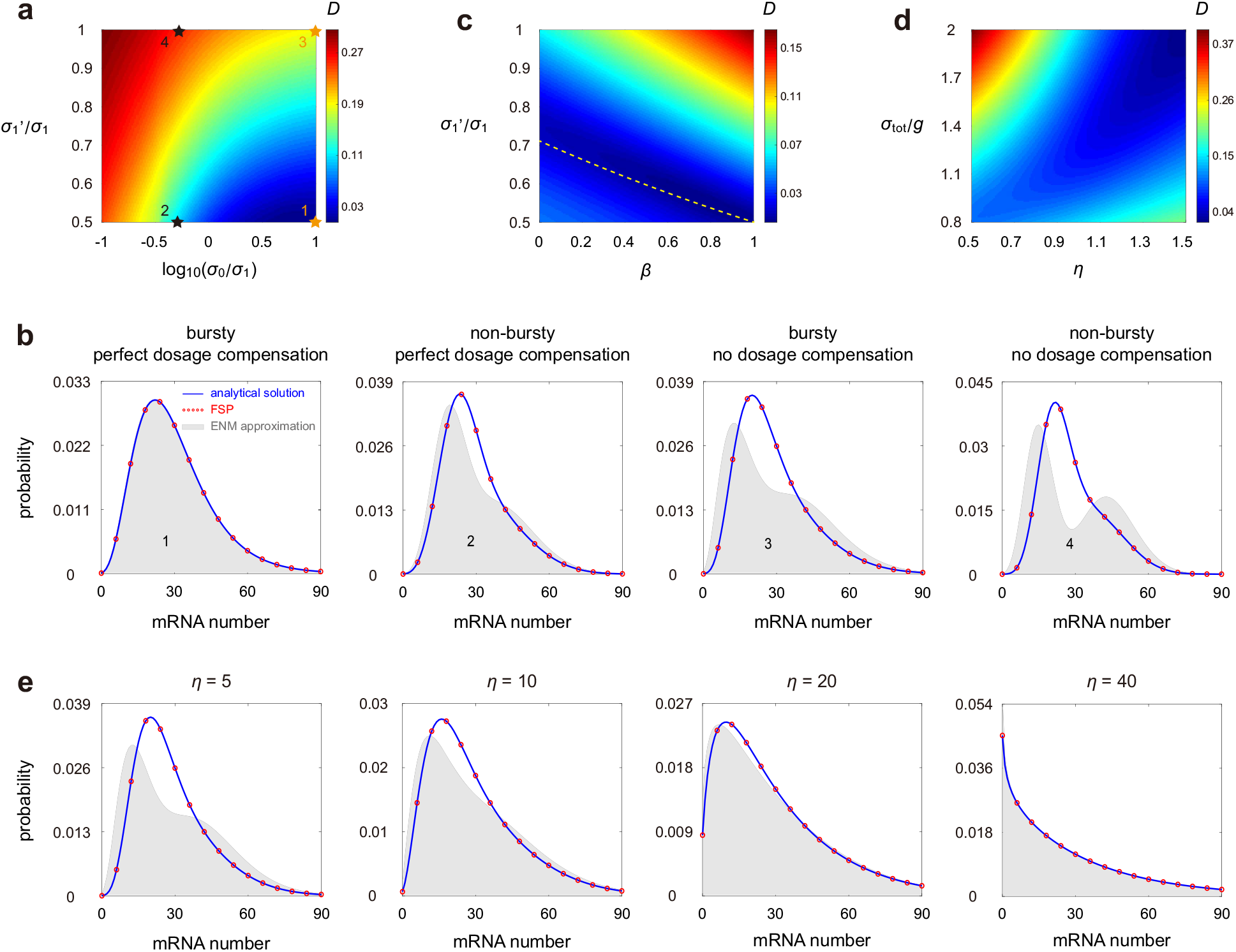
Comparison between the full model and the ENM. (**a**) Heat plot of the Hellinger distance *D* between lineage distributions for the full model and the ENM as *σ*_0_/*σ*_1_ and 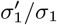 vary. The model parameters are chosen as *V_b_* = 1, *g* = 1, *β* = 1, *w* = 0.5, *d* = 5, *σ*_1_ = 30. (**b**) Comparison between the lineage distributions for the full model and the ENM as *σ*_0_/*σ*_1_ and 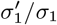 vary. The blue curves show the analytical distributions for the full model given in Eq. (31), the red circles show the numerical ones obtained from FSP, and the grey regions show the distributions for the ENM. The model parameters are chosen as in (a). The parameter *σ*_0_ is chosen as *σ*_0_ = 10*σ*_1_ (bursty case) and *σ*_0_ = 0.5*σ*_1_ (non-bursty case). The parameter 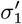 is chosen as 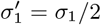 (perfect dosage compensation) and *σ*_0_ = *σ*_1_ (no dosage compensation). The parameters associated with the four panels are marked in (a) by stars. (**c**) Heat plot of *D* as *β* and 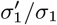 vary. The model parameters are chosen as *V_b_* = 1, *g* = 1, *w* = 0.5, *d* = 5, *σ*_0_ = 300, *σ*_1_ =30. (**d**) Heat plot of *D* as *η* and *σ*_tot_ /*g* vary. The model parameters are chosen as *V_b_* = 1, *g* = 1, *β* = 1, *w* = 0.5, *σ*_0_ = 2.5 *σ*_1_, 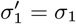. (**e**) The model parameters are chosen to be the same as in the third panel of (b) but *η* is varied. In (a)-(e) the parameter *ρ* is chosen so that 〈*n*〉_lin_ = 30.

It is still unclear how the ENM performs when mRNA synthesis is not balanced (*β* < 1). To see this, we further illustrate *D* as a function of *β* and 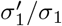 when gene expression is bursty (Fig. 5(c)). Interestingly, there is a region of parameter space (shown in dark blue) where *D* is minimised. In particular, when the mRNA synthesis rate is volume-independent (*β* = 0), the ENM works well when 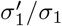 is between 0.65 and 0.8. This shows that to maintain the effectiveness of the ENM approximation, a lack of balanced mRNA synthesis requires also a lower degree of dosage compensation. Recent studies have shown that even when *β* < 1, strong concentration homeostasis (characterised by a small coefficient of variation of the mean concentrations across the cell cycle) can still be obtained when 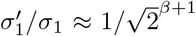 (shown by the yellow dashed line) and when replication occurs halfway through the cell cycle (*w* = 0.5) [48]. Note that the region where *D* is minimized is exactly around the yellow dashed line. This shows that the effectiveness of the ENM approximation is closely related to concentration homeostasis even when *β* < 1.

To further confirm our results, we use the transcriptional parameters inferred in [26]. In this case, the mRNA distributions for two bursty genes *Oct4* and *Nanog* in mouse embryonic stem cells were measured as a function of time in the cell cycle from which all the rate parameters involved in our model were estimated. Since the cell-to-cell variability in volume within each cell-cycle phase was quite small, it was assumed that *β* = 0, i.e. the mRNA synthesis rate is volume-independent. Dosage compensation was found to be apparent for both genes, with 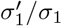 estimated to be 0.63 for *Oct4* and 0.71 for *Nanog.* Based on the inferred parameters, we compare the mRNA distributions of population measurements for the full model and the ENM (Supplementary Fig. S4(a)). We find that the ENM performs well for both genes. This agrees with our prediction that the ENM is valid when *β* = 0 and when 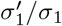 is around 0.7 (Fig. 5(c)). However, if we keep all rate parameters the same but reset 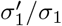 to 1 (no dosage compensation), then the EDM approximation will become significantly less satisfying (Supplementary Fig. S4(b)). This also coincides with the simulations shown in Fig. 5(c).

When mRNA synthesis is balanced and bursty, we have seen that the ENM approximation is accurate when dosage compensation is strong. However, in bacteria and budding yeast, there has been some evidence that dosage compensation is not widespread [50, 82]. It is unclear under what conditions the ENM is still valid when dosage compensation is weak. To see this, we also depict *D* as a function of *η* = *d/g* and *σ*_tot_/*g* when there is no dosage compensation, i.e. 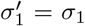 (Fig. 5(d)). In this case, we find that the ENM still works well when the mRNA is very unstable (*d* ≫ *g*) and when the total gene switching rate is very large (*σ*_tot_ ≫ *g*). This is fully consistent with our earlier theoretical predictions for the accuracy of the EDM, on which the ENM depends. In particular, when gene expression is bursty (*σ*_0_ ≫ *σ*_1_, *g*), increasing the mRNA degradation rate will give rise to a better ENM approximation (Fig. 5(e)). Note that this is not true when the total gene switching rate is slow (Fig. 5(d)). We emphasize that while Figs. 5(b),(e) show the mRNA distributions for lineage measurements, the same results are applicable for population measurements (Supplementary Fig. S5).

The value of *η* = *d/g* can be determined experimentally since both *d* and *g* can be measured. In bacteria, *η* is typically between 6 - 30, depending strongly on the strain and the growth condition; in yeast, it is typically between 3 - 8; in mammalian cells, it is typically between 2 - 4 (see Supplementary Section 4 for the median and range of *η* in various cell types). This suggests that the ENM approximation may be generally most useful in bacteria and less useful in yeast and mammalian cells.

### Including stochasticity in cell cycle duration and cell size dynamics

Thus far, we have considered a detailed telegraph model of gene expression with a cell cycle description when the cell volume dynamics and the cell cycle duration are deterministic. However, in naturally occurring systems, the cell cycle duration is appreciably stochastic (see Fig. 1(c) of [47] for experimental distributions of cell cycle durations in eight different cell types). Moreover, there has been ample evidence [33–39] that the amount of growth produced during the cell cycle must be controlled such that, on average, larger cells at birth have shorter cell cycle durations than smaller ones. This mechanism maintains size homeostasis.

To model cell-cycle duration variability and size homeostasis, we use the size-additive autoregressive model of stochastic cell volume dynamics [36, 83]. The model assumes that the volume at birth *V_b_* and the volume at division *V_d_* are connected by the relation

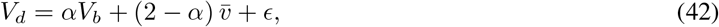

where 0 ≤ *α* ≤ 2 is the strength of size control, 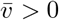 is the typical (average over generations) birth volume which is a time-independent constant, and 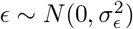 is a Gaussian noise term independent of *V_b_*. The idea behind the model is as follows: upon being born with volume *V_b_*, the cell attempts to grow for a period of time such that its target volume at division is 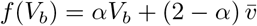, but due to stochasticity, the actual volume at division may deviate from the target volume. Due to exponential cell growth, the cell cycle duration *T* is given by

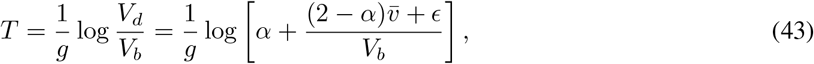

where for simplicity we have assumed constant growth rate *g* across generations. This implies that on average, larger cells at birth have shorter cell cycle durations than smaller ones. Different size control strategies correspond to different values of *α*. When *α* = 0, the target division volume 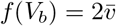 is constant; this corresponds to the sizer strategy, where cells have to reach a certain size before division occurs. When *α* = 1, the cell attempts to add a constant volume 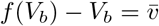 to its newborn size; this corresponds to the adder strategy. Since the growth is exponential, attempting to grow for a constant time is the same as having *f* (*V_b_*) = 2*V_b_*; hence *α* = 2 corresponds to the timer strategy. The adder or near-adder behavior has been observed in bacteria, budding yeast, and mammalian cells [35, 37, 39], while fission yeast exhibits a near-sizer behavior [33].

When *σ_ϵ_*, = 0, the model reduces to deterministic (previously considered) cell volume dynamics, in which case the timer, adder, and sizer strategies are exactly the same since 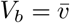 is a constant. As *σ_ϵ_*, increases, the time series of cell volume becomes much more noisy; however, it is difficult to identify whether there is a change in the magnitude of fluctuations solely from the time series of the mRNA number (Fig. 6(a)). Note that when *σ_ϵ_*, is small, the model produces a steady-state cell size distribution (from lineage simulations) characterized by three features: a fast increase in the size count for small cells, a slow decay for moderately large cells, and a fast decay for large cells (Fig. 6(b)). This is consistent with the cell size distribution in *E. coli* [70]. A natural question is what are the values of *σ_ϵ_*, in naturally occurring systems. To see this, we examined the publicly available lineage data of cell size in *E. coli* and fission yeast [36, 84] and found that the typical value of *σ_ϵ_*, is between 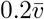 and 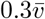 (see Supplementary Section 7 for a discussion about the inference of *σ_ϵ_*, and the estimated values of *σ_ϵ_*, in *E. coli* and fission yeast under different growth conditions).

**Figure 6:**
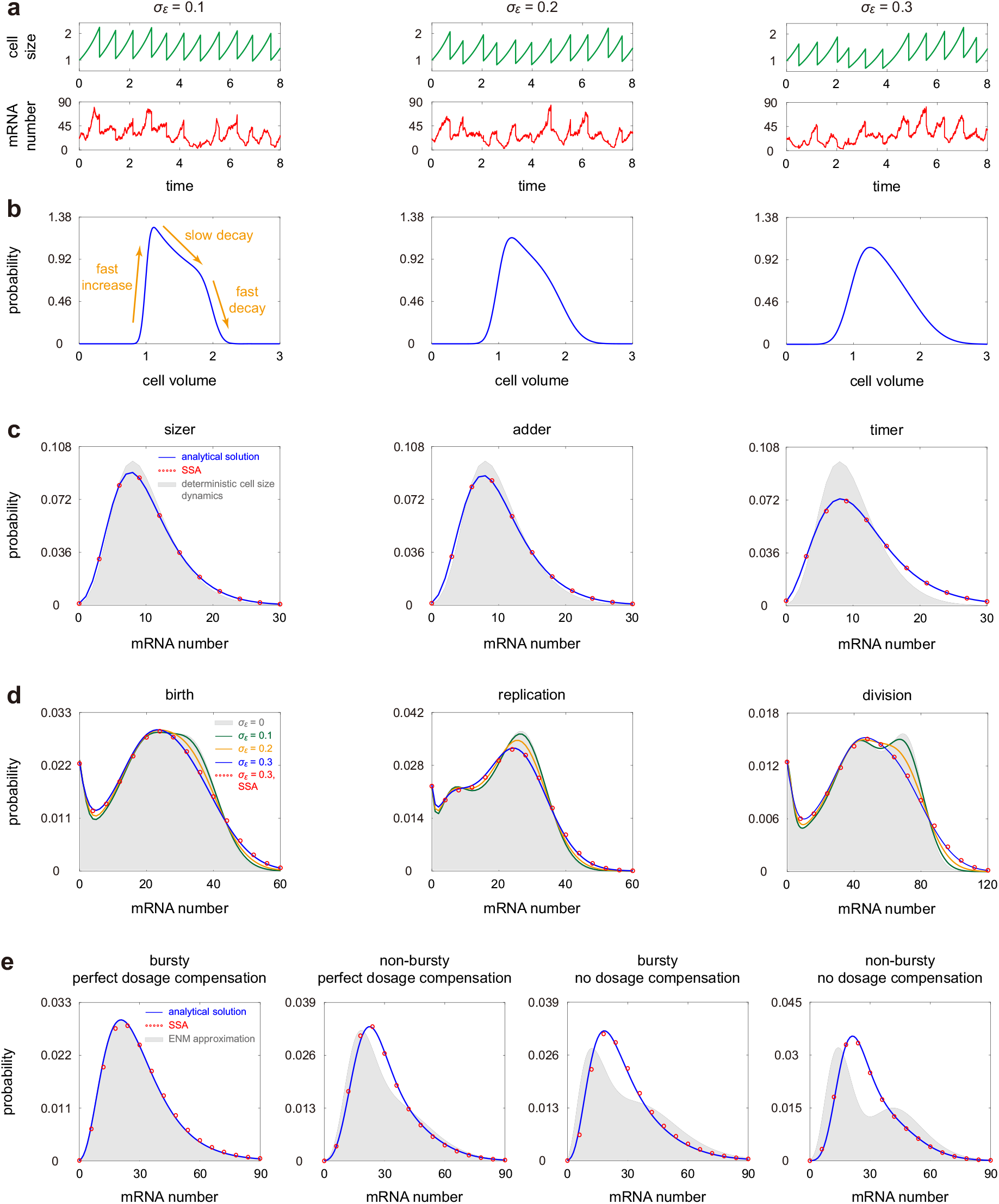
Effects of stochastic cell volume dynamics on mRNA fluctuations. (**a**) Typical trajectories of cell size and mRNA number as *σ_ϵ_* increases. (**b**) Cell volume distribution of lineage measurements as *σ_ϵ_* increases. In (a),(b), the model parameters are chosen as 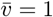, *g* = 1,*β* = 1,*w* = 0.4,*d* = 4, *ρ* = 20*d*_eff_,*σ*_0_ = 1.5, *σ*_1_ = 3, 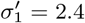, *α* =1. (**c**) Comparison between the steady-state mRNA distributions of lineage measurements for deterministic and stochastic cell size dynamics under different size control strategies. The blue curves show the analytical distributions for stochastic cell size dynamics, the red circles show the numerical ones obtained from the SSA, and the grey regions show the distributions for deterministic cell size dynamics. The model parameters are chosen as 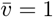, *g* = 1, *β* = 1, *w* = 0.5, *d* = 5, *σ*_0_ = *σ*_1_ = 100, 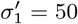, *σ_ϵ_* = 0.4. The parameter *ρ* is chosen so that 〈*n*〉_lin_ = 10 for deterministic cell size dynamics. Previous studies [85] have shown that the timer strategy with *α* = 2 is not stable since it cannot produce a finite and nonzero mean of cell volume. Hence we choose *α* = 1.8 for the timer strategy here. (**d**) Steady-state mRNA distributions at birth, replication, and division as *σ_ϵ_* increases. The model parameters are the same as in (a),(b). (**e**) Comparison between the lineage distributions of the full model and the ENM for stochastic cell size dynamics. The grey regions show the distributions for the ENM. The model parameters are chosen to be the same as in Fig. 5(b).

To compute the mRNA distribution for stochastic cell volume dynamics, note that the evolution of the system within a cell cycle is controlled by four random variables: (i) the gene state at birth *α_b_*, (ii) the mRNA number at birth *N_b_*, (iii) the birth volume *V_b_*, and (iv) the cell cycle duration *T*. Once the values of the four variables are fixed, the generating function *F* at any time *t* ∈ [0, *T*] within a cell cycle is given by Eq. (10), i.e.

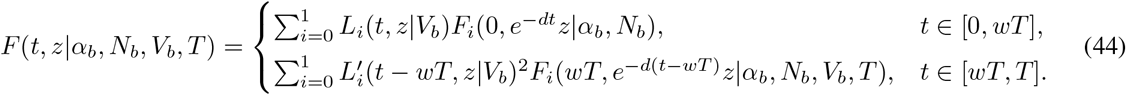

Here the initial conditions *F_i_*(0, *z*), *i* = 0,1 are determined by *α_b_* and *N_b_* as

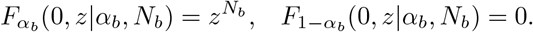

The functions *L_i_* and 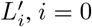 given in Eq. (9) depend on *V_b_* since the parameters *u* and *u*’ are functions of *V_b_*; the replication time *wT* depends on *T*. Hence the generating function *F* depends on all the four variables. Once we know the joint distribution of the four variables in some generation, we can use Eq. (42) to compute their joint distribution in the next generation. In this way, we obtain the full time-dependence of the mRNA distribution cross cell cycles. In Supplementary Section 7, we have generalized the analytical results obtained previously to the model with stochastic cell volume dynamics. Specifically, we have derived the exact time-dependent mRNA distribution for a cell of any age in any generation, as well as the exact steady-state distribution for lineage measurements.

To reveal the influence of cell-cycle duration variability and size homeostasis on gene expression, we compare the lineage distributions for the model with deterministic cell size dynamics and the model with stochastic cell size dynamics under different size control strategies (Fig. 6(c)). The two distributions deviate remarkably from each other for the timer strategy, but the deviation is much smaller for the adder and sizer strategies. This demonstrates the advantage of the adder and sizer strategies in reducing gene expression noise. In addition, Fig. 6(d) illustrates the steady-state mRNA distribution at three different points (birth, replication, and division) across the cell cycle as noise in cell size dynamics, characterized by *σ_ϵ_*, varies. Clearly, larger noise in cell size results in larger noise in gene expression, as expected. A sufficiently large *σ_ϵ_*, may even change the number of modes of the mRNA distribution. Interestingly, we find that when the mRNA distribution exhibits multimodality, increasing *σ_ϵ_*, will not change the height of the zero peak but may affect the height and position of non-zero peaks (Fig. 6(d)).

Finally we investigate the accuracy of the ENM approximation for stochastic cell volume dynamics. Note that we can no longer use the EDM to approximate the mRNA distributions at birth, replication, and division, since the cell volumes are stochastic. We compare the steady-state mRNA distributions at birth, replication, and division for the full model with their ENM approximations in Supplementary Fig. S6 and also compare the lineage distribution for the full model with its ENM approximation in Fig. 6(e) (see Supplementary Section 7 for the analytical expressions of the ENM approximations). The model parameters in the two figures are chosen to be the same as in Figs. 3(b),(c) and 5(b), respectively. We can see that in the presence of fluctuations in cell volume, the results of the present paper are still valid — the ENM does not work in general but performs well when mRNA synthesis is balanced and bursty and when dosage compensation is perfect. Comparing Supplementary Fig. S6 with Fig. 3(b),(c) and comparing Fig. 6(e) with Fig. 5(b), we also find that the differences between the mRNA distributions for the full model and the ENM are slightly diminished when the cell volume dynamics is stochastic.

## Discussion

In this work, we analytically solved a detailed model of stochastic gene expression with cell cycle and cell volume descriptions including gene switching, cell growth, cell division, volume-dependent transcription, gene replication, and gene dosage compensation. We first considered the case where the cell volume dynamics is deterministic and then generalized the results to include cell-cycle duration variability and cell-size control strategies. Previous models of stochastic mRNA dynamics in growing and dividing cells [22, 47] can be seen as special cases of the present modelling framework. For example, when mRNA synthesis scales with cell volume and when the gene inactivation rate is much higher than the gene activation rate, our model reduces to the one studied in [22]. Under this timescale separation assumption, there is essentially only one gene state and the computation is much easier than the one given in the present paper. If the intrinsic noise due to the random birth-death of transcripts is ignored, then our model reduces to the one-state model studied in [40]. In addition, we emphasize that our model not only characterizes the mRNA dynamics, but can also be used to describe the protein dynamics. For example, when gene expression is bursty and when the degradation rate is taken to be zero, our model reduces to the effective one-state model of the protein dynamics proposed in [41,42, 46].

Our work is also distinctive from recent related work [49] since our derivations of the distributions of mRNA numbers as a function of cell age and generation number, and of the distributions in steady-state growth do not need the assumption of stochastic concentration homeostasis (SCH); the relaxation of this assumption is crucial to model the variation of gene copy numbers across a cell cycle due to DNA replication. We have also investigated how well can the model be approximated by the effective dilution and extrinsic noise models (EDM/ENM). When gene replication is taken into account, we showed that the mRNA distributions of the full model may differ significantly from the predictions of the EDM/ENM. We elucidated three cases where the EDM/ENM makes accurate approximations.

The first case occurs when the mRNA is very unstable and the total gene switching rate (the sum of the gene activation and inactivation rates) is very large such that on the timescale of volume change, the mRNA distribution instantaneously equilibrates. This condition is intuitive and has been discussed in earlier work [57]. However as we showed using data from various cell types, the typical mRNA lifetime in eukaryotes (especially mammalian cells) is generally not small enough compared to the cell cycle duration to enforce instantaneous equilibrium; rather the fluctuations have memory of birth and replication events.

The second case takes place when mRNA synthesis is balanced and bursty, and when dosage compensation is perfect. While our model does not generally obey SCH due to gene copy number variation upon replication, however in this case parameter conditions effectively enforce SCH. Note that if expression is balanced and it is bursty with weak dosage compensation or else it is constitutive with perfect dosage compensation, there is an apparent breakdown of the EDM/ENM’s ability to accurately approximate the full model. This is since in these cases the dependence of the mean mRNA numbers with cellular volume is significantly influenced by the doubling of gene copy numbers at replication. Examples where expression is balanced but the effects of replication are not completely buffered by dosage compensation are starting to be uncovered, e.g. in human cells while the overall mRNA synthesis rates increase with cell volume, however S/G2-phase cells show increased synthesis rates compared to G1-phase cells of the same volume [86]. As pointed out in [61], this is reminiscent of a step-increase in RNA production during or after S phase which was previously observed in synchronized HeLa cell populations and other organisms [87] - this suggests that perfect dosage compensation in mammalian cells may not be common.

The third case is when mRNA synthesis is non-balanced and bursty, and when dosage compensation is of an intermediate strength such that concentration homeostasis is approximately maintained, i.e. there is only a small variation of the mean mRNA concentration throughout the cell cycle — note that this is a much weaker condition than SCH. We showed that this is indeed the case for two genes, *Oct4* and *Nanog* in mouse embryonic stem cells, whose parameters have been previously estimated before and after gene replication [26]. Recent studies [48] have shown, using both theory and data, that when gene expression is bursty, deviations from SCH show up as deviations from the gamma distribution in the mRNA concentration. This can be used to test whether SCH is approximately valid *in vivo.*

Our model is complex due to the coupling between gene expression dynamics, cell volume dynamics, and cell cycle events. A natural question is whether all the parameters involved in the model can be inferred accurately. In fact, parameter inference for models that are more complex than the telegraph model but simpler than our model has been made in our previous papers using the method of distribution matching [48] or power spectrum matching [66]. Whether accurate parameter estimation is possible by fitting mRNA distributions from population snapshot data to the analytical expression given by our calculations remains an open question.

## Limitations of the study

In summary, our work shows that caution is needed when the ENM is applied to explain data collected in growing and dividing cells and that the accuracy of this reduced model of gene expression cannot be *a priori* assumed genome-wide. Our model, though detailed, has some limitations. We have focused on models that explain cell-to-cell variability in the synthesis rates due to their dependence on cell size. However, likely other descriptors of cell state (such as shape, local cell crowding, mitochondrial abundance, capacity to respond to Ca^2+^) can explain a higher degree of cell-to-cell variability than cell size alone [88, 89]. In addition, here we have not considered the G0 phase, where cells are not growing and are outside of the replicative cell cycle. For the subpopulation of cells permanently in G0 such as senescent and many differentiated cells, since cells do not grow and divide, we can always use the ENM to characterize their gene expression dynamics.

Last but not least, here we have considered the expression of unregulated genes but it is well known that many genes regulate each other resulting in complex gene regulatory networks [90]. Overcoming the last limitation is particularly pressing but it is analytically challenging because such models have nonlinear propensities stemming from the modelling of bimolecular interactions between transcriptional factors and genes [91]. Progress in this direction will be reported in a separate paper. We also anticipate that the results of the present paper can be generalized to include more than two gene states [16–18].

## Resource availability

### Lead Contact

Further information and requests for resources should be directed to and will be fulfilled by the Lead Contact, Ramon Grima.

### Materials Availability

This study did not generate new unique reagents.

## Data and Code Availability

MATLAB codes for computing mRNA distributions using the FSP algorithm and the analytical solution can be found on GitHub https://github.com/chenjiacsrc/telegraph-model.

## Methods

### Special cases of the time-dependent mRNA distribution

Next we focus on two non-trivial special cases where the time-dependent distribution given in Eq. (11) can be greatly simplified. We assume the same setup as Fig. 2, i.e. initially there is no mRNA molecules in the cell and the gene is in the inactive state. In this case, we have *F*_0_(0, *z*) = 1 and *F*_1_(0, *z*) = 0.

The first special case occurs when the gene switches rapidly between the two states, i.e. *σ*_0_, *σ*_1_ ≫ *d, g*. In this case, we have *a, b, r* ≫ 1 and thus the confluent hypergeometric function term in Eq. (9) reduces to

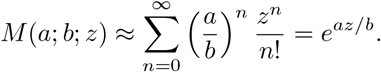

Direct computations show that the generating function given in Eq. (10) reduces to 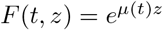,

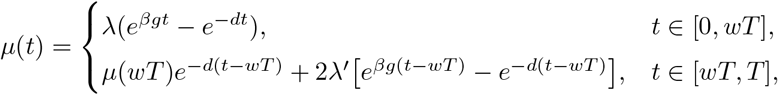

where with λ = *au/b* and *λ*′ = *a*′*u*′/*b*′. Then the time-dependent mRNA distribution is given by

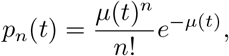

which is a Poisson distribution with mean *μ*(*t*).

The second special case occurs when the mRNA is produced in a bursty manner, i.e. *σ*_0_ ≫ *σ*_1_ and *ρ/σ*_0_ is finite [74]. In this case, the gene is mostly in the inactive state, but when it becomes active, it produces a large number of mRNA molecules. Clearly, we have *b* ≫ *a, u/b* is finite, and thus the confluent hypergeometric function terms in Eq. (9) reduce to

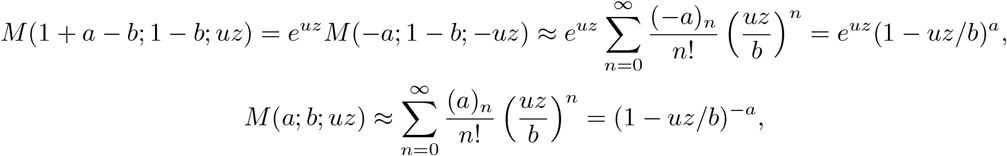

where we have used Kummer’s transformation in the first equation. Straightforward computations show that the generating function given in Eq. (10) can be simplified as

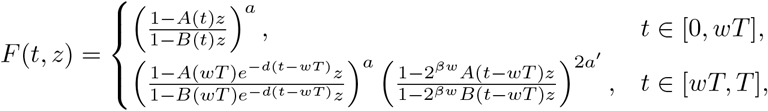

where *A*(*t*) = (*u/b*)*e*^-*dt*^ is a decay term due to mRNA degradation and *B*(*t*) = (*u/b*)*e*^*β_g_t*^ is the mean burst size at time *t* ∈ [0, *wT*]. In the bursty limit, the burst frequency for each gene copy decreases from *σ*_1_ to 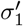 upon replication. When 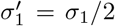, the total burst frequency does not change when replication occurs and dosage compensation is perfect. In this case, we have 2*a*′ = a and the generating function F reduces to

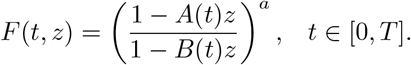

Then the time-dependent mRNA distribution is given by

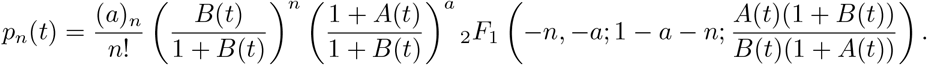

If the mRNA has a much shorter lifetime compared to the cell cycle duration, i.e. *d* ≫ *g*, then *A*(*t*) ≈ 0 and thus the mRNA number has the negative binomial distribution

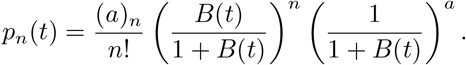

### Special cases of the steady-state mRNA distribution

In some special cases, the integrals in Eqs. (31) and (32) can be computed explicitly. The first case occurs when mRNA is unstable and the gene switches rapidly between the two states. In this case, we have *a, b* ≫ 1 and thus *M*(*a; b; z*) ≈ *e^az/b^*. It then follows from Eqs. (26) and (27) that the generating function *F^ss^* (*t, z*) at any time within a cell cycle is given by

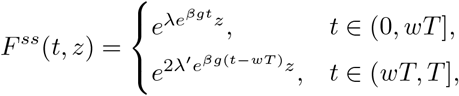

where *λ* = *au/b* and *λ*’ = *a*’*u*’/*b*’. Hence the steady-state distribution for lineage measurements can be recovered from the generating function *F*_lin_(*z*) given in Eq. (31) by taking the derivatives at *z* = −1, i.e.

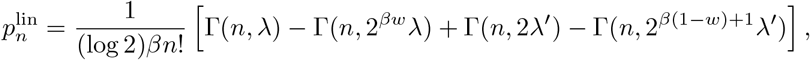

where 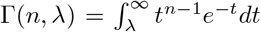 is the incomplete gamma function. Similarly, the steady-state distribution for population measurements can be recovered from the generating function *F*_pop_(*z*) given in Eq. (32) and is given by

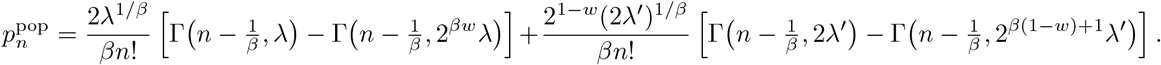

The second case occurs when mRNA is unstable and the gene switches slowly between the two states. In this case, we have *a, b* ≪ 1 and thus

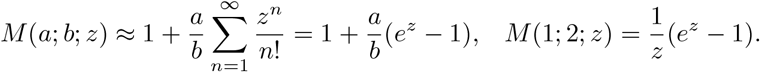

From Eqs. (26) and (27), the generating function *F^ss^* (*t, z*) at any time within a cell cycle is given by

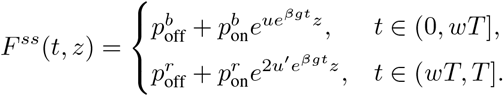

Hence it follows from Eq. (31) that the lineage distribution is given by

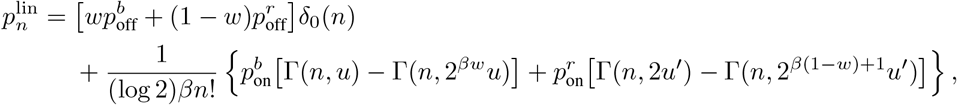

where *δ*_0_, (*n*) is the Kronecker delta which takes the value of 1 when *n* = 0 and takes the value of 0 otherwise, and it follows from Eq. (32) that the population distribution is given by

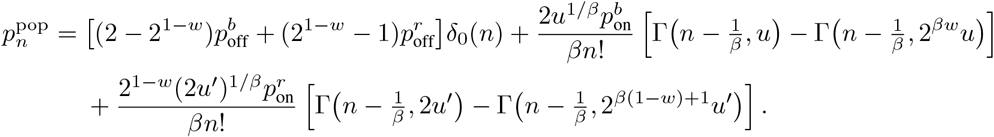

### Fluctuation relation between gene expression and cell volume

Here we give the proof of Eq. (38). For simplicity, we only focus on population measurements; the proof for lineage measurements is totally the same. Note that the Fano factor of the mRNA number in a population of cells can be decomposed as

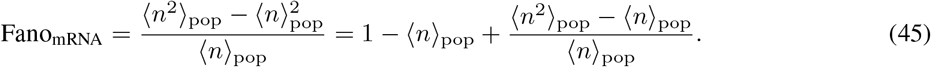

When mRNA synthesis is balanced (*β* = 1), it follows from Eqs. (35) and (37) that the mean mRNA number scales with the birth volume *V_b_* and the second factor moment scales with 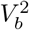, i.e.

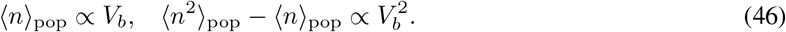

This is because the parameters *u, u*’, λ, λ’, *μ*, and *μ* are all proportional to *V_b_* when *β* = 1. On the other hand, the cell volume distribution for population measurements is given by Eq. (41). It is easy to see that the mean and variance of cell volume in a population of cells are given by

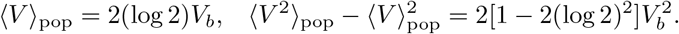

Hence the Fano factor of cell volume is given by

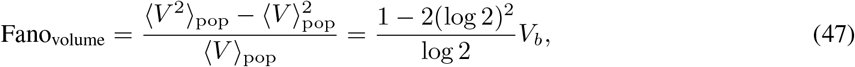

which also scales with *V_b_*. Inserting Eqs. (46) and (47) into Eq. (45), we immediately obtain Eq. (38).

### Emergence of concentration homeostasis

Here we focus on two special cases for mRNA to display concentration homeostasis. The first special case takes place where gene replication is not taken into account (*w* = 1), the gene activation and inactivation probabilities at birth are given by 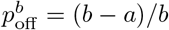 and 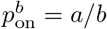. This implies that *μ* = 0 and thus the steady-state mean at birth can be simplified to a large extent as

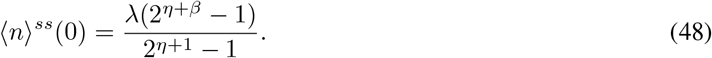

Inserting this equation into Eq. (35) yields

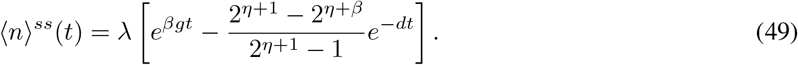

In particular, when *β* = 1, we have 〈*n*〉^*ss*^(0) = λ and 〈*n*〉^*ss*^(*t*) = λ*e*^*gt*^ = (λ/*V_b_*)*V*(*t*). In this case, the mRNA displays concentration homeostasis, i.e. constant mean concentration across the cell cycle.

Another special case occurs when the mRNA is produced in a bursty manner (*σ*_0_ ≫ *σ*_1_) and when dosage compensation is perfect 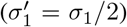. In this case the gene is mostly off and thus 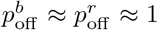. This further implies that λ’ = 2^β_w_-1^λ and *μ* = *μ*’ = 0. Inserting these equations into Eq. (36) shows that the steady-state mean at birth is still given by Eq. (48) and the time-dependent mean under cyclo-stationary conditions is still given by Eq. (49). In particular, when *β* = 1, we have 〈*n*〉^*ss*^(0) = λ and 〈*n*〉^*ss*^(*t*) = *λe*^*gt*^ = (*λ*/*V_b_*)*V*(*t*). Hence when mRNA synthesis is balanced and bursty and when dosage compensation is perfect, the mRNA also displays concentration homeostasis.

## Acknowledgements

We are grateful to Prof. Qiwen Sun and Prof. Feng Jiao for pointing us to useful references. C. J. acknowledges support from National Natural Science Foundation of China with grant No. U1930402, No. 12271020, and No. 12131005. R. G. acknowledges support from the Leverhulme Trust (RPG-2020-327).

## References

[1] Golding, I., Paulsson, J., Zawilski, S. M. & Cox, E. C. Real-time kinetics of gene activity in individual bacteria. Cell 123, 1025–1036 (2005).

[2] Raj, A. & van Oudenaarden, A. Nature, nurture, or chance: stochastic gene expression and its consequences. Cell 135, 216–226 (2008).

[3] Taniguchi, Y. et al. Quantifying E. coli proteome and transcriptome with single-molecule sensitivity in single cells. Science 329, 533–538 (2010).

[4] Swain, P. S., Elowitz, M. B. & Siggia, E. D. Intrinsic and extrinsic contributions to stochasticity in gene expression. Proceedings of the National Academy of Sciences 99, 12795–12800 (2002).

[5] Lenstra, T. L., Rodriguez, J., Chen, H. & Larson, D. R. Transcription dynamics in living cells. Annual review of biophysics 45, 25–47 (2016).

[6] Shapiro, E., Biezuner, T. & Linnarsson, S. Single-cell sequencing-based technologies will revolutionize whole-organism science. Nat. Rev. Genet. 14, 618–630 (2013).

[7] Kim, J. K. & Marioni, J. C. Inferring the kinetics of stochastic gene expression from single-cell RNA-sequencing data. Genome Biol. 14, 1–12 (2013).

[8] Vu, T. N. et al. Beta-Poisson model for single-cell RNA-seq data analyses. Bioinformatics 32, 2128–2135 (2016).

[9] Larsson, A. J. et al. Genomic encoding of transcriptional burst kinetics. Nature 565, 251–254 (2019).

[10] Chen, L., Zhu, C. & Jiao, F. A generalized moment-based method for estimating parameters of stochastic gene transcription. Math. Biosci. 345, 108780 (2022).

[11] Ko, M. S. A stochastic model for gene induction. J. Theor Biol. 153, 181–194 (1991).

[12] Peccoud, J. & Ycart, B. Markovian modeling of gene-product synthesis. Theor. Popul. Biol. 48, 222–234 (1995).

[13] Raj, A., Peskin, C. S., Tranchina, D., Vargas, D. Y. & Tyagi, S. Stochastic mRNA synthesis in mammalian cells. PLoS Biol. 4, e309 (2006).

[14] Iyer-Biswas, S., Hayot, F. & Jayaprakash, C. Stochasticity of gene products from transcriptional pulsing. Phys. Rev. E 79, 031911 (2009).

[15] Jiao, F., Tang, M. & Yu, J. Distribution profiles and their dynamic transition in stochastic gene transcription. J. Differ. Equations 254, 3307–3328 (2013).

[16] Zhou, T. & Zhang, J. Analytical results for a multistate gene model. SIAM J. Appl. Math. 72, 789–818 (2012).

[17] Chen, J. & Jiao, F. A novel approach for calculating exact forms of mRNA distribution in single-cell measurements. Mathematics 10, 27 (2021).

[18] Jia, C. & Li, Y. Analytical time-dependent distributions for gene expression models with complex promoter switching mechanisms. bioRxiv (2022).

[19] Suter, D. M. et al. Mammalian genes are transcribed with widely different bursting kinetics. Science 332, 472–474 (2011).

[20] Berg, O. G. A model for the statistical fluctuations of protein numbers in a microbial population. J. Theor. Biol. 71, 587–603 (1978).

[21] Paulsson, J., Berg, O. G. & Ehrenberg, M. Stochastic focusing: fluctuation-enhanced sensitivity of intracellular regulation. Proceedings of the National Academy of Sciences 97, 7148–7153 (2000).

[22] Cao, Z. & Grima, R. Analytical distributions for detailed models of stochastic gene expression in eukaryotic cells. Proc. Natl. Acad. Sci. USA 117, 4682–4692 (2020).

[23] Anders, S. & Huber, W. Differential expression analysis for sequence count data. Genome Biol. 11, R106 (2010).

[24] Shalek, A. K. et al. Single-cell transcriptomics reveals bimodality in expression and splicing in immune cells. Nature 498, 236–240 (2013).

[25] Singer, Z. S. et al. Dynamic heterogeneity and DNA methylation in embryonic stem cells. Molecular cell 55, 319–331 (2014).

[26] Skinner, S. O. et al. Single-cell analysis of transcription kinetics across the cell cycle. Elife 5, e12175 (2016).

[27] Huh, D. & Paulsson, J. Non-genetic heterogeneity from stochastic partitioning at cell division. Nat. Genet. 43, 95 (2011).

[28] Zhurinsky, J. et al. A coordinated global control over cellular transcription. Curr. Biol. 20 (2010).

[29] Sun, X.-M. et al. Size-dependent increase in RNA Polymerase II initiation rates mediates gene expression scaling with cell size. Curr. Biol. 30, 1217–1230 (2020).

[30] Padovan-Merhar, O. et al. Single mammalian cells compensate for differences in cellular volume and DNA copy number through independent global transcriptional mechanisms. Mol. Cell 58, 339–352 (2015).

[31] Kempe, H., Schwabe, A., Crémazy, F., Verschure, P. J. & Bruggeman, F. J. The volumes and transcript counts of single cells reveal concentration homeostasis and capture biological noise. Mol. Biol. Cell 26, 797–804 (2015).

[32] Ietswaart, R., Rosa, S., Wu, Z., Dean, C. & Howard, M. Cell-size-dependent transcription of FLC and its antisense long non-coding RNA COOLAIR explain cell-to-cell expression variation. Cell Syst. 4, 622–635 (2017).

[33] Fantes, P. & Nurse, P. Control of cell size at division in fission yeast by a growth-modulated size control over nuclear division. Exp. Cell Res. 107, 377–386 (1977).

[34] Campos, M. et al. A constant size extension drives bacterial cell size homeostasis. Cell 159, 1433–1446 (2014).

[35] Taheri-Araghi, S. et al. Cell-size control and homeostasis in bacteria. Curr. Biol. 25, 385–391 (2015).

[36] Tanouchi, Y. et al. A noisy linear map underlies oscillations in cell size and gene expression in bacteria. Nature 523, 357–360 (2015).

[37] Soifer, I., Robert, L. & Amir, A. Single-cell analysis of growth in budding yeast and bacteria reveals a common size regulation strategy. Curr. Biol. 26, 356–361 (2016).

[38] Facchetti, G., Chang, F. & Howard, M. Controlling cell size through sizer mechanisms. Curr. Opin. Syst. Biol. 5, 86–92 (2017).

[39] Cadart, C. et al. Size control in mammalian cells involves modulation of both growth rate and cell cycle duration. Nat. Commun. 9, 1–15 (2018).

[40] Antunes, D. & Singh, A. Quantifying gene expression variability arising from randomness in cell division times. J. Math. Biol. 71, 437–463 (2015).

[41] Soltani, M., Vargas-Garcia, C. A., Antunes, D. & Singh, A. Intercellular variability in protein levels from stochastic expression and noisy cell cycle processes. PLoS Comput. Biol. 12, e1004972 (2016).

[42] Soltani, M. & Singh, A. Effects of cell-cycle-dependent expression on random fluctuations in protein levels. R. Soc. Open Sci. 3, 160578 (2016).

[43] Sun, Q., Jiao, F., Lin, G., Yu, J. & Tang, M. The nonlinear dynamics and fluctuations of mRNA levels in cell cycle coupled transcription. PLoS Comput. Biol. 15, e1007017 (2019).

[44] Dessalles, R., Fromion, V. & Robert, P. Models of protein production along the cell cycle: An investigation of possible sources of noise. PLoS one 15, e0226016 (2020).

[45] Jedrak, J., Kwiatkowski, M. & Ochab-Marcinek, A. Exactly solvable model of gene expression in a proliferating bacterial cell population with stochastic protein bursts and protein partitioning. Phys. Rev. E 99, 042416 (2019).

[46] Beentjes, C. H., Perez-Carrasco, R. & Grima, R. Exact solution of stochastic gene expression models with bursting, cell cycle and replication dynamics. Phys. Rev. E 101, 032403 (2020).

[47] Perez-Carrasco, R., Beentjes, C. & Grima, R. Effects of cell cycle variability on lineage and population measurements of messenger RNA abundance. J. R. Soc. Interface 17, 20200360 (2020).

[48] Jia, C., Singh, A. & Grima, R. Concentration fluctuations in growing and dividing cells: Insights into the emergence of concentration homeostasis. PLoS Comput. Biol. 18, e1010574 (2022).

[49] Thomas, P. & Shahrezaei, V. Coordination of gene expression noise with cell size: extrinsic noise versus agent-based models of growing cell populations. J. R. Soc. Interface 18, 20210274 (2021).

[50] Wang, M., Zhang, J., Xu, H. & Golding, I. Measuring transcription at a single gene copy reveals hidden drivers of bacterial individuality. Nat. Microbiol. 4, 2118–2127 (2019).

[51] Kalita, I., Iosub, I. A., Granneman, S. & El Karoui, M. Fine-tuning of RecBCD expression by post-transcriptional regulation is required for optimal DNA repair in Escherichia coli. bioRxiv (2021).

[52] Marguerat, S. & Bähler, J. Coordinating genome expression with cell size. Trends in Genetics 28, 560–565 (2012).

[53] Neurohr, G. E. et al. Excessive cell growth causes cytoplasm dilution and contributes to senescence. cell 176, 1083–1097 (2019).

[54] Dolatabadi, S. et al. Cell cycle and cell size dependent gene expression reveals distinct subpopulations at single-cell level. Frontiers in genetics 8, 1 (2017).

[55] Swaffer, M. P. et al. Transcriptional and chromatin-based partitioning mechanisms uncouple protein scaling from cell size. Molecular Cell 81, 4861–4875 (2021).

[56] Sherman, M. S., Lorenz, K., Lanier, M. H. & Cohen, B. A. Cell-to-cell variability in the propensity to transcribe explains correlated fluctuations in gene expression. Cell systems 1, 315–325 (2015).

[57] Ham, L., Brackston, R. D. & Stumpf, M. P. Extrinsic noise and heavy-tailed laws in gene expression. Phys. Rev. Lett. 124, 108101 (2020).

[58] Ham, L., Jackson, M. & Stumpf, M. P. Pathway dynamics can delineate the sources of transcriptional noise in gene expression. Elife 10, e69324 (2021).

[59] Wang, P. et al. Robust growth of Escherichia coli. Curr. Biol. 20, 1099–1103 (2010).

[60] Eun, Y.-J. et al. Archaeal cells share common size control with bacteria despite noisier growth and division. Nat. Microbiol. 3, 148–154 (2018).

[61] Berry, S. & Pelkmans, L. Mechanisms of cellular mRNA transcript homeostasis. Trends in Cell Biology (2022).

[62] Wang, Q. & Lin, J. Heterogeneous recruitment abilities to RNA polymerases generate nonlinear scaling of gene expression with cell volume. Nature communications 12, 1–11 (2021).

[63] Dowling, M. R. et al. Stretched cell cycle model for proliferating lymphocytes. Proc. Natl. Acad. Sci. USA 111, 6377–6382 (2014).

[64] Deng, Q., Ramsköld, D., Reinius, B. & Sandberg, R. Single-cell RNA-seq reveals dynamic, random monoallelic gene expression in mammalian cells. Science 343, 193–196 (2014).

[65] Sepúlveda, L. A., Xu, H., Zhang, J., Wang, M. & Golding, I. Measurement of gene regulation in individual cells reveals rapid switching between promoter states. Science 351, 1218–1222 (2016).

[66] Jia, C. & Grima, R. Frequency domain analysis of fluctuations of mRNA and protein copy numbers within a cell lineage: theory and experimental validation. Phys. Rev. X 11, 021032 (2021).

[67] Nicolas, D., Phillips, N. E. & Naef, F. What shapes eukaryotic transcriptional bursting? Molecular BioSystems 13, 1280–1290 (2017).

[68] Reverón-Gómez, N. et al. Accurate recycling of parental histones reproduces the histone modification landscape during DNA replication. Mol. Cell 72, 239–249 (2018).

[69] Voichek, Y., Bar-Ziv, R. & Barkai, N. Expression homeostasis during DNA replication. Science 351, 1087–1090 (2016).

[70] Jia, C., Singh, A. & Grima, R. Cell size distribution of lineage data: analytic results and parameter inference. iScience 24, 102220 (2021).

[71] Jia, C., Singh, A. & Grima, R. Characterizing non-exponential growth and bimodal cell size distributions in fission yeast: An analytical approach. PLoS Comput. Biol. 18, e1009793 (2022).

[72] Paulsson, J. Models of stochastic gene expression. Phys. Life Rev. 2, 157–175 (2005).

[73] Jia, C., Zhang, M. Q. & Hong, Q. Emergent Lévy behavior in single-cell stochastic gene expression. Phys. Rev. E 96, 040402(R) (2017).

[74] Jia, C. Simplification of Markov chains with infinite state space and the mathematical theory of random gene expression bursts. Phys. Rev. E 96, 032402 (2017).

[75] Jiao, F. & Tang, M. Quantification of transcription noise¡^-^s impact on cell fate commitment with digital resolutions. Bioinformatics 38, 3062–3069 (2022).

[76] Munsky, B. & Khammash, M. The finite state projection algorithm for the solution of the chemical master equation. J. Chem. Phys. 124, 044104 (2006).

[77] Friedman, N., Cai, L. & Xie, X. S. Linking stochastic dynamics to population distribution: an analytical framework of gene expression. Phys. Rev. Lett. 97, 168302 (2006).

[78] Jia, C., Qian, H., Chen, M. & Zhang, M. Q. Relaxation rates of gene expression kinetics reveal the feedback signs of autoregulatory gene networks. J. Chem. Phys. 148, 095102 (2018).

[79] Pierson, E. & Yau, C. ZIFA: Dimensionality reduction for zero-inflated single-cell gene expression analysis. Genome Biol. 16, 241 (2015).

[80] Risso, D., Perraudeau, F., Gribkova, S., Dudoit, S. & Vert, J.-P. A general and flexible method for signal extraction from single-cell RNA-seq data. Nat. Commun. 9, 284 (2018).

[81] Jia, C. Kinetic foundation of the zero-inflated negative binomial model for single-cell RNA sequencing data. SIAM J. Appl. Math. 80, 1336–1355 (2020).

[82] Torres, E. M., Springer, M. & Amon, A. No current evidence for widespread dosage compensation in S. cerevisiae. Elife 5, e10996 (2016).

[83] Amir, A. Cell size regulation in bacteria. Phys. Rev. Lett. 112, 208102 (2014).

[84] Nakaoka, H. & Wakamoto, Y. Aging, mortality, and the fast growth trade-off of Schizosaccharomyces pombe. PLoS Biol. 15, e2001109 (2017).

[85] Vargas-Garcia, C. A., Soltani, M. & Singh, A. Conditions for cell size homeostasis: a stochastic hybrid system approach. IEEE Life Sci. Lett. 2, 47–50 (2016).

[86] Berry, S., Müller, M., Rai, A. & Pelkmans, L. Feedback from nuclear RNA on transcription promotes robust RNA concentration homeostasis in human cells. Cell Systems (2022).

[87] Mitchison, J. Growth during the cell cycle. International review of cytology 166–258 (2003).

[88] Battich, N., Stoeger, T. & Pelkmans, L. Control of transcript variability in single mammalian cells. Cell 163, 1596–1610 (2015).

[89] Foreman, R. & Wollman, R. Mammalian gene expression variability is explained by underlying cell state. Molecular systems biology 16, e9146 (2020).

[90] Hasty, J., McMillen, D., Isaacs, F. & Collins, J. J. Computational studies of gene regulatory networks: in numero molecular biology. Nature Reviews Genetics 2, 268–279 (2001).

[91] Cao, Z. & Grima, R. Linear mapping approximation of gene regulatory networks with stochastic dynamics. Nat. Commun. 9, 3305 (2018).

