## Supplementary Information for "Coupling gene expression dynamics to cell size dynamics and cell cycle events: exact and approximate solutions of the extended telegraph model"

### Supplementary Material —
Accuracy and limitations of extrinsic noise models to describe gene expression in growing cells

Chen Jia    Ramon Grima

#### Contents

**1 Derivation of the generating functions before replication 2**
1.1 Derivation of the generating function $F$ before replication . . . . . . . . . . . . . . . . . . 2
1.2 Derivation of the generating functions $F_0$ and $F_1$ before replication . . . . . . . . . . . . 4

**2 Derivation of the generating functions after replication 6**
2.1 Derivation of the generating function $F$ after replication . . . . . . . . . . . . . . . . . . 6
2.2 Derivation of the generating functions $F_0$ and $F_1$ after replication . . . . . . . . . . . . . 9

**3 Modified FSP algorithm 10**

**4 Typical values of $\eta$ in various cell types 10**

**5 Further moment analysis for lineage and population measurements 11**

**6 Comparison with the conventional telegraph model 12**

**7 Generalization of the theory to stochastic cell volume dynamics 13**

### 1 Derivation of the generating functions before replication

#### 1.1 Derivation of the generating function $F$ before replication

Here we derive the analytical expression of the generating function $F$ before replication. In the main text, we have shown that in terms of the generating functions, the master equation can be converted into the PDEs

$$\begin{aligned}\partial_t F_0 &= -dz\partial_z F_0 + \sigma_0 F_1 - \sigma_1 F_0,\\ \partial_t F_1 &= s(t)zF_1 - dz\partial_z F_1 + \sigma_1 F_0 - \sigma_0 F_1,\end{aligned} \tag{1}$$

where $s(t) = \rho V(t)^\beta$. Adding the two identities in Eq. (1) shows that $F_1$ can be represented by $F$ as

$$F_1 = \frac{\partial_t F + dz\partial_z F}{s(t)z}. \tag{2}$$

Inserting this equation into the second equation of (1) shows that $F$ satisfies the second-order parabolic PDE

$$\partial_{tt}F + 2dz\partial_{tz}F + d^2z^2\partial_{zz}F + [r - d - s(t)z]\partial_t F + dz[r - s(t)z]\partial_z F - \sigma_1 s(t)zF = 0, \tag{3}$$

where $r = \sigma_0 + \sigma_1 - \beta g$. Following [1], we introduce a new variable $\tau = \log z - dt$. Let $\tilde{F}(\tau, z)$ and $\tilde{F}_i(\tau, z)$ be the functions with variables $\tau$ and $z$ that are associated with $F(t, z)$ and $F_i(t, z)$, respectively, i.e.

$$F(t, z) = \tilde{F}(\log z - dt, z), \quad F_i(t, z) = \tilde{F}_i(\log z - dt, z), \quad i = 0, 1.$$

Then Eq. (3) can be simplified to a large extent as

$$d^2 z\partial_{zz}\tilde{F} + d[r - s(t)z]\partial_z\tilde{F} - \sigma_1 s(t)\tilde{F} = 0. \tag{4}$$

If we fix the variable $\tau$, this is an ordinary differential equation (ODE) with respect to the variable $z$. Note that

$$s(t) = \rho V_b^\beta e^{\beta g t} = \rho V_b^\beta e^{\beta g(\log z - \tau)/d} = \rho V_b^\beta e^{-(\beta g/d)\tau} z^{\beta g/d}. \tag{5}$$

Inserting this equation into Eq. (4) yields

$$d^2 z\partial_{zz}\tilde{F} + d[r - \rho V_b^\beta e^{-(\beta g/d)\tau} z^{\beta g/d+1}]\partial_z\tilde{F} - \sigma_1\rho V_b^\beta e^{-(\beta g/d)\tau} z^{\beta g/d}\tilde{F} = 0.$$

This is a modified version of the confluent hypergeometric differential function and its solution can be written in general form as

$$\tilde{F}(\tau, z) = \phi_0(e^\tau)H_0(\tau, z) + \phi_1(e^\tau)H_1(\tau, z), \tag{6}$$

where

$$\begin{aligned}H_0(\tau, z) &= M(a; b; ue^{-(\beta g/d)\tau} z^{\beta g/d+1}),\\ H_1(\tau, z) &= z^{1-r/d} M(1 + a - b; 2 - b; ue^{-(\beta g/d)\tau} z^{\beta g/d+1}).\end{aligned} \tag{7}$$

with $a = \sigma_1/(d + \beta g)$, $b = (\sigma_0 + \sigma_1)/(d + \beta g)$, $u = \rho V_b^\beta/(d + \beta g)$, and with $M(a; b; x)$ being the confluent hypergeometric function.

The remaining question is how to determine the functions $\phi_0$ and $\phi_1$ based on the initial conditions. By the definition of $\tilde{F}_i$, it is easy to see that

$$\tilde{F}_i(\log z, z) = F_i(0, z), \quad i = 0, 1.$$

Taking $\tau = \log z$ in Eq. (6) yields

$$\phi_0(z)I_0(z) + \phi_1(z)I_1(z) = \tilde{F}(\log z, z) = F(0, z), \tag{8}$$

where

$$I_0(z) = H_0(\log z, z) = M(a; b; uz),$$
$$I_1(z) = H_1(\log z, z) = z^{1-r/d} M(1 + a - b; 2 - b; uz).$$

Moreover, it follows from Eq. (2) that

$$d\partial_z \tilde{F}(\tau, z) = s(t)\tilde{F}_1(\tau, z) = \rho V_b^\beta e^{-(\beta g/d)\tau} z^{\beta g/d} \tilde{F}_1(\tau, z).$$

This shows that

$$d\partial_z \tilde{F}(\log z, z) = \rho V_b^\beta \tilde{F}_1(\log z, z) = u(d + \beta g) F_1(0, z).$$

It follows from Eq. (6) that

$$\partial_z \tilde{F}(\log z, z) = \phi_0(z)\partial_z H_0(\log z, z) + \phi_1(z)\partial_z H_1(\log z, z).$$

Combining the above two equations yields

$$\phi_0(z)J_0(z) + \phi_1(z)J_1(z) = u(\beta g/d + 1)F_1(0, z), \tag{9}$$

where

$$J_0(z) = \partial_z H_0(\log z, z) = \frac{au(\beta g/d + 1)}{b} M(1 + a; 1 + b; uz),$$
$$J_1(z) = \partial_z H_1(\log z, z) = (1 - b)(\beta g/d + 1) z^{-r/d} M(1 + a - b; 1 - b; uz).$$

Combining Eqs. (8) and (9), we obtain

$$\begin{pmatrix} I_0(z) & I_1(z) \\ J_0(z) & J_1(z) \end{pmatrix} \begin{pmatrix} \phi_0(z) \\ \phi_1(z) \end{pmatrix} = \begin{pmatrix} F(0, z) \\ u(\beta g/d + 1)F_1(0, z) \end{pmatrix}.$$

This shows that

$$\begin{aligned} \phi_0(z) &= \frac{J_1(z)F_0(0, z) + [J_1(z) - u(\beta g/d + 1)I_1(z)]F_1(0, z)}{I_0(z)J_1(z) - I_1(z)J_0(z)} \\ \phi_1(z) &= \frac{[u(\beta g/d + 1)I_0(z) - J_0(z)]F_1(0, z) - J_0(z)F_0(0, z)}{I_0(z)J_1(z) - I_1(z)J_0(z)}. \end{aligned} \tag{10}$$

By means of the Wronskian of confluent hypergeometric functions, it is easy to check that

$$I_0(z)J_1(z) - I_1(z)J_0(z) = (1 - b)(\beta g/d + 1) z^{-r/d} e^{uz}.$$

Moreover, straightforward computations show that

$$J_1(z) - u(\beta g/d + 1)I_1(z) = (1 - b)(\beta g/d + 1) z^{-r/d} M(a - b; 1 - b; uz),$$
$$u(\beta g/d + 1)I_0(z) - J_0(z) = \frac{(b - a)u(\beta g/d + 1)}{b} M(a; 1 + b; uz).$$

Inserting the above two equations into Eq. (10), we obtain

$$\begin{aligned} \phi_0(z) &= e^{-uz}[M(1 + a - b; 1 - b; uz)F_0(0, z) + M(a - b; 1 - b; uz)F_1(0, z)], \\ \phi_1(z) &= \frac{u}{b(b - 1)} z^{r/d} e^{-uz}[aM(1 + a; 1 + b; uz)F_0(0, z) - (b - a)M(a; 1 + b; uz)]F_1(0, z)]. \end{aligned} \tag{11}$$

In terms of the original variables $t$ and $z$, it follows from Eq. (6) that the generating function $F$ is given by

$$F(t, z) = \tilde{F}(\log z - dt, z) = \phi_0(e^{-dt}z)H_0(\log z - dt, z) + \phi_1(e^{-dt}z)H_1(\log z - dt, z).$$

Interesting Eqs. (7) and (11) into the above equation, we finally obtain the analytical expression of the generating function $F$, which is given by

$$F(t,z) = L_0(t,z)F_0(0,e^{-dt}z) + L_1(t,z)F_1(0,e^{-dt}z). \tag{12}$$

Here $F_0(0,z)$ and $F_1(0,z)$ are the generating functions at $t = 0$ which can be determined by the initial conditions, and the functions $L_0$ and $L_1$ are given by

$$\begin{aligned} L_0(t,z) &= \Big[M(1+a-b;1-b;ue^{-dt}z)M(a;b;ue^{\beta gt}z) + \frac{auz}{b(b-1)}e^{-rt} \\ &\quad \times M(1+a;1+b;ue^{-dt}z)M(1+a-b;2-b;ue^{\beta gt}z)\Big]e^{-ue^{-dt}z}, \\ L_1(t,z) &= \Big[M(a-b;1-b;ue^{-dt}z)M(a;b;ue^{\beta gt}z) - \frac{(b-a)uz}{b(b-1)}e^{-rt} \\ &\quad \times M(a;1+b;ue^{-dt}z)M(1+a-b;2-b;ue^{\beta gt}z)\Big]e^{-ue^{-dt}z}. \end{aligned} \tag{13}$$

#### 1.2 Derivation of the generating functions $F_0$ and $F_1$ before replication

Here we derive the analytical expression of the generating functions $F_0$ and $F_1$ before replication. From the first equation of (1), $F_1$ can be represented by $F_0$ as

$$F_1 = \frac{1}{\sigma_0}(\partial_t F_0 + dz\partial_z F_0 + \sigma_1 F_0). \tag{14}$$

Inserting into the second equation of (1) shows that $F_0$ satisfies the second-order parabolic PDE

$$\partial_{tt}F_0 + 2dz\partial_{tz}F_0 + d^2z^2\partial_{zz}F_0 + [R-d-s(t)z]\partial_t F_0 + dz[R-s(t)z]\partial_z F_0 - \sigma_1 s(t)zF_0 = 0, \tag{15}$$

where $R = \sigma_0 + \sigma_1 + d$. Using the new variables $\tau$ and $z$, Eq. (15) can be simplified to

$$d^2z\partial_{zz}\tilde{F}_0 + d[R-s(t)z]\partial_z\tilde{F}_0 - \sigma_1 s(t)\tilde{F}_0 = 0. \tag{16}$$

If we fix the variable $\tau$, this is exactly an ODE with respect to the variable $z$. Inserting Eq. (5) into Eq. (16) yields

$$z\partial_{zz}\tilde{F}_0 + [R - \rho e^{-\beta gu}z^{\beta g+1}]\partial_z\tilde{F}_0 - \sigma_1\rho e^{-\beta gu}z^{\beta g}\tilde{F}_0 = 0.$$

This is a modified version of the confluent hypergeometric differential function and its solution can be written in general form as

$$\tilde{F}_0(\tau,z) = \phi_0(e^\tau)H_0(\tau,z) + \phi_1(e^\tau)H_1(\tau,z), \tag{17}$$

where

$$\begin{aligned} H_0(\tau,z) &= M(a;1+b;ue^{-(\beta g/d)\tau}z^{\beta g/d+1}), \\ H_1(\tau,z) &= z^{1-R/d}M(a-b;1-b;ue^{-(\beta g/d)\tau}z^{\beta g/d+1}). \end{aligned} \tag{18}$$

The remaining question is how to determine the functions $\phi_0$ and $\phi_1$ based on the initial conditions. By the definition of $\tilde{F}_i$, it is easy to see that

$$\tilde{F}_i(\log z, z) = F_i(0,z), \quad i = 0,1.$$

Taking $\tau = \log z$ in Eq. (17) yields

$$\phi_0(z)I_0(z) + \phi_1(z)I_1(z) = \tilde{F}_0(\log z, z) = F_0(0,z), \tag{19}$$

where

$$\begin{aligned}
I_0(z) &= H_0(\log z, z) = M(a; 1+b; uz), \\
I_1(z) &= H_1(\log z, z) = z^{1-R} M(a-b; 1-b; uz).
\end{aligned}$$

Moreover, it follows from Eq. (14) that

$$\partial_z \tilde{F}_0 = \frac{1}{dz}[\sigma_0 \tilde{F}_1 - \sigma_1 \tilde{F}_0] = \frac{\beta g/d + 1}{z}[(b-a)\tilde{F}_1 - a\tilde{F}_0].$$

This shows that

$$\partial_z \tilde{F}_0(\log z, z) = \frac{\beta g/d + 1}{z}[(b-a)F_1(0,z) - aF_0(0,z)].$$

It follows from Eq. (17) that

$$\partial_z \tilde{F}_0(\log z, z) = \phi_0(z)\partial_z H_0(\log z, z) + \phi_1(z)\partial_z H_1(\log z, z).$$

Combining the above two equations yields

$$\phi_0(z)J_0(z) + \phi_1(z)J_1(z) = \frac{\beta g/d + 1}{z}[(b-a)F_1(0,z) - aF_0(0,z)], \tag{20}$$

where

$$\begin{aligned}
J_0(z) &= \partial_z H_0(\log z, z) = \frac{au(\beta g/d + 1)}{1+b} M(1+a; 2+b; uz), \\
J_1(z) &= \partial_z H_1(\log z, z) = -b(\beta g/d + 1) z^{-R/d} M(a-b; -b; uz).
\end{aligned}$$

Combining Eqs. (19) and (20), we obtain

$$\begin{pmatrix} I_0(z) & I_1(z) \\ J_0(z) & J_1(z) \end{pmatrix} \begin{pmatrix} \phi_0(z) \\ \phi_1(z) \end{pmatrix} = \begin{pmatrix} F_0(0,z) \\ \frac{\beta g/d+1}{z}[(b-a)F_1(0,z) - aF_0(0,z)] \end{pmatrix}.$$

This shows that

$$\begin{aligned}
\phi_0(z) &= \frac{[zJ_1(z) + (\beta g/d + 1)aI_1(z)]F_0(0,z) - (\beta g/d + 1)(b-a)I_1(z)F_1(0,z)}{z[I_0(z)J_1(z) - I_1(z)J_0(z)]} \\
\phi_1(z) &= \frac{(\beta g/d + 1)(b-a)I_0(z)F_1(0,z) - [zJ_0(z) + (\beta g/d + 1)aI_0(z)]F_0(0,z)}{z[I_0(z)J_1(z) - I_1(z)J_0(z)]}.
\end{aligned} \tag{21}$$

By means of the Wronskian of confluent hypergeometric functions, it is easy to check that

$$I_0(z)J_1(z) - I_1(z)J_0(z) = -b(\beta g/d + 1)z^{-R/d}e^{uz}.$$

Moreover, straightforward computations show that

$$\begin{aligned}
zJ_1(z) + (\beta g + 1)aI_1(z) &= -(b-a)(\beta g/d + 1)z^{1-R/d}M(1+a-b; 1-b; uz), \\
zJ_0(z) + (\beta g + 1)aI_0(z) &= a(\beta g/d + 1)M(1+a; 1+b; uz).
\end{aligned}$$

Inserting the above two equations into (21), we obtain

$$\begin{aligned}
\phi_0(z) &= \frac{b-a}{b}e^{-uz}[M(1+a-b; 1-b; uz)F_0(0,z) + M(a-b; 1-b; uz)F_1(0,z)], \\
\phi_1(z) &= \frac{1}{b}z^{R/d-1}e^{-uz}[aM(1+a; 1+b; uz)F_0(0,z) - (b-a)M(a; 1+b; uz)]F_1(0,z)].
\end{aligned} \tag{22}$$

In terms of the original variables $t$ and $z$, it follows from Eq. (17) that the generating function $F_0$ is given by

$$F_0(t,z) = \tilde{F}_0(\log z - t, z)e^{vz(e^{\beta g t} - e^{-dt})} = \phi_0(e^{-dt}z)H_0(\log z, z) + \phi_1(e^{-dt}z)H_1(\log z, z),$$

Inserting Eqs. (18) and (22) into the above equation, we finally obtain the analytical expression of the generating function $F_0$, which is given by

$$F_0(t,z) = K_{00}(t,z)F_0(0,e^{-dt}z) + K_{01}(t,z)F_1(0,e^{-dt}z). \tag{23}$$

Here $F_0(0,z)$ and $F_1(0,z)$ are the generating functions at $t=0$ which can be determined by the initial conditions, and the functions $K_{00}$ and $K_{01}$ are given by

$$\begin{aligned} K_{00}(t,z,V_b) = \frac{b-a}{b}\Big[&M(1+a-b;1-b;ue^{-dt}z)M(a;1+b;ue^{\beta gt}z) + \frac{a}{b-a}e^{-(r+\beta g)t} \\ &\times M(1+a;1+b;ue^{-dt}z)M(a-b;1-b;ue^{\beta gt}z)\Big]e^{-ue^{-dt}z}, \\ K_{01}(t,z,V_b) = \frac{b-a}{b}[&M(a-b;1-b;ue^{-dt}z)M(a;1+b;ue^{\beta gt}z) - e^{-(r+\beta g)t} \\ &\times M(a;1+b;ue^{-dt}z)M(a-b;1-b;ue^{\beta gt}z)]e^{-ue^{-dt}z}, \end{aligned}$$

where we have used the fact that $R = r + d + \beta g$. Since we have derived both $F_0$ and $F$, we finally obtain the explicit expression of $F_1 = F - F_0$, which is given by

$$F_1(t,z) = K_{10}(t,z)F_0(0,e^{-dt}z) + K_{11}(t,z)F_1(0,e^{-dt}z), \tag{24}$$

where

$$\begin{aligned} K_{10}(t,z,V_b) = \frac{a}{b}[&M(1+a-b;1-b;ue^{-dt}z)M(1+a;1+b;ue^{\beta gt}z) - e^{-(r+\beta g)t} \\ &\times M(1+a;1+b;ue^{-dt}z)M(1+a-b;1-b;ue^{\beta gt}z)]e^{-ue^{-dt}z}, \\ K_{11}(t,z,V_b) = \frac{a}{b}\Big[&M(a-b;1-b;ue^{-dt}z)M(1+a;1+b;ue^{\beta gt}z) + \frac{b-a}{a}e^{-(r+\beta g)t} \\ &\times M(a;1+b;ue^{-dt}z)M(1+a-b;1-b;ue^{\beta gt}z)]e^{-ue^{-dt}z}. \end{aligned}$$

### 2 Derivation of the generating functions after replication

#### 2.1 Derivation of the generating function $F$ after replication

We next focus on the dynamics after replication for haploid cells. Since there are two daughter gene copies after replication, to distinguish them, we call them daughter copy $A$ and daughter copy $B$. For convenience, we assume that all mRNA molecules are allocated to daughter copy $A$ when replication occurs, while no molecules are allocated to daughter copy $B$. Similarly, the microstate of each daughter copy can be described by an ordered pair $(i,n)$, where $i = 0,1$ denotes the gene state and $n$ denotes the number of mRNA molecules belonging to that daughter copy, i.e. the molecules allocated to that daughter copy at replication and the molecules produced by that daughter copy after replication. The total number of molecules after replication is then the sum of the numbers of molecules that belong to the two daughter copies. Note that other choices for how the mRNA molecules are allocated to each of the gene copies have no effect on the calculation of the statistics of the total number of molecules after replication.

Let $p^A_{i,n}(t)$ denote the probability of having $n$ transcripts that belong to daughter copy $A$ at time $t \in [wT,T]$ when daughter copy $A$ is in state $i$. Similarly, let $p^B_{i,n}(t)$ denote the same quantity for daughter copy $B$. Recall that the gene activation rate decreases from $\sigma_1$ to $\sigma_1'$ upon replication. Then the stochastic gene expression dynamics for each gene copy after replication is governed by the CMEs

$$\begin{aligned} \dot{p}^l_{0,n} &= d[(n+1)p^l_{0,n+1} - np^l_{0,n}] + [\sigma_0 p^l_{1,n} - \sigma_1' p^l_{0,n}], \\ \dot{p}^l_{1,n} &= \rho V(t)^\beta[p^l_{1,n-1} - p^l_{1,n}] + d[(n+1)p^l_{1,n+1} - np^l_{1,n}] + [\sigma_1' p^l_{0,n} - \sigma_0 p^l_{1,n}], \end{aligned} \tag{25}$$

where $l = A, B$ indicates which daughter copy is considered. To solve these, for each $l = A, B$, we define a pair of generating functions

$$F_i^l(t,z) = \sum_{n=0}^{\infty} p_{i,n}^l(t)(z+1)^n, \quad i = 0, 1. \tag{26}$$

In addition, let $p_n^l(t) = p_{0,n}^l(t) + p_{1,n}^l(t)$ denote the probability of having $n$ transcripts that belong to daughter copy $l$ at time $t$ and let $F^l(t,z) = F_0^l(t,z) + F_1^l(t,z)$ be the corresponding generating function. Then Eq. (25) can be converted into the PDEs

$$\begin{aligned} \partial_t F_0^l &= -dz\partial_z F_0^l + \sigma_0 F_1^l - \sigma_1' F_0^l, \\ \partial_t F_1^l &= s(t)zF_1^l - dz\partial_z F_1^l + \sigma_1' F_0^l - \sigma_0 F_1^l. \end{aligned} \tag{27}$$

For any $t \in [0, wT]$, let $\alpha(t)$ denote the state of the mother copy and let $X(t)$ denote the number of mRNA molecules at time $t$. For any $t \in [wT, T]$ and $j = A, B$, let $\alpha^j(t)$ denote the state of daughter copy $j$ and let $X^j(t)$ denote the number of mRNA molecules that belong to daughter copy $j$ at time $t$. Since the two daughter copies inherit the gene state of the mother copy at replication, we have

$$\alpha^A(wT) = \alpha^B(wT) = \alpha(wT).$$

Since all molecules are allocated to daughter copy $A$ and no molecules are allocated to daughter copy $B$ at replication, we have

$$X^A(wT) = X(wT), \quad X^B(wT) = 0.$$

Since $\alpha^A(wT) = \alpha^B(wT)$, the initial distributions for the two daughter copies are correlated and thus the dynamics for the two daughter copies after replication are not independent of each other. However, once the gene state of the mother copy at replication is fixed (conditioned on $\alpha(wT) = k$ with $k = 0, 1$), the initial distributions for the two daughter copies are (conditionally) independent of each other, and hence the dynamics for the two daughter copies after replication are also (conditionally) independent of each other. We now use the conditional independence of the two daughter copies to compute the generating function $F$ after replication.

Recall that $F(t,z)$ is the generating function of

$$p_n(t) = \mathbb{P}(X^A(t) + X^B(t) = n),$$

which denotes the probability of having $n$ mRNA molecules in the cell at time $t$. Moreover, recall that $F^A(t,z)$ is the generating function of

$$p_n^A(t) = \mathbb{P}(X^A(t) = n),$$

which denotes the probability of having $n$ mRNA molecules that belong to daughter copy $A$ at time $t$. In addition, recall that $F^B(t,z)$ is the generating function of

$$p_n^B(t) = \mathbb{P}(X^B(t) = n),$$

which denotes the probability of having $n$ mRNA molecules that belong to daughter copy $B$ at time $t$. Using the probabilistic notation, the generating functions $F(t,z)$, $F^A(t,z)$, and $F^B(t,z)$ can be represented by

$$F(t,z) = \mathbb{E}(z+1)^{X_A(t)+X_B(t)}, \quad F^A(t,z) = \mathbb{E}(z+1)^{X_A(t)}, \quad F^B(t,z) = \mathbb{E}(z+1)^{X_B(t)},$$

where $1_A$ denotes the indicator function of the set $A$.

We now make a crucial observation that conditioned on $\alpha(wT) = j$, i.e. the gene state of the mother copy is $j$ at replication, the dynamics $\{\alpha^A(t), X^A(t)\}_{wT \leq t \leq T}$ for daughter copy $A$ and the dynamics $\{\alpha^B(t), X^B(t)\}_{wT \leq t \leq T}$

for daughter copy $B$ are independent of each other. This shows that

$$\begin{aligned}F(t,z) &= \mathbb{E}(z+1)^{X_A(t)+X_B(t)} \\ &= \sum_{k=0}^{1} \mathbb{E}[(z+1)^{X_A(t)+X_B(t)}|\alpha(wT)=k]\mathbb{P}(\alpha(wT)=k) \\ &= \sum_{k=0}^{1} \mathbb{E}[(z+1)^{X_A(t)}|\alpha(wT)=k]\mathbb{E}[(z+1)^{X_B(t)}|\alpha(wT)=k]F_k(wT,0),\end{aligned} \tag{28}$$

where we have used the fact that $F_k(wT,0) = \mathbb{P}(\alpha(wT)=k)$. Note that $\mathbb{E}[(z+1)^{X_A(t)}|\alpha(wT)=k]$ is the generating function of $p_n^A(t)$ conditioned on $\alpha(wT)=k$. Similarly to the proof of Eqs. (12), it is easy to prove that

$$\mathbb{E}[(z+1)^{X_A(t)}|\alpha(wT)=k] = \sum_{j=0}^{1} L'_j(t-wT,z)F_j^A(wT,e^{-d(t-wT)}z|\alpha(wT)=k), \tag{29}$$

where $L'_j$, $i,j=0,1$ are functions obtained from $L_j$ by substituting the parameters $r$, $a$, $b$, and $u$ with the parameters $r'$, $a'$, $b'$, and $u'$, respectively,

$$F_0^A(wT,z|\alpha(wT)=0) = \frac{F_0(wT,z)}{F_0(wT,0)}, \quad F_1^A(wT,z|\alpha(wT)=0)=0,$$

and

$$F_0^A(0,z|\alpha(wT)=1)=0, \quad F_1^A(0,z|\alpha(wT)=1) = \frac{F_1(wT,z)}{F_1(wT,0)}.$$

Note that $\mathbb{E}[(z+1)^{X_B(t)}|\alpha(wT)=k]$ is the generating function of $p_n^B(t)$ conditioned on $\alpha(wT)=k$. Similarly to the proof of Eq. (29), we have

$$\mathbb{E}[(z+1)^{X_B(t)}|\alpha(wT)=k] = \sum_{j=0}^{1} L'_j(t-wT,z)F_j^B(wT,e^{-d(t-wT)}z|\alpha(wT)=k), \tag{30}$$

where

$$F_0^B(wT,z|\alpha(wT)=0)=1, \quad F_1^A(wT,z|\alpha(wT)=0)=0,$$

and

$$F_0^B(0,z|\alpha(wT)=1)=0, \quad F_1^A(0,z|\alpha(wT)=1)=1.$$

Inserting Eqs. (29) and (30) into Eq. (28), we finally obtain the generating function $F$ after replication, i.e.

$$F(t,z) = L'_0(t-wT,z)^2F_0(wT,e^{-d(t-wT)}z) + L'_1(t-wT,z)^2F_1(wT,e^{-d(t-wT)}z), \quad t\in[wT,T]. \tag{31}$$

In summary, we have derived the analytical expression of the generating function $F$ at any time $t\in[0,T]$ within a cell cycle, which is given by

$$F(t,z) = \begin{cases} \sum_{i=0}^{1} L_i(t,z)F_i(0,e^{-dt}z), & t\in[0,wT], \\ \sum_{i=0}^{1} L'_i(t-wT,z)^2F_i(wT,e^{-d(t-wT)}z), & t\in[wT,T], \end{cases} \tag{32}$$

where $F_i(wT,z)$, $i=0,1$ are determined by Eqs. (23) and (24). The time-dependent distribution of the mRNA number can be recovered by taking the derivatives of the generating function $F$ at $z=-1$, i.e.

$$p_n(t) = \frac{1}{n!}\frac{\partial^n}{\partial z^n}F(t,z)\Big|_{z=-1}. \tag{33}$$

#### 2.2 Derivation of the generating functions $F_0$ and $F_1$ after replication

We next compute the generating functions $F_i$, $i = 0, 1$ after replication. Recall that $F_i(t, z)$ is the generating function of

$$p_{i,n}(t) = \mathbb{P}(\alpha^A(t) = i,\ X^A(t) + X^B(t) = n),$$

which denotes the probability of having $n$ mRNA molecules in the cell at time $t$ when the daughter copy $A$ is in state $i$. In addition, recall that $F_i^A(t, z)$ is the generating function of

$$p_{i,n}^A(t) = \mathbb{P}(\alpha^A(t) = i,\ X^A(t) = n),$$

which denotes the probability of having $n$ mRNA molecules that belong to daughter copy $A$ at time $t$ when daughter copy $A$ is in state $i$. It is easy to see that the generating functions $F_i(t, z)$ and $F_i^A(t, z)$ can be represented by

$$F_i(t, z) = \mathbb{E}(z + 1)^{X_A(t)+X_B(t)} 1_{\{\alpha^A(t)=i\}}, \quad F_i^A(t, z) = \mathbb{E}(z + 1)^{X_A(t)} 1_{\{\alpha^A(t)=i\}},$$

where $1_A$ denotes the indicator function of the set $A$.

We now make a crucial observation that conditioned on $\alpha(wT) = j$, i.e. the gene state of the mother copy is $j$ at replication, the dynamics $\{\alpha^A(t), X^A(t)\}_{wT \le t \le T}$ for daughter copy $A$ and the dynamics $\{\alpha^B(t), X^B(t)\}_{wT \le t \le T}$ for daughter copy $B$ are independent of each other. This shows that

$$\begin{aligned} F_i(t, z) &= \mathbb{E}(z + 1)^{X_A(t)+X_B(t)} 1_{\{\alpha^A(t)=i\}} \\ &= \sum_{k=0}^{1} \mathbb{E}[(z + 1)^{X_A(t)+X_B(t)} 1_{\{\alpha^A(t)=i\}} | \alpha(wT) = k] \mathbb{P}(\alpha(wT) = k) \\ &= \sum_{k=0}^{1} \mathbb{E}[(z + 1)^{X_A(t)} 1_{\{\alpha^A(t)=i\}} | \alpha(wT) = k] \mathbb{E}[(z + 1)^{X_B(t)} | \alpha(wT) = k] F_k(wT, 0), \end{aligned} \tag{34}$$

where we have used the fact that $F_k(wT, 0) = \mathbb{P}(\alpha(wT) = k)$. Note that $\mathbb{E}[(z + 1)^{X_A(t)} 1_{\{\alpha^A(t)=i\}} | \alpha(wT) = k]$ is the generating function of $p_{i,n}^A(t)$ conditioned on $\alpha(wT) = k$. Similarly to the proof of Eqs. (23) and (24), it is easy to prove that

$$\mathbb{E}[(z + 1)^{X_A(t)} 1_{\{\alpha^A(t)=i\}} | \alpha(wT) = k] = \sum_{j=0}^{1} K'_{ij}(t - wT, z) F_j^A(wT, e^{-d(t-wT)} z | \alpha(wT) = k), \tag{35}$$

where $K'_{ij}$, $i, j = 0, 1$ are functions obtained from $K_{ij}$ by substituting the parameters $r$, $a$, $b$, $u$, and $v$ with the parameters $r'$, $a'$, $b'$, $u'$, and $v'$, respectively,

$$F_0^A(wT, z | \alpha(wT) = 0) = \frac{F_0(wT, z)}{F_0(wT, 0)}, \quad F_1^A(wT, z | \alpha(wT) = 0) = 0,$$

and

$$F_0^A(0, z | \alpha(wT) = 1) = 0, \quad F_1^A(0, z | \alpha(wT) = 1) = \frac{F_1(wT, z)}{F_1(wT, 0)}.$$

Inserting Eqs. (30) and (35) into Eq. (34), we obtain

$$\begin{aligned} F_i(t, z) = &K'_{i0}(t - wT, z) L'_0(t - wT, z) F_0(wT, e^{-d(t-wT)} z) \\ &+ K'_{i1}(t - wT, z) L'_1(t - wT, z) F_1(wT, e^{-d(t-wT)} z), \quad t \in [wT, T]. \end{aligned}$$

### 3 Modified FSP algorithm

**Input**:
1. All model parameters $V_b, g, \beta, w, d, \rho, \sigma_0, \sigma_1, \sigma_1'$
2. Truncation size $N = 8\rho/d_{\text{eff}}$
3. Initial distribution of the gene state and mRNA number

**Output**:
1. the time-dependent mRNA distribution across cell cycles
2. the steady-state mRNA distribution at birth
3. the steady-state mRNA distribution at replication
4. the steady-state mRNA distribution at division
5. the steady-state mRNA distribution for lineage measurements
6. the steady-state mRNA distribution for population measurements

**Algorithm**:
1. Set the initial Hellinger distance $D = 1$
2. Generate the initial distribution of the gene state and mRNA number
**while** $D > 10^{-4}$
 3. Solve the truncated CME before replication numerically using FSP
 4. Generate the distribution of the gene state and mRNA number for the two daughter copies at replication
 5. Solve the truncated CME after replication numerically using FSP
 6. Generate the distribution of the gene state and mRNA number for the mother copy at birth in the next generation based on binomial partitioning of molecules at division
 7. Compute the Hellinger distance $D$ between the mRNA distributions at birth in two successive generations
**end**
8. Record the steady-state mRNA distribution at birth
9. Solve the truncated CME before replication numerically using FSP
10. Record the steady-state mRNA distribution at replication
11. Compute $a_n^{\text{lin}} = \frac{1}{T}\int_0^{wT} p_n(t)dt$ for lineage measurements
12. Compute $a_n^{\text{pop}} = 2g\int_0^{wT} p_n(t)e^{-gt}dt$ for population measurements
13. Solve the truncated CME after replication numerically using FSP
14. Record the steady-state mRNA distribution at division
15. Compute $b_n^{\text{lin}} = \frac{1}{T}\int_{wT}^{T} p_n(t)dt$ for lineage measurements
16. Compute $b_n^{\text{pop}} = 2g\int_{wT}^{T} p_n(t)e^{-gt}dt$ for population measurements
17. Compute the steady-state mRNA distribution $p_n^{\text{lin}} = a_n^{\text{lin}} + b_n^{\text{lin}}$ for lineage measurements
18. Compute the steady-state mRNA distribution $p_n^{\text{pop}} = a_n^{\text{pop}} + b_n^{\text{pop}}$ for population measurements

### 4 Typical values of $\eta$ in various cell types

The following table (Table S1) shows the typical values of the doubling time $T = (\log 2)/g$, the mRNA half-life $T_H = \log(2)/d$, and the parameter $\eta = d/g = T/T_H$ in various cell types.

| Cell types | mean doubling time $T$ | median mRNA half-life $T_H$ | $\eta = T/T_H$ |
|---|---|---|---|
| *E. coli* (K10 + M9 + 30°C) | 73 min [2] | 2.1 min (0.6 - 24.6 min) [2] | 34.8 (3.0 - 121.7) |
| *E. coli* (DF261 + M9 + 30°C) | 78 min [2] | 2.8 min (0.8 - 43.5 min) [2] | 27.9 (1.8 - 97.5) |
| *E. coli* (N3433 + M9 + 30°C) | 75 min [2] | 3.7 min (1.0 - 44.2 min) [2] | 20.3 (1.7 - 75.0) |
| *E. coli* (SU02 + M9 + 30°C) | 75 min [2] | 4.0 min (1.6 - 55.5 min) [2] | 18.8 (1.4 - 46.9) |
| *E. coli* (YHC012 + M9 + 30°C) | 84 min [2] | 4.0 min (1.2 - 50.2 min) [2] | 21.0 (1.7 - 70.0) |
| *E. coli* (SH3208 + M9 + 30°C) | 81 min [2] | 3.2 min (1.1 - 24.6 min) [2] | 25.3 (3.3 - 73.6) |
| *E. coli* (BZ453 + M9 + 30°C) | 84 min [2] | 6.0 min (1.7 - 38.6 min) [2] | 14.0 (2.2 - 49.4) |
| *E. coli* (MG1655 + M9 + 30°C) | 30 min [3] | 6.8 min (0.7 - 34.5 min) [3] | 4.4 (0.9 - 42.9) |
| *E. coli* (NCM3416 + M9 + 30°C) | 90 min [4] | 5.4 min (1.3 - 31.4 min) [4] | 16.7 (2.9 - 69.2) |
| *E. coli* (NCM3416 + LB + 30°C) | 30 min [4] | 4.7 min (1.1 - 24.8 min) [4] | 6.4 (1.2 - 27.3) |
| *M. acetivorans* (WWM82 + MeOH + 37°C) | 7.5 h [5] | 1.0 h (4 min - 5.8 h) [5] | 7.5 (1.3 - 112.5) |
| *M. acetivorans* (WWM82 + TMA + 37°C) | 8.9 h [5] | 1.1 h (14 min - 8.3 h) [5] | 8.1 (1.1 - 38.1) |
| *M. acetivorans* (WWM82 + acetate + 37°C) | 24.6 h [5] | 2.8 h (11 min - 9.9 h) [5] | 8.8 (2.5 - 134.2) |
| *S. cerevisiae* | 1.5 - 2.5 h [6, 7] | 20 min (3 - $>$ 90 min) [8] | $4.5 - 7.5$ ($<$ 1.0 - 50.0) |
| *S. pombe* | 1.8 - 4.0 h [7, 9] | 33 min (10 - 96 min) [10] | $3.3 - 7.3$ (1.1 - 24.0) |
| Mouse Fibroblasts (N1H3T3) | 27.5 h [11] | 9 h (1.4 - 40 h) [11] | 3.1 (0.7 - 19.6) |
| Mouse embryonic stem cells | about 12 h [12] | 7.1 h (0.3 - $>$ 24 h) [13] | 1.7 ($<$ 0.5 - 40.0) |
| Human (HeLa) | 24.7 h [14] | 9.5 h (0.8 - 130 h) [15] | 2.6 (0.2 - 30.9) |
| Human cancer cells (HepG2) | 24 - 48 h [16] | 10 h (0.3 - 27.2 h) [16] | 3.6 (0.9 - 160.0) |

Table S1. **Biological values of the doubling time (cell-cycle duration), the mRNA half-life and the non-dimensional parameter $\eta$ for various cell types including bacteria, yeast and mammalian cells.** The bracket in the first column gives the strain, medium, and temperature for prokaryotes or the cell type for eukaryotes. The brackets in the last two columns give the ranges of the mRNA half-life for all studied genes in an experiment.

### 5 Further moment analysis for lineage and population measurements

Recall that the steady-state means of the mRNA number for lineage and population measurements are given by

$$\langle n\rangle_{\mathrm{lin}} = \frac{1}{T}\int_0^T \langle n\rangle^{ss}(t)dt, \quad \langle n\rangle_{\mathrm{pop}} = 2g\int_0^T \langle n\rangle^{ss}(t)e^{-gt}dt \tag{36}$$

If gene replication is not taken into account, i.e. $w = 1$, we have shown that the steady-state mRNA distribution at any time within a cell cycle is given by

$$\langle n\rangle^{ss}(t) = \lambda\left[e^{\beta gt} - \frac{2^{\eta+1}-2^{\eta+\beta}}{2^{\eta+1}-1}e^{-dt}\right]. \tag{37}$$

Inserting Eq. (37) into Eq. (36) gives the explicit expressions of the lineage and population means, i.e.

$$\langle n\rangle_{\mathrm{lin}} = \frac{\lambda}{\log 2}\left[\frac{2^\beta-1}{\beta} - \frac{(2-2^\beta)(2^\eta-1)}{\eta(2^{\eta+1}-1)}\right], \quad \langle n\rangle_{\mathrm{pop}} = \frac{\lambda(2-2^\beta)(\eta+\beta)}{(1-\beta)(\eta+1)}. \tag{38}$$

The explicit expression in the general case is too complicated and is omitted here.

We have seen that the mRNA means for the two types of measurements have different expressions. A natural question is how far the lineage statistics deviates from the population one. Fig. S2 (a),(b) show the the ratio of the lineage mean to the population mean, $R_1 = \langle n\rangle_{\mathrm{lin}}/\langle n\rangle_{\mathrm{pop}}$, as functions of $\beta$, $\eta$, $w$, and $\sigma_1'/\sigma_1$. Clearly, $R_1$ is always greater than 1, which means that the lineage mean is greater than the population mean [17]. As well, $R_1$ is largest when the mRNA synthesis rate scales with cell volume ($\beta = 1$), mRNA is unstable ($\eta \gg 1$), and there is no dosage compensation ($\sigma_1' = \sigma_1$). When these three conditions are satisfied, it follows from Eq. (35) in the main text that

$$\langle n\rangle(t) = \begin{cases} \lambda e^{gt}, & t\in(0,wT], \\ 2^{1+w}\lambda e^{g(t-wT)}, & t\in(wT,T]. \end{cases}$$

In this case, the lineage and population means can be obtained exactly as

$$\langle n\rangle_{\rm lin} = \frac{\lambda}{\log 2}(3-2w),\quad \langle n\rangle_{\rm pop} = (2\log 2)\lambda(2-w),$$

and it is easy to see that $R_1$ attains its maximum of $R_1^{\max}\approx 1.1$ when $w\approx 0.6$ (Fig. S2(b)). In other words, the lineage mean can differ from the population mean by at most $10\%$.

Moreover, we have also compared the variances of the mRNA number for lineage and population measurements and the ratio $R_2$ of the lineage variance to the population variance is shown in Fig. S2(c),(d) as functions of $\beta$, $\eta$, $w$, and $\sigma_1'/\sigma_1$. Similarly, $R_2$ is also large when the mRNA synthesis rate scales with cell volume, mRNA is unstable, and there is no dosage compensation.

### 6 Comparison with the conventional telegraph model

Another important question is whether the conventional telegraph model without a cell cycle description can capture the dynamic properties of the detailed telegraph model with such a description. Note that this is impossible when gene replication is taken into account since the mRNA distribution for the former has at most two modes, while the latter can have more than two modes. Hence, in the following, we only focus on the case when gene replication is not taken into account, i.e. $w=1$ [18].

Recall that the conventional telegraph model (with no volume-dependent rates) is characterized by the effective reactions

$$G \xrightarrow{\sigma_1} G^*,\quad G^* \xrightarrow{\sigma_0} G,\quad G^* \xrightarrow{\bar{\rho}} G^*+M,\quad M \xrightarrow{d_{\rm eff}} \varnothing,$$

where $d_{\rm eff}=d+g$ is the effective decay rate of mRNA. To make a fair comparison of the two models, the mRNA synthesis rate $\bar{\rho}$ of the conventional model is taken to be the time-average of the detailed model within a cell cycle, i.e.

$$\bar{\rho} = \frac{1}{T}\int_0^T \rho V(t)^\beta dt = \frac{(2^\beta-1)\rho V_b^\beta}{(\log 2)\beta}.$$

It is well known [19] that the steady-state mRNA mean for the conventional model is given by

$$\langle n\rangle_{\rm conv} = \frac{\sigma_1}{\sigma_0+\sigma_1}\cdot\frac{\bar{\rho}}{d_{\rm eff}} = \frac{(2^\beta-1)(\eta+\beta)\lambda}{(\log 2)\beta(\eta+1)}, \tag{39}$$

where $\lambda=au/b$ and $\eta=d/g$. Interestingly, when the mRNA synthesis rate is proportional to cell volume ($\beta=1$), we have $\langle n\rangle_{\rm conv}=\lambda/(\log 2)$, which agrees with the lineage mean given in Eq. (38). When the mRNA synthesis rate is independent of cell volume ($\beta=0$), we have $\langle n\rangle_{\rm conv}=\eta\lambda/(\eta+1)$, which coincides with the population mean given in (38). In addition, it follows from Eqs. (38) and (39) that the means for the conventional and detailed models are related by

$$\langle n\rangle_{\rm conv} = \frac{(2^\beta-1)(1-\beta)}{(\log 2)\beta(2-2^\beta)}\langle n\rangle_{\rm pop}.$$

This shows that the mean ratio $R'=\langle n\rangle_{\rm conv}/\langle n\rangle_{\rm pop}$ for the two models only depends on $\beta$ (Fig. S3(a)). It attains its minimum $R'_{\rm min}=1$ when $\beta=0$ and attains its maximum $R'_{\rm max}=1/2(\log 2)^2\approx 1.04$ when $\beta=1$. As a result, the conventional model can accurately capture the mRNA mean of the detailed model.

While the conventional model can capture the mean of the detailed model, it cannot accurately capture the mRNA distribution. To see this, recall that the steady-state distribution $p_n^{\rm conv}$ of the mRNA number for the conventional telegraph model has the generating function [19]

$$F_{\rm conv}(z) = \sum_{n=0}^\infty p_n^{\rm conv}(z+1)^n = M(\bar{a};\bar{b};\bar{u}z), \tag{40}$$

where $\bar{a} = \sigma_1/d_{\text{eff}}$, $\bar{b} = (\sigma_0 + \sigma_1)/d_{\text{eff}}$, and $\bar{u} = \bar{\rho}/d_{\text{eff}}$. Fig. S3(b) illustrates the Hellinger distance between the distributions for the two models as a function of $\beta$ and $\eta$. It can be seen that they coincide with each other when the transcription rate is volume-independent ($\beta = 0$) and mRNA is unstable ($\eta \gg 1$). In other cases, they deviate from each other significantly — the conventional model has a much smaller gene expression noise than the detailed model (Fig. S3(c)). This can be explained as follows. When $w = 1$, the steady-state inactive probability of the gene at birth is $p_{\text{off}}^b = (b-a)/b$ and the active probability of the gene at birth is $p_{\text{on}}^b = a/b$. Hence for unstable mRNAs, the time-dependent generating function can be simplified significantly as

$$F^{ss}(t,z) = \frac{b-a}{b}L_0(t,z) + \frac{a}{b}L_1(t,z) = M(a;b;ue^{\beta g t}z), \quad t \in (0,T]. \tag{41}$$

In particular, when $\beta = 0$ and $\eta \gg 1$, the steady-state mRNA distribution at any time within a cell cycle is independent of time $t$. Hence the generating functions of the lineage and population distributions are the same and are given by

$$F_{\text{lin}}(z) = F_{\text{pop}}(z) = M(a;b;uz), \tag{42}$$

where $a = \sigma_1/d$, $b = (\sigma_0+\sigma_1)/d$, and $u = \rho/d$. When $\beta = 0$ and $\eta \gg 1$, we have $\bar{\rho} = \rho$ and $d/d_{\text{eff}} \approx 1$. In this case, the two generating functions given in Eqs. (40) and (42) are approximately equal and thus the conventional model makes the correct prediction. We emphasize that there have been numerous studies that estimated the rate constants of stochastic gene expression dynamics based on the conventional telegraph model [20, 21]. Our results suggest that parameters estimated using this approach maybe unreliable.

While the conventional model fails to capture the mRNA distribution of the detailed model, we find that that it is capable of capturing the modality (unimodality or bimodality) of the distribution. To see this, following [22, 23], we define the strength of bimodality as

$$\kappa = \frac{H_{\text{low}} - H_{\text{valley}}}{H_{\text{high}}},$$

where $H_{\text{low}}$ is the height of the lower peak, $H_{\text{high}}$ is the height of the higher peak, and $H_{\text{valley}}$ is the height of the valley between them. For unimodal distributions, $\kappa$ is automatically set to be 0. For bimodal distributions, $\kappa$ is a quantity between 0 and 1 since $H_{\text{valley}} < H_{\text{low}} \leq H_{\text{high}}$. In general, to display strong bimodality, the following two conditions are necessary: (i) the two peaks should have similar heights and (ii) there should be a deep valley between them. The former ensures that the time periods spent in the low and high expression states are comparable, while the latter guarantees that the two expression levels are distinguishable. Clearly, $\kappa$ is large if both conditions are satisfied and is small if any one of the two conditions is violated. Hence, $\kappa$ serves as an effective indicator that characterizes the strength of bimodality.

Fig. S3(d),(e) illustrate $\kappa$ as a function of the gene switching rates $\sigma_0$ and $\sigma_1$ for the two models. Clearly, both models display unimodality in the regime of fast gene switching and display bimodality in the slow switching regime. Furthermore, we find that the regions in parameter space where the two models show bimodality are very close to each other, except that the detailed model needs a larger gene activation rate $\sigma_1$ to obtain the same strength of bimodality (shown by the blue and orange dashed lines in Fig. S3(d),(e)). This indicates the two models in general show the same modality but the heights of the modes may be different.

### 7 Generalization of the theory to stochastic cell volume dynamics

#### 7.1 Estimation of $\sigma_\epsilon$ in naturally occurring systems

To determine the value of $\sigma_\epsilon$ in real systems, we examined the lineage data collected in *E. coli* and haploid fission yeast cells using a mother machine [24, 25]. The *E. coli* data set contains the lineage measurements of cell length at three different temperatures (25°C, 27°C, and 37°C). The fission yeast data set contains the lineage measurements of cell area under seven different growth conditions (Edinburgh minimal medium (EMM) at 28°C, 30°C, 32°C, and 34°C and yeast extract medium (YE) at 28°C, 30°C, and 34°C).

The inference of $\sigma_\epsilon$ can be divided into the following three steps. First, the mean of the birth volume $V_b$ across all generations gives an estimate of $\bar{v}$. Next, the slope of the regression line of the division volume $V_d$ on the birth volume $V_b$ gives an estimate of $\alpha$. Finally, since $\alpha$ and $\bar{v}$ have been determined, $\sigma_\epsilon$ can be estimated as the sample standard deviation of $\epsilon = V_d - \alpha V_b - (2-\alpha)\bar{v}$. The estimated values of $\sigma_\epsilon$ in *E. coli* and fission yeast under all growth conditions are listed in Table S2.

| *E. coli* | | | |
|---|---|---|---|
| condition | 25°C | 27°C | 37°C |
| $\sigma_\epsilon$ | 0.3850 | 0.2401 | 0.2146 |

| fission yeast | | | | | | | |
|---|---|---|---|---|---|---|---|
| condition | EMM 28°C | EMM 30°C | EMM 32°C | EMM 34°C | YE 28°C | YE 30°C | YE 34°C |
| $\sigma_\epsilon$ | 0.2767 | 0.2518 | 0.2609 | 0.2593 | 0.2342 | 0.2093 | 0.2160 |

Table S2. **Estimate of $\sigma_\epsilon$ in *E. coli* and fission yeast under different growth conditions.**

#### 7.2 Analytical distributions for the model with stochastic cell volume dynamics

Given the values of the four variables $\alpha_b$, $N_b$, $V_b$, and $T$, the generating function $F$ at any time $t \in [0,T]$ within a cell cycle is given by Eqs. (12) and (31), i.e.

$$F(t,z|\alpha_b,N_b,V_b,T) = \begin{cases} \sum_{i=0}^{1} L_i(t,z|V_b)F_i(0,e^{-dt}z|\alpha_b,N_b), & t\in[0,wT], \\ \sum_{i=0}^{1} L_i'(t-wT,z|V_b)^2F_i(wT,e^{-d(t-wT)}z|\alpha_b,N_b,V_b,T), & t\in[wT,T]. \end{cases} \tag{43}$$

Here the initial conditions $F_i(0,z)$, $i=0,1$ are determined by $\alpha_b$ and $N_b$ as

$$F_{\alpha_b}(0,z|\alpha_b,N_b) = z^{N_b}, \quad F_{1-\alpha_b}(0,z|\alpha_b,N_b) = 0. \tag{44}$$

Let $\Pi^{(k)}(i,n,x,\tau) = \mathbb{P}^{(k)}(\alpha_b = i, N_b = n, V_b = x, T = \tau)$ denote the joint distribution of the four variables in generation $k$. Then the generating function $F$ at any fixed proportion $\theta \in [0,1]$ of the cell cycle in that generation is given by

$$\sum_{i=0}^{1}\sum_{n=0}^{\infty}\int_0^\infty\int_0^\infty F(\theta\tau,z|\alpha_b=i,N_b=n,V_b=x,T=\tau)\Pi^{(k)}(i,n,x,\tau)dxd\tau. \tag{45}$$

We next focus on the mRNA distribution in generation $k+1$. Similarly, once the values of the four variables are fixed, the generating functions $F_i$, $i=0,1$ at division are given by Eq. (14) in the main text, i.e.

$$F_i(T,z|\alpha_b,N_b,V_b,T) = \sum_{j=0}^{1}\tilde{K}_{ij}(z|V_b,T)F_j(0,e^{-dT}z|\alpha_b,N_b), \tag{46}$$

where the initial conditions $F_i(0,z)$, $i=0,1$ are determined by Eq. (44). To proceed, let $\alpha_d$ denote the state of daughter copy $A$ at division, let $N_d$ denote the mRNA number at division, and let $V_d$ denote the cell volume at division. From Eq. (46), we know the conditional joint distribution of $\alpha_d$ and $N_d$, i.e.

$$\mathbb{P}(\alpha_d = j, N_d = m|\alpha_b = i, N_b = n, V_b = x, T = \tau).$$

Hence the joint distribution of $\alpha_d$, $N_d$, $V_b$, and $T$ in generation $k$ is given by

$$\begin{aligned} &\mathbb{P}^{(k)}(\alpha_d = j, N_d = m, V_b = x, T = \tau) \\ &= \sum_{i=0}^{1}\sum_{n=0}^{\infty}\mathbb{P}(\alpha_d = j, N_d = m|\alpha_b = i, N_b = n, V_b = x, T = \tau)\Pi^{(k)}(i,n,x,\tau). \end{aligned}$$

From this it follows that the joint distribution of $\alpha_d$, $N_d$, and $V_d$ in generation $k$ is given by

$$\begin{aligned}\mathbb{P}^{(k)}(\alpha_d = j, N_d = m, V_d = y) &= \int_0^\infty \mathbb{P}^{(k)}(\alpha_d = j, N_d = m, V_d = y, T = \tau)d\tau,\\ &= \int_0^\infty \mathbb{P}^{(k)}(\alpha_d = j, N_d = m, V_b = e^{-g\tau}y, T = \tau)e^{-g\tau}d\tau.\end{aligned}$$

Since we have assumed binomial partitioning of molecules at division, the joint distribution of $\alpha_b$, $N_b$, and $V_b$ in generation $k+1$ is given by

$$\begin{aligned}\mathbb{P}^{(k+1)}(\alpha_b = i, N_b = n, V_b = x) &= 2\mathbb{P}^{(k)}(\alpha_d = i, N_d = m, V_d = 2x)\binom{m}{n}\left(\frac{1}{2}\right)^m\\ &= \mathbb{P}^{(k)}(\alpha_d = i, N_d = m, V_d = 2x)\binom{m}{n}\left(\frac{1}{2}\right)^{m-1}.\end{aligned}$$

Hence the joint distribution of the four variables $\alpha_b$, $N_b$, $V_b$, and $T$ in generation $k+1$ is given by

$$\Pi^{(k+1)}(i,n,x,\tau) = \mathbb{P}^{(k+1)}(\alpha_b = i, N_b = n, V_b = x)\mathbb{P}(T = \tau|V_b = x), \tag{47}$$

where the conditional distribution of $T$ given $V_b$ can be computed from Eq. (43) in the main text as

$$\mathbb{P}(T = \tau|V_b = x) = \frac{gxe^{g\tau}}{\sqrt{2\pi\sigma_\epsilon^2}}\exp\left[-\frac{(xe^{g\tau} - \alpha x - (2-\alpha)\bar{v})^2}{2\sigma_\epsilon^2}\right]. \tag{48}$$

Applying Eq. (45) and (47) repeatedly, we can compute the exact mRNA distribution at any time within a cell cycle in all generations. Finally, the steady-state joint distribution of the four variables $\alpha_b$, $N_b$, $V_b$, and $T$ is given by

$$\Pi^{ss}(i,n,x,\tau) = \lim_{k\to\infty}\Pi^{(k)}(i,n,x,\tau).$$

Inserting this equation into Eq. (45) gives the transient mRNA distribution under cyclo-stationary conditions.

Once the joint distribution of the four variables is known, it follows from Eq. (45) that the steady-state mRNA distribution for lineage measurements is given by

$$F^{\text{lin}}(z) = \frac{1}{\langle T\rangle}\sum_{i=0}^{1}\sum_{n=0}^{\infty}\int_0^\infty\int_0^\infty\left[\int_0^\tau F(t,z|\alpha_b = i, N_b = n, V_b = x, T = \tau)dt\right]\Pi^{ss}(i,n,x,\tau)dxd\tau, \tag{49}$$

where $\langle T\rangle \approx \log(2)/g$ is the mean doubling time. The exact expression for the steady-state population distribution is difficult to obtain. However, when the variability is cell cycle duration is small ($\sigma_\epsilon \ll 1$), an approximation of the steady-state population distribution is given by

$$F^{\text{pop}}(z) = 2g\sum_{i=0}^{1}\sum_{n=0}^{\infty}\int_0^\infty\int_0^\infty\left[\int_0^\tau F(t,z|\alpha_b = i, N_b = n, V_b = x, T = \tau)e^{-gt}dt\right]\Pi^{ss}(i,n,x,\tau)dxd\tau.$$

Note that when the cell volume dynamics is stochastic, the analytical mRNA distributions involve multiple integration, which is difficult to compute using the conventional multi-grid method. An alternative strategy to compute the multiple integration is to use the Monte Carlo method with the joint distribution $\Pi^{ss}(i,n,x,\tau)$ being randomly sampled.

#### 7.3 ENM approximations for stochastic cell volume dynamics

Recall that before replication, there is only one gene copy in the cell and thus the EDM is given by

$$G \xrightarrow{\sigma_1} G^*,\quad G^* \xrightarrow{\sigma_0} G,\quad G^* \xrightarrow{\rho V^\beta} G^* + M,\quad M \xrightarrow{d_{\text{eff}}} \varnothing. \tag{50}$$

After replication, there are two gene copies in the cell and thus the EDM should be modified as

$$\begin{aligned} & G_A \xrightarrow{\sigma_1'} G_A^*, \quad G_A^* \xrightarrow{\sigma_0} G_A, \quad G_B \xrightarrow{\sigma_1'} G_B^*, \quad G_B^* \xrightarrow{\sigma_0} G_B, \\ & G_A^* \xrightarrow{\rho V^\beta} G_A^* + M, \quad G_B^* \xrightarrow{\rho V^\beta} G_B^* + M, \quad M \xrightarrow{d_{\mathrm{eff}}} \varnothing, \end{aligned} \tag{51}$$

At cell birth, there is only one gene copy and thus the ENM approximation of the mRNA distribution is given by

$$p_{\mathrm{birth}}^{\mathrm{ENM}}(n) = \int_0^\infty p^{\mathrm{EDM}}(n|x)\mathbb{P}(V_b = x)dx,$$

where $p^{\mathrm{EDM}}(n|V)$ is the steady-state distribution of the EDM given in Eq. (50) and

$$\mathbb{P}(V_b = x) = \sqrt{\frac{4-\alpha}{2\pi\sigma_\epsilon^2}} e^{-\frac{4-\alpha}{2\sigma_\epsilon^2}(x-\bar{v})^2} \tag{52}$$

is the cell volume distribution at birth which is Gaussian [26]. Similarly, we can construct the ENM approximations for the mRNA distributions at replication and division.

We next construct the ENM approximation for the mRNA distribution of lineage measurements. This is much more complicated because for a given cell of volume $V$, it is unclear whether it has one or two gene copies. To determine the number of gene copies in the cell, we also need the information of the birth volume $V_b$ and the cell cycle duration $T$. Once the values of $V$, $V_b$, and $T$ are known, the age of the cell is $t = (1/g)\log(V/V_b)$. It has only one gene copy when $t < wT$ and has two gene copies when $t \geq wT$. Hence the ENM approximation for the lineage distribution is given by

$$p_{\mathrm{lin}}^{\mathrm{ENM}}(n) = \frac{1}{\langle T \rangle} \int_0^\infty \int_0^\infty \left[ \int_0^\tau p^{\mathrm{EDM}}(n|t,x,\tau)dt \right] \mathbb{P}(V_b = x, T = \tau)dxd\tau.$$

Here $p^{\mathrm{EDM}}(n|t,x,\tau)$ is steady-state distribution of the EDM for a cell of age $t$ given that the birth volume is $x$ and the cell cycle duration is $\tau$, and

$$\begin{aligned} \mathbb{P}(V_b = x, T = \tau) &= \mathbb{P}(V_b = x)\mathbb{P}(T = \tau | V_b = x) \\ &= \frac{\sqrt{4-\alpha}}{2\pi\sigma_\epsilon^2} gxe^{g\tau} \exp\left[ -\frac{(xe^{g\tau} - \alpha x - (2-\alpha)\bar{v})^2 + (4-\alpha)(x-\bar{v})^2}{2\sigma_\epsilon^2} \right] \end{aligned}$$

is the joint distribution of $V_b$ and $T$. Note that the reaction scheme given in Eq. (50) should be used for the EDM when $t < w\tau$ and the reaction scheme given in Eq. (51) should be used when $t \geq w\tau$.

### References

[1] Iyer-Biswas, S., Hayot, F. & Jayaprakash, C. Stochasticity of gene products from transcriptional pulsing. *Phys. Rev. E* **79**, 031911 (2009).

[2] Bernstein, J. A., Lin, P.-H., Cohen, S. N. & Lin-Chao, S. Global analysis of Escherichia coli RNA degradosome function using DNA microarrays. *Proc. Natl. Acad. Sci. USA* **101**, 2758–2763 (2004).

[3] Selinger, D. W., Saxena, R. M., Cheung, K. J., Church, G. M. & Rosenow, C. Global RNA half-life analysis in Escherichia coli reveals positional patterns of transcript degradation. *Genome Res.* **13**, 216–223 (2003).

[4] Bernstein, J. A., Khodursky, A. B., Lin, P.-H., Lin-Chao, S. & Cohen, S. N. Global analysis of mRNA decay and abundance in Escherichia coli at single-gene resolution using two-color fluorescent DNA microarrays. *Proc. Natl. Acad. Sci. USA* **99**, 9697–9702 (2002).

[5] Peterson, J. R. *et al.* Genome-wide gene expression and RNA half-life measurements allow predictions of regulation and metabolic behavior in Methanosarcina acetivorans. *BMC Genomics* **17**, 924 (2016).

[6] Khmelinskii, A. *et al.* Tandem fluorescent protein timers for in vivo analysis of protein dynamics. *Nat. Biotechnol.* **30**, 708 (2012).

[7] Christiano, R., Nagaraj, N., Fröhlich, F. & Walther, T. C. Global proteome turnover analyses of the yeasts S. cerevisiae and S. pombe. *Cell Rep.* **9**, 1959–1965 (2014).

[8] Wang, Y. *et al.* Precision and functional specificity in mRNA decay. *Proc. Natl. Acad. Sci. USA* **99**, 5860–5865 (2002).

[9] Siegal-Gaskins, D. & Crosson, S. Tightly regulated and heritable division control in single bacterial cells. *Biophys. J.* **95**, 2063–2072 (2008).

[10] Lackner, D. H. *et al.* A network of multiple regulatory layers shapes gene expression in fission yeast. *Mol. Cell* **26**, 145–155 (2007).

[11] Schwanhäusser, B. *et al.* Global quantification of mammalian gene expression control. *Nature* **473**, 337 (2011).

[12] Festuccia, N., Gonzalez, I. & Navarro, P. The epigenetic paradox of pluripotent ES cells. *J. Mol. Biol.* **429**, 1476–1503 (2017).

[13] Sharova, L. V. *et al.* Database for mRNA half-life of 19 977 genes obtained by DNA microarray analysis of pluripotent and differentiating mouse embryonic stem cells. *DNA Res.* **16**, 45–58 (2009).

[14] Boisvert, F.-M. *et al.* A quantitative spatial proteomics analysis of proteome turnover in human cells. *Mol. Cell Proteomics* **11** (2012).

[15] Lim, J. *et al.* Uridylation by TUT4 and TUT7 marks mRNA for degradation. *Cell* **159**, 1365–1376 (2014).

[16] Yang, E. *et al.* Decay rates of human mRNAs: correlation with functional characteristics and sequence attributes. *Genome Res.* **13**, 1863–1872 (2003).

[17] Beentjes, C. H., Perez-Carrasco, R. & Grima, R. Exact solution of stochastic gene expression models with bursting, cell cycle and replication dynamics. *Phys. Rev. E* **101**, 032403 (2020).

[18] Thomas, P. & Shahrezaei, V. Coordination of gene expression noise with cell size: extrinsic noise versus agent-based models of growing cell populations. *J. R. Soc. Interface* **18**, 20210274 (2021).

[19] Peccoud, J. & Ycart, B. Markovian modeling of gene-product synthesis. *Theor. Popul. Biol.* **48**, 222–234 (1995).

[20] Suter, D. M. *et al.* Mammalian genes are transcribed with widely different bursting kinetics. *Science* **332**, 472–474 (2011).

[21] Larsson, A. J. *et al.* Genomic encoding of transcriptional burst kinetics. *Nature* **565**, 251–254 (2019).

[22] Jia, C. & Grima, R. Small protein number effects in stochastic models of autoregulated bursty gene expression. *J. Chem. Phys.* **152**, 084115 (2020).

[23] Jia, C. & Grima, R. Dynamical phase diagram of an auto-regulating gene in fast switching conditions. *J. Chem. Phys.* **152**, 174110 (2020).

[24] Tanouchi, Y. *et al.* A noisy linear map underlies oscillations in cell size and gene expression in bacteria. *Nature* **523**, 357–360 (2015).

[25] Nakaoka, H. & Wakamoto, Y. Aging, mortality, and the fast growth trade-off of Schizosaccharomyces pombe. *PLoS Biol.* **15**, e2001109 (2017).

[26] Amir, A. Cell size regulation in bacteria. *Phys. Rev. Lett.* **112**, 208102 (2014).

[27] Skinner, S. O. *et al.* Single-cell analysis of transcription kinetics across the cell cycle. *Elife* **5**, e12175 (2016).

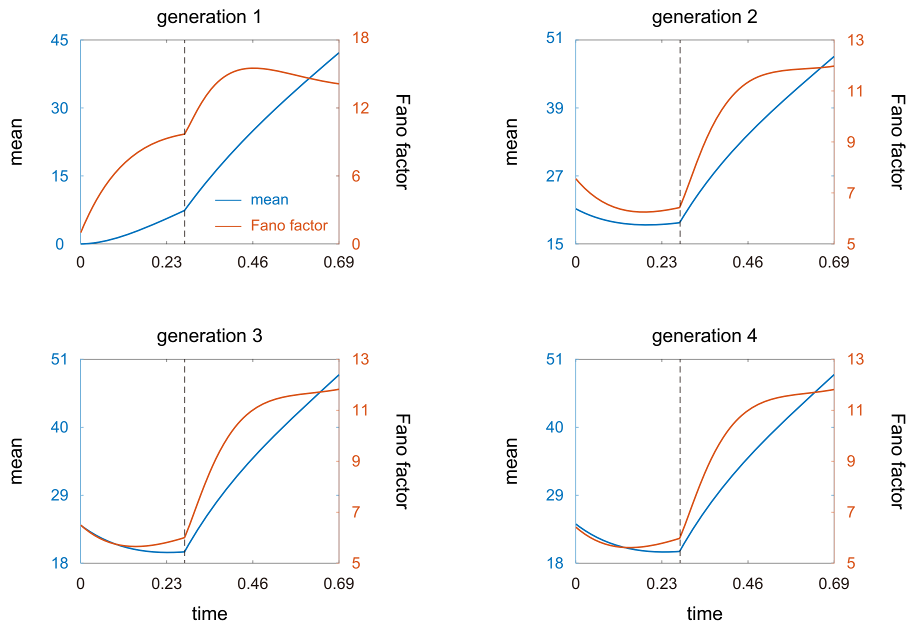

Figure S1. **Time-dependent mean and Fano factor of the mRNA number across four cell cycles.** Here we assume that initially there is no mRNA in the cell and the gene is inactive. The model parameters are chosen as $V_b = 1, g = 1, \beta = 1, w = 0.4, d = 5, \rho = 20d_{\mathrm{eff}}, \sigma_0 = 1.5, \sigma_1 = 3, \sigma_1' = 2.4$. Under these parameters, the cell cycle duration is given by $T = (\log 2)/g \approx 0.69$ and replication occurs at $t = wT \approx 0.28$, which is shown by a vertical dashed line.

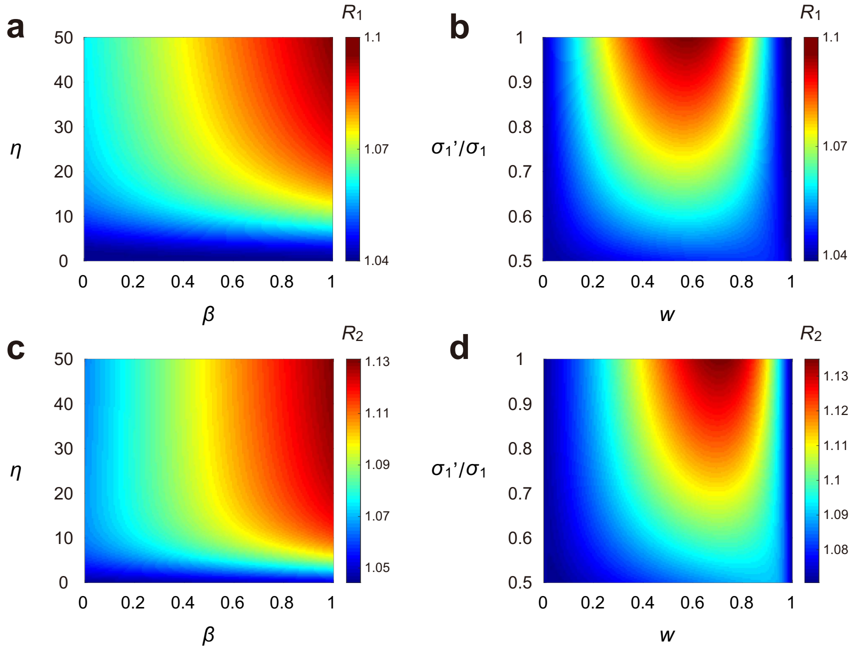

Figure S2. **Comparison between the moment statistics for lineage and population measurements.** (**a**) Heat plot of the ratio $R_1$ of the lineage mean to the population mean as $\beta$ and $\eta$ vary. (**b**) Heat plot of $R_1$ as $w$ and $\sigma_1'/\sigma_1$ vary. (**c**) Heat plot of the ratio $R_2$ of the lineage variance to the population variance as $\beta$ and $\eta$ vary. (**d**) Heat plot of $R_2$ as $w$ and $\sigma_1'/\sigma_1$ vary. In (a),(c), the model parameters are chosen as $V_b = 1, g = 1, w = 0.6, \rho = 10d_{\mathrm{eff}}, \sigma_0 = 100, \sigma_1 = \sigma_1' = 10$. In (b),(d), the model parameters are chosen as $V_b = 1, g = 1, \beta = 1, d = 50, \rho = 10d_{\mathrm{eff}}, \sigma_0 = 100, \sigma_1 = 10$.

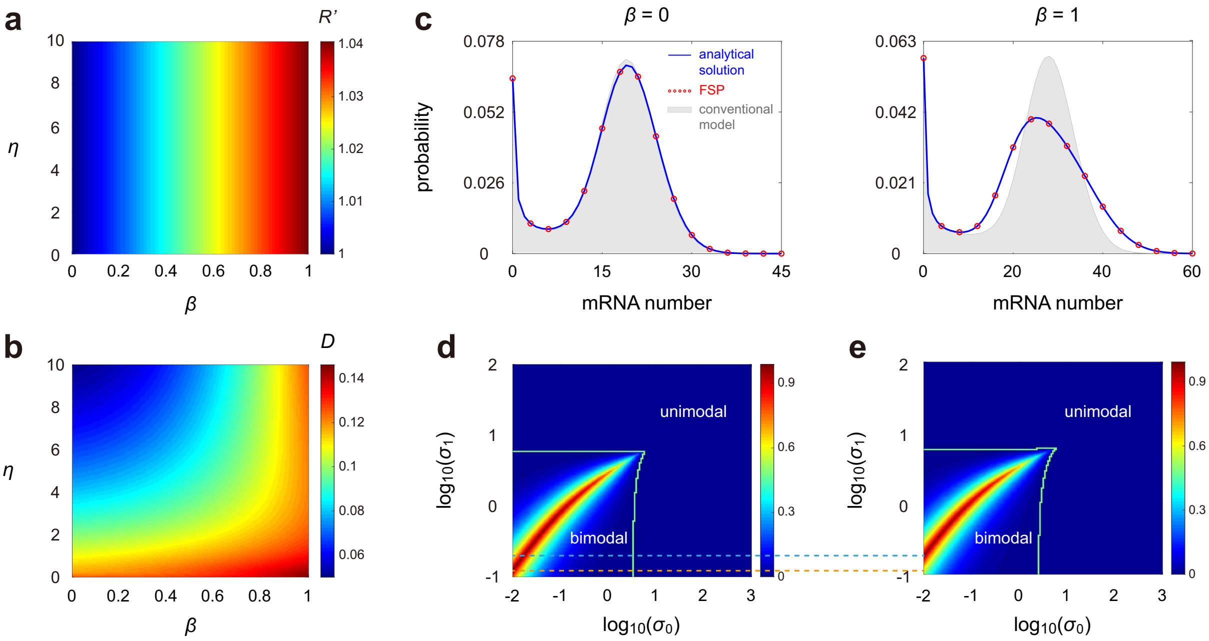

Figure S3. **Comparison between the conventional and detailed telegraph models when gene replication is not taken into account.** (**a**) Heat plot of the ratio $R'$ of the steady-state mean of the conventional model to the population mean of the full model as $\beta$ and $\eta$ vary. (**b**) Heat plot of the Hellinger distance $D$ between the steady-state distributions of the two models as $\beta$ and $\eta$ vary. Here the lineage distribution is used for the full model. (**c**) Comparison of the steady-state distributions of the two models for unstable mRNAs when $\beta = 0$ and $\beta = 1$. The blue curves and the red circles show the analytical and numerical distributions for the full model, and the grey regions show the distributions for the conventional model. In (a)-(c), the model parameters are chosen as $V_b = 1, g = 1, w = 1, d = \eta g, \rho = 20d_{\mathrm{eff}}, \sigma_0 = 3, \sigma_1 = 15$, and $\eta = 50$ in (c). (**d**) Heat plot of the strength $\kappa$ of bimodality of the steady-state distribution of the conventional model as $\sigma_0$ and $\sigma_1$ vary. The green curve encloses the region where the distribution exhibits bimodality. (**e**) Same as (c) but for the lineage distribution of the detailed model. In (d),(e), the model parameters are chosen as $V_b = 1, g = 1, \beta = 1, w = 1, d = 5, \rho = 20d_{\mathrm{eff}}$. The blue and orange dashed lines show the values of $\sigma_1$ at which bimodality for the two models is the strongest when $\sigma_0 = 0.01$.

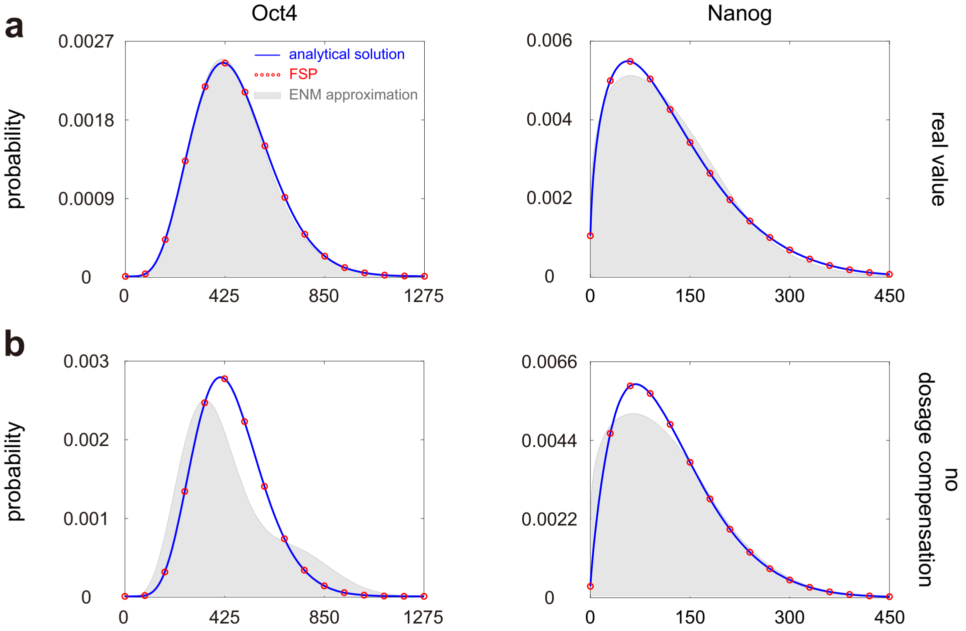

Figure S4. **Comparison between the mRNA distributions for the full model and the ENM under two sets of parameters obtained from experiments.** (**a**) The population mRNA distributions for Oct4 and Nanog genes in wild-type R1 mouse embryonic stem cells. Here we use the parameters estimated in [27]. For Oct4, the model parameters are given by $T = 13$ hr, $\beta = 0$, $w = 0.67$, $d = 0.0023$ min$^{-1}$, $\sigma_0 = 0.018$ min$^{-1}$, $\sigma_0 = 0.0092$ min$^{-1}$, $\sigma_1' = 0.63\sigma_0$. The parameter $\rho$ is determined so that $\langle n\rangle_{\text{pop}} = 477$. For Nanog, the model parameters are given by $T = 13$ hr, $\beta = 0$, $w = 0.49$, $d = 0.0022$ min$^{-1}$, $\sigma_0 = 0.007$ min$^{-1}$, $\sigma_0 = 0.0019$ min$^{-1}$, $\sigma_1' = 0.71\sigma_0$. The parameter $\rho$ is determined so that $\langle n\rangle_{\text{pop}} = 125$. The blue curves and the red circles show the analytical and numerical distributions for the full model, and the grey regions show the distributions of the ENM. An important property of the full model is that when the steady-state mean of the mRNA number is fixed, the steady-state distribution is actually independent of the birth volume $V_b$. Hence here we simply take $V_b = 1$. (**b**) Same as (a) but in the case of no dosage compensation ($\sigma_1' = \sigma_1$).

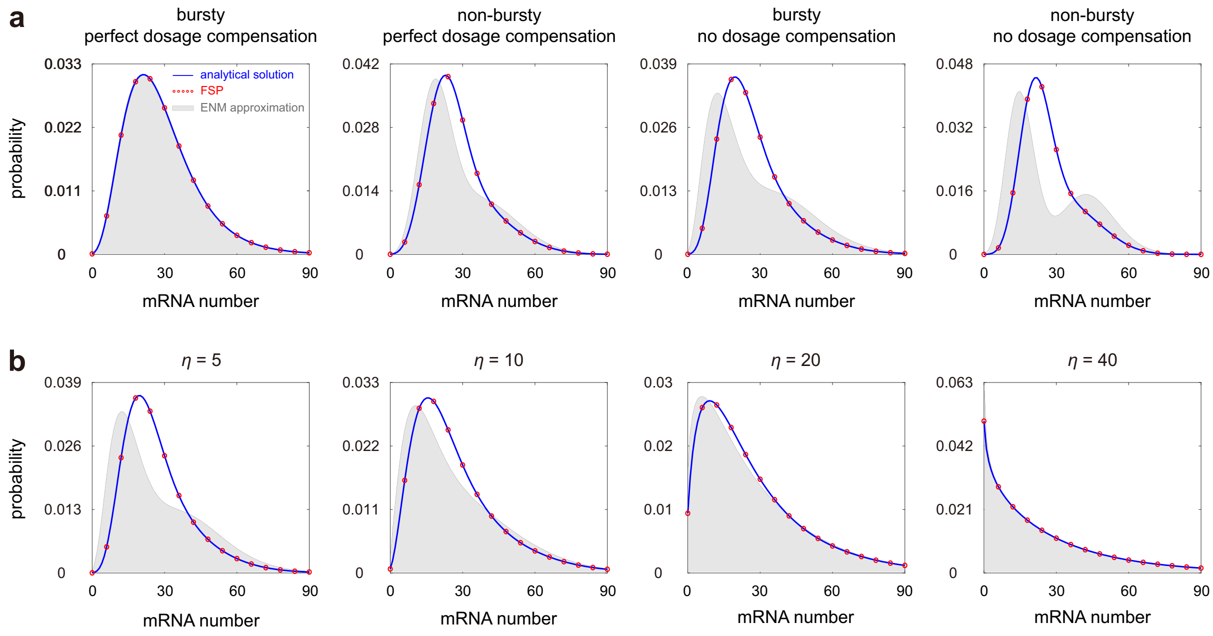

Figure S5. **Comparison between the full model and the ENM for population measurements.** (**a**) Comparison between the population distributions for the full model and the ENM as $\sigma_0/\sigma_1$ and $\sigma_1'/\sigma_1$ vary. The blue curves show the analytical distributions for the full model, the red circles show the numerical ones obtained from FSP, and the grey regions show the distributions for the ENM. The model parameters are chosen as $V_b = 1, g = 1, \beta = 1, w = 0.5, d = 5, \sigma_1 = 30$. The parameter $\sigma_0$ is chosen as $\sigma_0 = 10\sigma_1$ (bursty case) and $\sigma_0 = 0.5\sigma_1$ (non-bursty case). The parameter $\sigma_1'$ is chosen as $\sigma_1' = \sigma_1/2$ (perfect dosage compensation) and $\sigma_0 = \sigma_1$ (no dosage compensation). (**a**) The model parameters are chosen to be the same as in the third panel of (a) but $\eta$ is varied. In (a),(b) the parameter $\rho$ is chosen so that $\langle n\rangle_{\mathrm{lin}} = 30$.

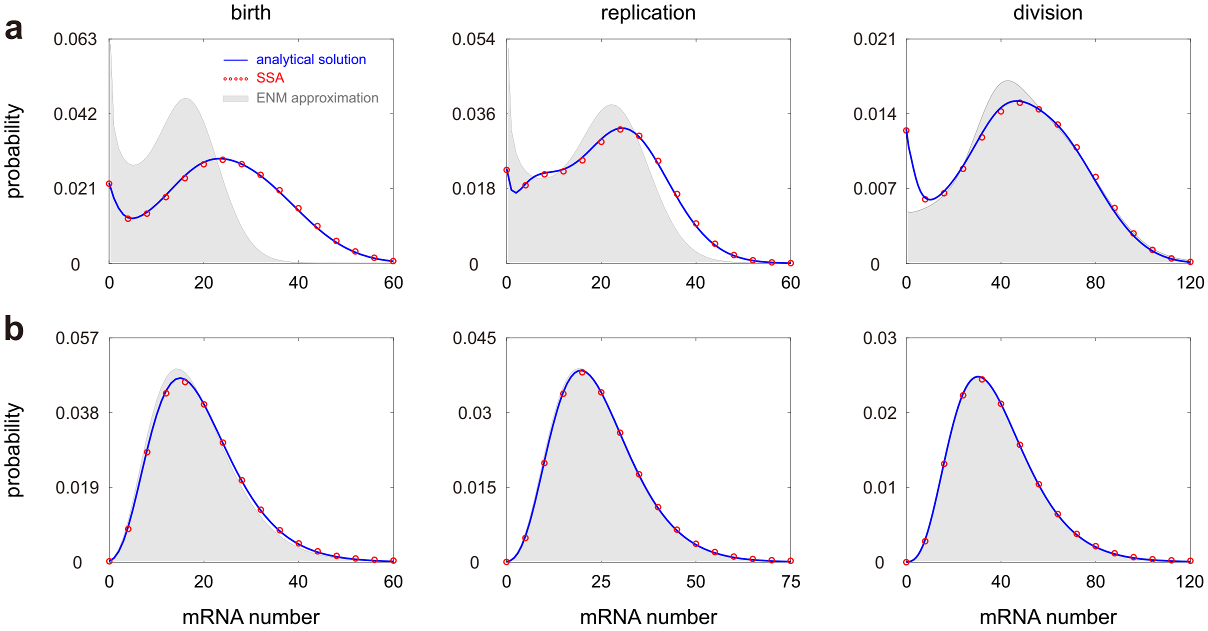

Figure S6. **Comparison between the time-dependent distributions of the full model and the ENM under cyclo-stationary conditions for stochastic cell volume dynamics.** (**a**) Steady-state distributions at birth, replication, and division for the full model and the ENM. The blue curves show the analytical distributions given in Eqs. (43) and (49), the red circles show the numerical ones obtained from the SSA, and the grey regions show the distributions of the ENM (see Section 8.3 for the computation of the latter). The model parameters are chosen as $\bar{v} = 1, g = 1, \beta = 1, w = 0.4, d = 4, \rho = 20d_{\text{eff}}, \sigma_0 = 1.5, \sigma_1 = 3, \sigma_1' = 2.4, \alpha = 1, \sigma_\epsilon = 0.3$. (**b**) Same as (a) but in the special case where mRNA synthesis is balanced and bursty and dosage compensation is perfect. The model parameters are chosen as $\bar{v} = 1, g = 1, \beta = 1, w = 0.4, d = 4, \rho = 200d_{\text{eff}}, \sigma_0 = 300, \sigma_1 = 30, \sigma_1' = 15, \alpha = 1, \sigma_\epsilon = 0.3$.
